# Elucidation of Sequence Constraints for Protein Folding on the Ribosome

**DOI:** 10.64898/2026.11.07.765108

**Authors:** Rohan Bhardwaj, Caden Sillman, Charles P. Kuntz, Brianna Corman, J. Paul Robinson, Jonathan P. Schlebach

## Abstract

The pioneering round of protein folding on the ribosome is critical for the proper assembly and function of many proteins. Emerging evidence suggests some proteins sample distinct conformational intermediates during translation, which implies the mechanisms of cotranslational folding may impose specific constraints on protein sequence. To assess how folding on ribosome may shape mutational tolerance, we paired a fluorogenic arrest peptide sensor with deep mutational scanning to survey how 9,480 mutations modulate *E. coli* dihydrofolate reductase (DHFR) folding on the ribosome in live cells. By controlling for differences in variant abundance, we assess how these mutations specifically impact the efficiency of DHFR folding on stalled ribosomes. Mutagenic trends are broadly correlated with computational estimates for the change in the free energy of the native fold, suggesting that the sequence constraints associated with late-stage folding on the ribosome are broadly similar to those of post-translational folding. However, a close inspection of mutagenic trends reveals that folding on the ribosome is vulnerable to perturbations of specific contacts within a central portion of the adenosine binding domain and the C-terminal β-strands of the discontinuous loop domain. Some of these regions are known to retain structure in the context of conformational intermediates that form on and off the ribosome, while others appear to be specific to cotranslational interactions. Together, our findings highlight how the constraints of folding on the ribosome may constrain evolutionary sequence space.

## Introduction

Protein evolution is guided by selective pressures that suppress the accumulation of sequence variants that reduce fitness- an aggregate descriptor for how well a genetic variant supports organismal viability. Though there are many ways that mutations modulate the net activity of a protein, which is typically correlated with fitness, their impacts on protein folding represent a key factor that shapes evolvability.^1–4^ Mutations that destabilize the native fold reduce the yield of protein biogenesis and create proteostatic hazards by promoting misfolding and premature degradation. The negative effects of these variants can also be magnified through epistatic interactions with other mutations that perturb conformational equilibria.^5–9^ While computational estimates of folding energetics of full-length proteins have provided a useful lens to rationalize protein evolution, mounting evidence suggest that the relevant folding transitions for many proteins must occur during translation before the native structure can be formed.^10,11^ Moreover, the genomic clustering of rare codons suggests these processes do indeed impose a selection pressure on certain genes.^12–14^ While cotranslational folding transitions are sure to differ from those involved in post-translational folding,^15,16^ it is unclear how the mechanistic basis of vectorial assembly may differentially constrain protein sequence.

It has been estimated that a third of the proteome relies on the pioneering round of cotranslational folding to achieve its native structure.^10^ During cotranslational folding, the ribosome effectively restricts the conformational search of the nascent chain in a manner that guides it through a series of conformational intermediates that may often diverge from those sampled by full-length proteins.^15–17^ As protein synthesis is generally slower than protein folding, the pioneering round of folding is typically initiated prior to the completion of translation.^18^ Thus, many nascent polypeptides undergo compaction, folding, cofactor binding, and begin to form quality control interactions in or near the ribosomal exit tunnel during synthesis.^19–21^ Pioneering applications of hydrogen-deuterium exchange (HDX),^15,22,23^ NMR spectroscopy,^24–26^ single-molecule force spectroscopy,^27^ and cryo-EM^16,20^ to cotranslational folding suggest this process can be understood using some of the guiding principles that shape the conformational energy landscapes of full-length proteins. And while there are indications that cotranslational processes impact fitness,^28^ few generalizable principles have emerged concerning the manner in which the distorted cotranslational energy landscapes may alter fitness effects. Additional insights into how mutations differentially impact co-translational folding and post-translational folding are needed.

Many recent efforts to explore cotranslational folding intermediates have leveraged translational arrest peptides from the *Escherichia coli* SecM protein to detect conformational transitions in the nascent chain that occur in the context of stalled ribosomes.^19,21,27,29–32^ These peptides form specific interactions within the ribosomal exit tunnel to induce a translational stall that can be reversed by mechanical tensions produced by folding transitions in the nascent chain.^33^ SecM readthrough efficiency can therefore serve as a detector for cotranslational folding on the ribosome. SecM sensors are typically scanned across the length of the protein of interest in order to detect the points in protein synthesis where the nascent chain begins to form structure. However, these experimental designs provide little insight into the nature of the contacts that form within these intermediates or how they may shape mutational tolerance. In this work, we combine an *in vivo* arrest peptide reporter with deep mutational scanning (DMS) to survey how folding on the ribosome may constrain the sequence of *Escherichia coli* dihydrofolate reductase (DHFR), a historically important model for protein folding studies. By normalizing the SecM read-through signal generated by stalled DHFR variants according to steady-state expression levels, we map the effects of 9,480 mutations on the folding of DHFR on the ribosome. Our results are broadly correlated with computational estimates of folding energetics, suggesting these mutations generally perturb native-like structural contacts. However, mutagenic trends suggest that the formation of native contacts within the central portion of the adenosine binding domain as well as in two C-terminal β-stands of the discontinuous loop domain are particularly critical for this folding transition. Together, our results provide evidence of how cotranslational folding may constrain protein sequence.

## Results

### An In vivo Sensor for DHFR Folding on the Ribosome

To probe DHFR folding on the ribosome within living cells, we developed a fluorescence reporter in which the *E. coli* DHFR open reading frame (ORF) is positioned upstream of a cassette encoding a C-terminal linker, a SecM arrest peptide, and a GFP reporter gene (Fig. 1A). This construct is designed such that the tension generated by the folding of the nascent DHFR domain promotes readthrough of the SecM-mediated stall in a manner that facilitates the translation of the downstream GFP ORF. We also included an independent red fluorescent protein (mKate) cassette as an internal expression control. The design, which is akin to that of a recently described reporter from the Kaiser group,^27^ facilitates quantitative comparisons of SecM readthrough (GFP: mKate ratio) in the context of living *E. coli*. Cellular GFP: mKate ratios for a negative control construct bearing a premature termination codon in the GFP cassette are 15.6-fold lower than those of cells expressing a positive control construct bearing a SecM mutation that prevents stalling (P166A). This observation confirms that cellular GFP: mKate ratios are limited by SecM readthrough.

**Figure 1.**
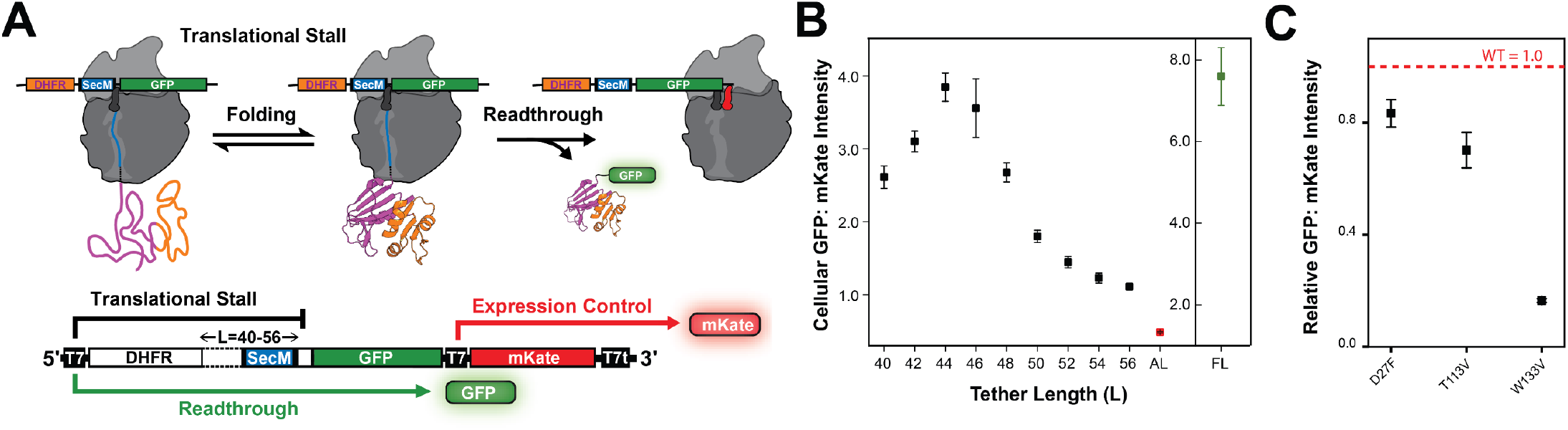
An *In vivo* Reporter for DHFR Folding on the Ribosome. A) A schematic depicts how the coupling between the folding of nascent DHFR and SecM read through promote the selective translation of GFP. A red fluorescent protein (mKate) produced from a secondary promoter serves as an expression control. B) Flow cytometry was used to compare the cellular GFP: mKate intensity ratio among *E. coli* expressing a series of reporter constructs in which the linker region separating the DHFR gene from the SecM sequence was varied from 40-56 codons. The signals from a negative control bearing premature termination codon (AL, L=46) and a positive control bearing a mutation that prevents stalling (FL, L=46) are shown for reference. C) A plot depicts the relative cellular GFP: mKate intensity ratios derived from *E. coli* expressing reporters (L=46) encoding three DHFR variants (D27F, T113V, and W133V). A dashed line indicates the signal of the corresponding WT DHFR reporter. Ratios represent the average from three biological replicates and whiskers reflect the standard deviation.

Previous investigations showed that the coupling between DHFR folding and SecM readthrough is maximized when there are roughly 40 residues separating the C-terminus of DHFR from the stall site.^21^ These residues reside within the exit tunnel and allow the nascent DHFR to clear the ribosome while the ribosome is stalled (Fig.1A). Consistent with these previous *in vitro* measurements,^21^ we find that cellular GFP: mKate intensity ratios are maximized at linker lengths of 44-46 residues (Fig. 1B). To confirm that SecM readthrough is driven by DHFR folding, we assessed the effects of a previously characterized DHFR-destabilizing mutation (W133V)^34,35^ and two stabilizing mutations (T113V and D27F)^36^ on reporter signal at a fixed linker length (L=46). The stabilizing mutations cause a modest effect on cellular GFP: mKate intensities (Fig. 1C). By comparison, the destabilizing mutation causes a large (84 ± 4%) decrease in the cellular GFP: mKate intensity ratio (Fig. 1C). These results show that mutations that suppress folding also decrease SecM readthrough. Together, these findings demonstrate that our *in vivo* reporter recapitulates observations for arrest peptide-based *in vitro* measurements of DHFR folding on the ribosome.

### Deep mutational scanning of stalled DHFR

To map the sequence constraints of folding on the ribosome, we carried out a deep mutational scan of the DHFR domain in the context of this SecM-based fluorogenic reporter. We first generated a genetic library in which each DHFR codon was individually randomized. We then integrated this pooled library of 9,480 DHFR variants into two reporter backbones-one where the library is fused to a native SecM-GFP and another in which the library is instead fused to a defective SecM-GFP (P166A). Comparing variant intensity ratios in the context of the native SecM backbone (stalled) to those in the defective SecM backbone (un-stalled) provides a means to account for differences in the solubility/ accumulation of mutated DHFR-GFP fusion products-a crucial consideration to ensure deviations in signal reflect differences in readthrough specifically. *E. coli* expressing these two libraries each exhibit a comparable range of mKate intensities, which suggests these constructs do not perturb plasmid copy number (Fig. 2A). Cells expressing the WT SecM library exhibit a broad distribution of GFP intensities (Fig. 2B). In contrast, this same collection of variants is bimodal with respect to GFP intensity upon mutagenic disruption of SecM (Fig. 2B), which suggests a subset of the mutated reporter variants are targeted for degradation. These wide variations in GFP create similar patterns in single-cell GFP: mKate intensity ratios (Fig. 2C), which tracks with changes in DHFR folding (Fig. 1C). To quantitatively compare the SecM readthrough induced by each variant, we first used fluorescence activated cell sorting (FACS) to fractionate cells expressing each version of the library (stalled and un-stalled) according to cellular GFP: mKate intensity ratios. We then used deep sequencing to track the relative abundance of each variant within each cellular fraction which is then used to estimate the GFP: mKate intensity ratio for each variant as previously described.^37^

**Figure 2.**
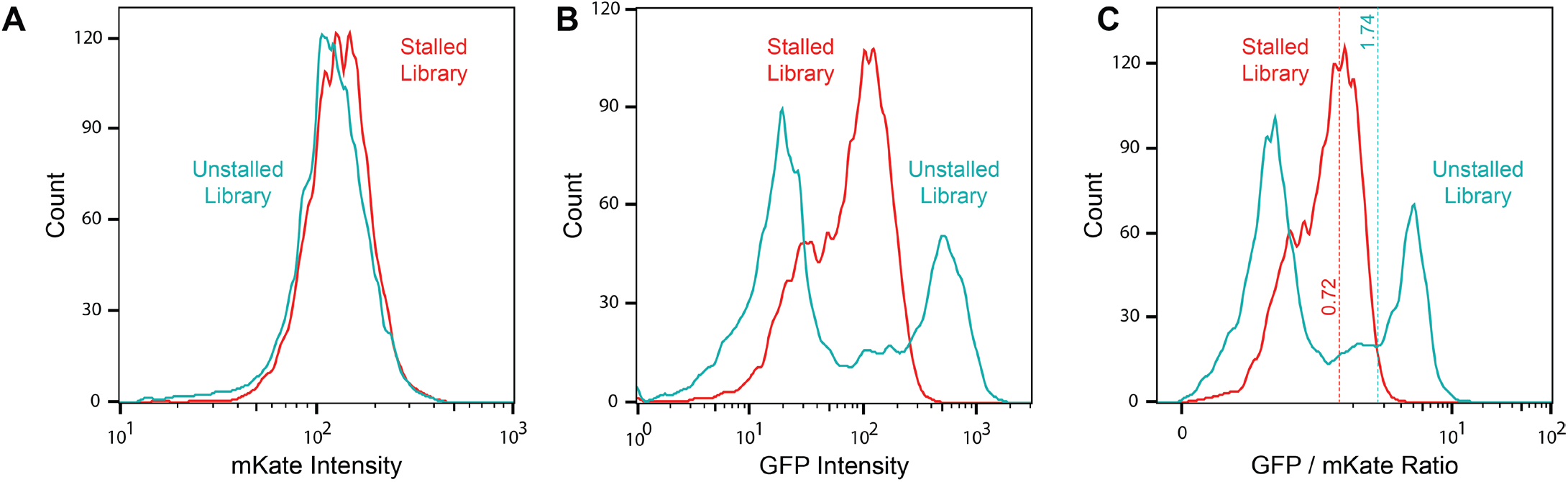
Expression of a Pooled DHFR Sensor Library. Flow cytometry was used to survey the distribution of cellular fluorescence intensities among *E. coli* expressing a pooled library of DHFR variants in the context of the stalled SecM-GFP sensor (red) or un-stalled P166A SecM-GFP sensor (blue) constructs. A) A histogram compares the distributions of the cellular fluorescence intensity associated with the constitutively expressed mKate protein generated by each version of the library. B) A histogram compares the distributions of the cellular fluorescence intensity associated with the GFP protein generated by each version of the library. C) A histogram compares the distributions of the cellular GFP: mKate intensity ratios generated by each version of the library. The dashed lines represent the GFP: mKate ratio of wild-type DHFR in SecM and P166A-SecM reporters. Measurements are derived from one representative biological replicate.

Overall, our results confidently determined the effects of 9,480 DHFR variants in the context of the stalled and non-stalled versions of the sensor (Table S1). Mutagenic trends appear to be generally similar regardless of whether the SecM sequence induces a translational stall (Spearman’s σ = 0.43, *p* = <0.0001, Fig. 3A). These similarities potentially suggest that the effects of mutations on readthrough mirror their effects on expression-mutations that suppress folding on the ribosome may also promote cellular degradation. Indeed, trends in variant GFP: mKate intensity ratio are also broadly correlated with computational estimates of mutagenic perturbations of the thermodynamic stability of the native DHFR conformation as determined by ThermoMPNN^38^ (Spearman’s ρ = -0.64, *p* = <0.0001, Fig. 3B). This observation suggests that the structural intermediates that induce readthrough are likely to resemble the native structure. Consistent with this observation, residues that are buried within the core of the native fold exhibit lower mutational tolerance (Spearman’s ρ = 0.57, p = <0.0001, Fig. 3C). Interestingly, mutagenic trends in readthrough efficiency are also highly correlated with previously reported measurements describing the extent to which DHFR variant fitness depends on the cytosolic Lon protease protein^39^-mutations that suppress folding on the ribosome also exhibit proteolytic susceptibility *in vivo* (Spearman’s ρ = 0.42, *p* = <0.0001, Fig. 3D). Notably, this correlation is much weaker upon deactivation of the SecM stalling sequence (Spearman’s ρ = 0.11, *p* = <0.0001, Fig. 3E), suggesting the cotranslational effects of these mutations is most closely associated with susceptibility to Lon-mediated degradation. Taken together, these observations strongly suggest that the formation of the native fold on the ribosome is the dominant factor that drives readthrough and that this conformational transition is potentially consequential for DHFR proteostasis.

**Figure 3.**
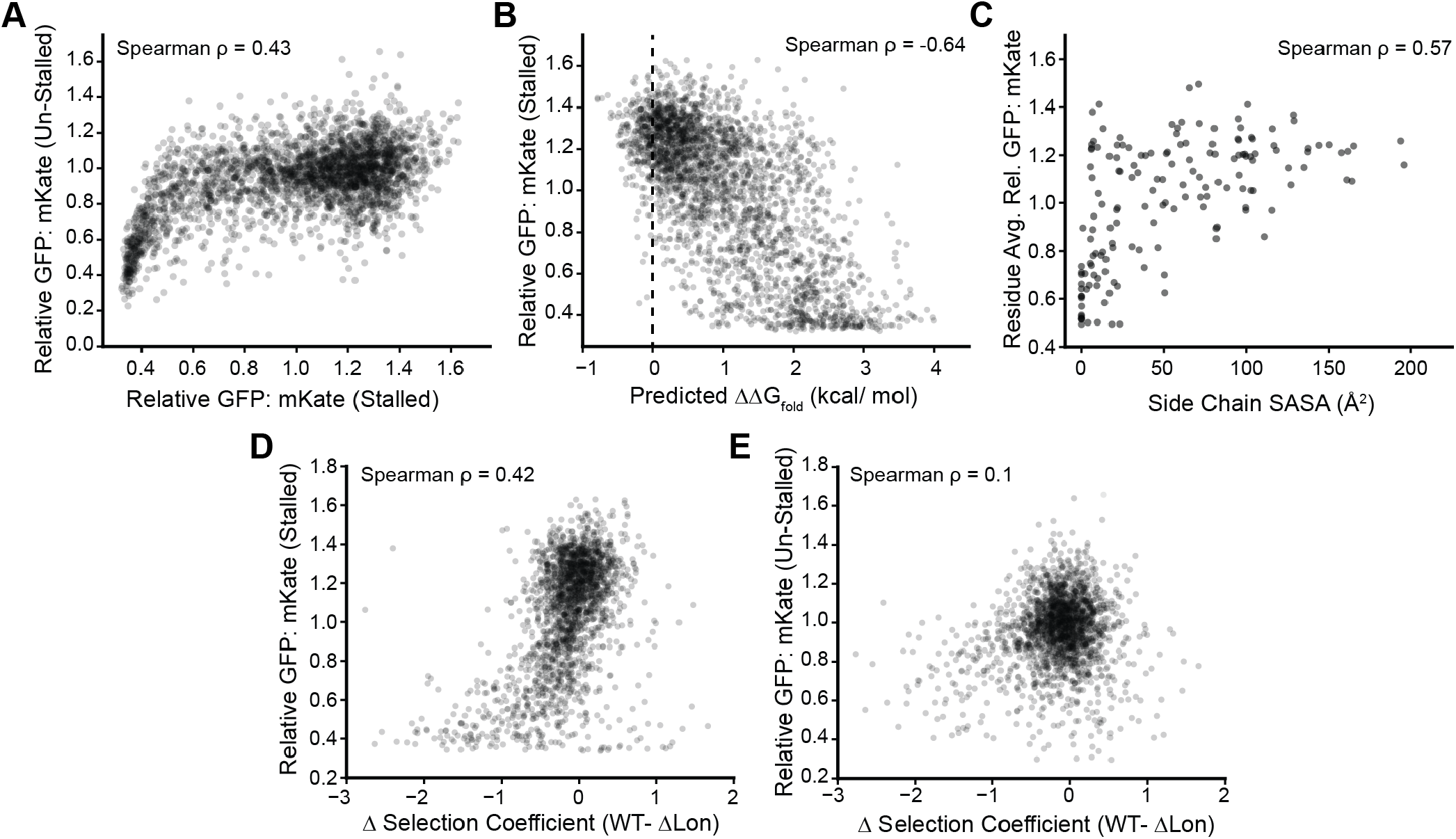
Mutagenic Trends in Variant Readthrough Efficiency. Deep mutational scanning (DMS) measurements comparing the effects of 9,480 codon-level DHFR mutations on the efficiency of SecM readthrough are analyzed in relation to their structural, energetic, biological contexts. For this analysis, variants encoding the same amino acid substitution were collapsed to the amino acid level, yielding 2,763 unique amino acid variants A) The cellular GFP: mKate intensity ratios produced by each variant in the context of the translationally stalled SecM sensor are plotted against the corresponding DMS measurements in the context of the un-stalled P166A SecM sensor (Spearman ρ = 0.43). B) The cellular GFP: mKate intensity ratios produced in the context of the translationally stalled SecM sensor are plotted against their predicted impact on the free energy of folding (ΔΔG_fold_) as predicted by ThermoMPNN (Spearman ρ = -0.64). C) The average cellular GFP: mKate intensity ratio among missense variants at each position in the context of the stalled sensor are plotted against the corresponding solvent accessible surface area (SASA) of the corresponding side chain in the native DHFR structure (PDB 4PSY, Spearman ρ = 0.57). D-E) The cellular GFP: mKate intensity ratios produced by each variant (N = 2,050) in the context of the D) translationally stalled SecM sensor (Spearman ρ = 0.42) or E) in the context of the un-stalled P166A SecM sensor (Spearman ρ = 0.11) are plotted against previously reported differences in the selection coefficient associated with the activity of DHFR variants in WT *E. coli* relative to that in a Lon protease knockout line.

### Identification of Features that are Critical for DHFR Folding on the Ribosome

A survey of individual variant scores reveals a variety of meaningful trends. Notably, there are several regions that exhibit limited mutational tolerance that map to native secondary structure elements. For instance, SecM readthrough is suppressed by most mutations within the first six β strands (β_A-F_, Fig. 4A), which form a twisted parallel β sheet running through the native protein core (see purple strands in Fig. 5A). However, these mutations also reduce GFP: mKate intensity ratios in the context of the un-stalled sensor, suggesting these mutations disrupt DHFR folding both on and off the ribosome. With the exception of certain residues that pack against this β-sheet (i.e. W30, W47, A81, V99, and Y100), most α-helical residues within the loops that connect the strands exhibit greater mutational tolerance (Fig. 4A). These observations suggest that the formation of the parallel β-sheet within the protein core is generally critical for DHFR folding.

**Figure 4.**
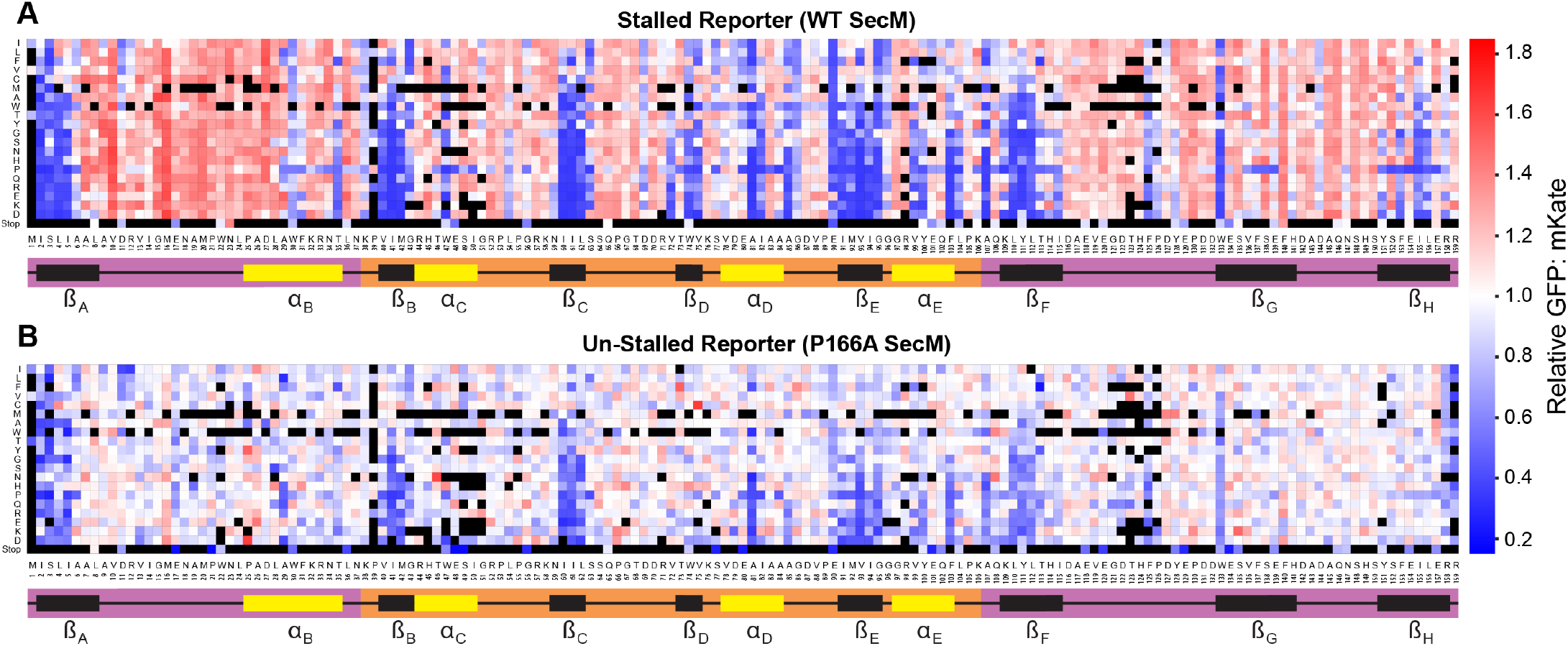
Deep Mutational Scanning of DHFR on the Ribosome. Deep mutational scanning was used to compare the effects of DHFR mutations on SecM readthrough based on changes in cellular GFP: mKate intensity ratios. Heatmaps depict the relative GFP: mKate intensity ratios for each DHFR missense variant in the context of either A) the stalled reporter featuring the WT SecM sequence that induces a translational stall or B) the un-stalled reporter featuring the defective P166A SecM sequence. Values are normalized relative to the GFP: mKate intensity ratio of recombinant cells expressing WT DHFR in the context of the corresponding sensor construct. Values represent the average across three biological replicates. The native amino acid is plotted on the x-coordinate with the positions of native secondary structures indicated for references. Substitutions are organized from hydrophobic (top) to polar (bottom). Red variants enhance readthrough/ abundance, blue variants suppress readthrough/ abundance, and white variants have no effect. Black variants lack sufficient coverage to generate reliable scores.

**Figure 5.**
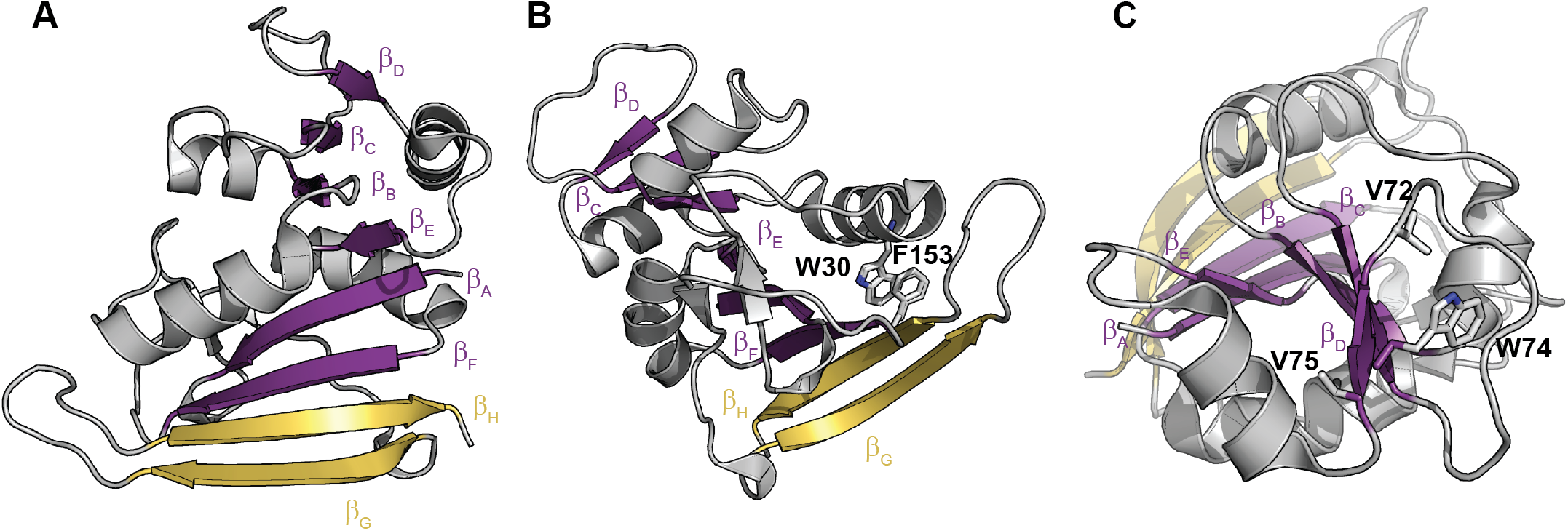
Structural Features Involved in Folding on the Ribosome. Structural features that appear to be critical for DHFR folding on the ribosome are shown in the context of a crystallographic model of the native DHFR fold (PDB 4PSY). A)β sheet (β_A_-β_F_) are shown in purple, while the two C-terminal antiparallel β-strands (β_G_-β_H_) are shown in yellow. B) The structure is rotated to show a key pi stacking interaction between W30 and F153. C) The structure is rotated to show the positions of three buried hydrophobic side chains in the adenosine binding domain (V72, W74, and V75) that are key for folding in the ribosome.

Interestingly, there are several regions of the protein in which mutations strongly attenuate SecM readthrough yet have minimal impact on the signal generated in the context of the un-stalled sensor (Fig. 4). These context-dependent effects highlight contacts that are particularly critical for folding on the ribosome. For instance, the central β sheet is capped off by two anti-parallel β-strands (β_G_ & β_H_, see yellow strands in Fig. 5A) where proline substitutions disproportionately suppress the GFP: mKate intensity in the context of the stalled SecM reporter (Fig. 4). Proline substitutions in these strands are likely to disrupt both backbone hydrogen bonding and to perturb critical tertiary contacts. Tertiary interactions that appear to be particularly important include those formed by a string of hydrophobic side chains in β_H_ (F153, I155, and L156) where polar substitutions selectively suppress readthrough (Figs. 4). In the native structure, F153 forms a sequence-distant pi-stacking interaction with W30 that helps staple together the discontinuous loop domain (Fig. 5B). Importantly, most mutations in W30 also suppress readthrough without decreasing cellular DHFR abundance (Fig. 4). These observations suggest that the formation of these C-terminal antiparallel β -strands and their sequence-distant tertiary interactions is particularly important for folding on the ribosome. Additional context-dependent variations occur within the fourth β strand (β_D_) and its adjacent α-helix (α_D_)-two secondary structure elements within the adenosine binding domain (Fig. 4). These effects are most prominent within a cluster of hydrophobic residues (V72, W74, and V75) within β_D_ that project inward toward the protein core (Fig. 5C). Interestingly, this region was recently found to undergo transient partial unfolding on the ribosome (see *Discussion*).^15^ Together, these identify regions of the protein where mutations specifically perturb DHFR folding on the ribosome.

## Discussion

Protein folding energetics have a dominant influence on protein evolution.^1–4^ Recent analyses of ribosome-nascent chain complexes have revealed that emerging polypeptides populate distinct conformational intermediates during synthesis,^15,16,22^ which suggest the mechanisms of cotranslational folding potentially impose distinct constraints on protein sequence. In this work, we integrate an arrest peptide-based sensor for folding on the ribosome with DMS to survey the constraints of folding on the ribosome. We carried out this analysis in the context of a sensor with a linker that maximizes the coupling between nascent DHFR folding and SecM readthrough (Fig. 1). Interestingly, mutagenic perturbations of readthrough are broadly correlated with those in the context of a deactivated SecM stalling sequence (P166A, Fig. 3A), the latter of which reveals how mutations impact the cellular accumulation of the DHFR-GFP readthrough product. The correlation implies variants that compromise folding on the ribosome also enhance cellular DHFR degradation (Fig. 3A). This interpretation is corroborated by the robust correlation between DMS measurements and computational ΔΔG_fold_ estimates (Fig. 3B). Despite these general trends, our analysis reveals key differences in the mutational tolerance of a central portion of the adenosine binding domain as well as two C-terminal β-strands in the context of the SecM-stalled DHFR (Figs. 4-5). Together, these results provide evidence that the final stages of cotranslational folding may be sensitive to mutagenic perturbations of structural elements that are proximal to the ribosome.

The DMS data reported herein provide complementary insights to those derived from previous investigations of DHFR folding on the ribosome. Recent HDX measurements have elucidated a series of intermediates that are likely to form at certain points in translation.^15^ Interestingly, these investigations revealed that certain positions within the adenosine binding domain exhibit slow exchange kinetics, which typically arise from transient partial unfolding,^40^ once the full-length protein has emerged from the ribosome.^15^ This region includes residues 63-92 (β_D_-α_D_), where we find many mutations selectively perturb folding on the ribosome (Fig. 4). These convergent trends suggest cotranslational folding efficiency is particularly vulnerable to mutations within regions that retain structure during transient partial unfolding on the ribosome. It is also noteworthy that most mutations within a directly-adjacent region (residues 94-117, β_E_-α_E_-β_F_) both suppress readthrough and reduce abundance in the context of an un-stalled sensor. The multifaceted effects of these mutations potentially arise from the fact that this region achieves native-like structure in the context of both a cotranslational folding intermediate^15^ and a partially unfolded kinetic/ equilibrium intermediate (i.e. I_HF_) that is sampled by the full-length protein.^41^ Finally, we note that our maps also show that the two C-terminal strands (β_G_ & β_H_) appear to specifically disrupt folding on the ribosome despite the fact that these segments exhibit prolonged protection in HDX experiments.^15^ This discrepancy suggests that regions proximal to the ribosome may exhibit stronger sequence constrains, potentially due to their heightened propensity to form non-native interactions with the ribosome. When viewed in conjunction wealth of biophysical studies on DHFR folding, our observations show that certain classes of mutations elicit distinct, context-dependent effects during protein folding on the ribosome.

Mutations that destabilize the native fold often reduce evolutionary fitness^4,42^ and can produce long-range epistatic couplings that constrain protein evolution.^6,8,9,43^ Divergence between the folding pathways that operate on and off the ribosome could give rise to distinct sequence constraints associated with cotranslational and post-translational folding coordinates. Indeed, the differences between our two DMS maps reveal specific contexts where this occurs, despite the fact that the ribosome itself tends to buffer the effects of destabilizing mutations.^25^ The extent to which such constraints may weigh on the broader fitness coordinate likely depends on the properties of the protein and its proteostatic context. Full length *E. coli* DHFR can transition from its native conformation to a globally unfolded ensemble (and back) within minutes,^44,45^ which implies that most DHFR molecules sample non-native states at many points during their lifetime. For such proteins, the pioneering round of cotranslational folding may be less consequential and may therefore not have a prominent impact on the fitness coordinate. Indeed, the mutations in β_G_ and β_H_ that suppress cotranslational folding appear to have minimal impact on DHFR fitness,^46^ though they do seem to impact proteostatic fate more generally (Fig. 3 D-E). Nevertheless, mutagenic perturbations of cotranslational processes could weigh more heavily on the fitness coordinate for large and/ or oligomeric proteins that rely on elaborate assembly mechanisms. For instance, the CFTR protein forms dozens of protein-protein interactions during synthesis that are essential for its cotranslational assembly, and perturbations of these interactions promote misfolding and premature degradation. ^47–49^ The relevance of the cotranslational assembly to CFTR fitness landscape is illustrated by the fact that silent mutations that alter translation kinetics can create genetic interactions and alter the biosynthetic yield of functional protein.^50,51^ While it is currently unclear which classes of proteins are likely to face similar constraints, 10-20% of the proteome lacks intrinsic conformational stability^52,53^ while upwards of ∼50% are incapable of spontaneously refolding on biological time scales *in vitro*.^10-11^ Additional investigations are needed to gain a more comprehensive understanding of how cotranslational folding shapes protein evolution.

## Materials and Methods

### Plasmid construction

A SecM-based cotranslational folding reporter was assembled in pETDuet by cloning a DHFR–L56–SecM–GFP cassette into multiple cloning site-I and an mKate expression cassette into multiple cloning site-II of the pETDuet vector using NEBuilder HiFi DNA Assembly Master Mix (New England Biolabs, Ipswich, MA; E2621S). Assembly reactions were incubated at 50 °C for 60 min and transformed into chemically competent 5-alpha *E. coli* (New England Biolabs; C2987I) by heat shock, plated on LB agar supplemented with ampicillin (100 µg/mL), and incubated at 37 °C for 16 h. Single colonies were miniprepped and confirmed by Sanger sequencing (GeneWiz, South Plainfield, NJ). Unless otherwise noted, the L56 tether construct served as the parent template for downstream generation of alternate tether lengths and control variants. A list of the vector sequences used in these studies can be found in Document S1.

### Tether-length variant generation by inverse PCR deletion

Tether-length variants (L54, L52, L50, L48, L46, L44, L42, and L40) were generated from the L56 construct by inverse PCR (whole-plasmid PCR) using back-to-back primers designed to delete the desired number of base pairs from the tether region (Table S2). Reactions (25 µL total) contained 12.5 µL KAPA HiFi HotStart ReadyMix (Roche, Basel, Switzerland; Material No. 07958935001), 20 ng plasmid template, and 0.3 µM each forward and reverse primer. Thermocycling was performed as follows: 95 °C for 3 min; 20 cycles of 98 °C for 20 s, 65 °C for 15 s, and 72 °C for 5 min; followed by 72 °C for 10 min and a 4 °C hold. Reaction products were digested with DpnI to remove parental plasmid by adding 2.5 µL 10× rCutSmart buffer (New England BiolabsIpswitch, MA; B6004S) and 1.25 µL DpnI (New England Biolabs, Ipswitch, MA; R0176S) per reaction (25 U total; corresponding to 1 U per µL of PCR reaction volume) and incubating at 37 °C for 16 h. Digested linear amplicons were purified using DNA Clean & Concentrator-5 (Zymo Research, Irvine, CA; D4004). 300 ng of the resulting linear amplicons were circularized using NEBuilder HiFi DNA Assembly Master Mix as per the manufacturer’s instructions (New England Biolabs, Ipswitch, MA; E2621S), transformed, and verified by Sanger sequencing.

### Site-directed mutagenesis for control constructs and point mutants

Arrest-defective (P166A-SecM) and arrested-length control constructs (stop codon immediately following SecM P166) as well as DHFR point mutants were generated by site-directed mutagenesis using mutagenic primers (0.26 µM each) (Table S2) and PrimeSTAR HS DNA polymerase (Takara Bio USA, San Jose, CA; R010A) using manufacturer’s protocol. Following amplification, reactions were digested with DpnI, purified, transformed and verified by Sanger sequencing.

### Bacterial transformation and expression conditions

Sequence-verified reporter plasmids were transformed into T7 Express *lysY/I*^*q*^ chemically competent *E. coli* (New England Biolabs, Ipswich, MA; C3013I) according to the manufacturer’s protocol and selected on LB agar (Fisher Scientific, Waltham, MA; DF0140-01-0) containing ampicillin (100 µg/mL) at 37 °C for 16 h. Single colonies were used to inoculate 5 mL LB media (Fisher Scientific, Waltham, MA; DF0446-17-3) containing ampicillin (100 µg/mL) and grown overnight at 37 °C with shaking at 250 rpm. Overnight cultures were used to inoculate fresh 5 mL LB containing ampicillin (100 µg/mL) and grown at 37 °C with shaking at 250 rpm to OD600 0.4 - 0.6, at which point expression was induced with IPTG at a final concentration of 1 mM for 2 h under the same conditions; non-induced controls were handled identically without IPTG.

### Sample preparation for flow cytometry

Following induction, cultures were pelleted by centrifugation at 4,000 × g for 5 min, the supernatant was removed, and pellets were washed twice in 1× PBS (pH 7.4; Gibco, Grand Island, NY; 10010-023) and twice in FACS buffer (2% fetal bovine serum in 1× PBS). Cells were resuspended in 1 mL FACS buffer and filtered into 5 mL polystyrene tubes using Flowmi strainers (SP Bel-Art, Wayne, NJ; H13680-0040) prior to acquisition. Samples were diluted in FACS buffer as needed to maintain acquisition rates of ≤3,000 events/s and protected from light until analysis.

### Flow cytometry acquisition and analysis

Flow cytometry was performed on a Bigfoot Cell Sorter (Thermo Fisher Scientific, Waltham, MA) using the following settings: FSC and SSC were collected using the 488 nm laser with FSC-A at 294 V and SSC-A at 376 V; GFP was measured using the 488 nm laser at 600 V (507/19-nm bandpass filter). mKate was measured using the 561 nm laser at either 900 V (for acquisition experiments) or 587 V (for sorting experiments) (625/15-nm bandpass filter). Events were gated sequentially for cells by FSC/SSC, singlets by FSC-H vs FSC-A, and mKate-positive events using a non-induced control and/or untransformed cells to define the mKate gate. For each sample, ≥40,000 total events were collected and ≥10,000 mKate-positive events were retained for analysis. SecM readthrough was quantified as the ratio of mean GFP fluorescence to mean mKate fluorescence for gated mKate-positive singlets.

### DHFR-P166A-SecM Library construction

A modified site-directed mutagenesis protocol was used to introduce the P166A mutation in the SecM motif within the DHFR variant library to generate the corresponding nonfunctional P166A-SecM library. PCR reactions (50 µL total) contained 20 ng of the WT-SecM-DHFR plasmid library, 0.26 µM each forward and reverse mutagenic primer (Table S2), 0.2 mM each dNTP, and 0.5 µL PrimeSTAR HS DNA Polymerase (Takara Bio USA, San Jose, CA; R010A). Thermocycling was performed as follows: 98 °C for 1 min; 15 cycles of 98 °C for 10 s, 55 °C for 5 s, and 72 °C for 7 min 18 s; followed by a 4 °C hold. PCR products were digested with DpnI (New England Biolabs, Ipswich, MA; R0176S) at a ratio of 1 U per µL of PCR reaction volume for 16 h at 37 °C and purified using DNA Clean & Concentrator-5 (Zymo Research, Irvine, CA; D4004). Next, to circularize the mutagenized product, approximately 1.3 µg of purified product was first incubated in T4 polynucleotide kinase reaction buffer at 70 °C for 10 min and immediately chilled on ice. ATP, DTT, and T4 polynucleotide kinase (New England Biolabs, Ipswich, MA; M0201S) were then added to final concentrations of 1 mM ATP and 5 mM DTT and a final enzyme amount of 10 U in a total reaction volume of 50 µL. Phosphorylation reactions were incubated at 37 °C for 30 min, heat-inactivated at 65 °C for 20 min and purified using Zymo DNA Clean & Concentrator-5. 100 ng of phosphorylated DNA was diluted to 5 µL with nuclease-free water, combined with 5 µL Blunt/TA Ligase Master Mix (New England Biolabs, Ipswich, MA; M0367), incubated at room temperature for 15 min, and placed on ice. The ligation product was electroporated into NEB 10-beta Electrocompetent *E. coli* (New England Biolabs, Ipswich, MA; C3020) according to the manufacturer’s instructions. Cells were recovered in NEB 10-beta/Stable Outgrowth Medium for 1 h at 37 °C with shaking at 270 rpm. Serial dilutions of the recovered cells ranging from 10^−1^ to 10^−6^ were prepared, and 50 µL of each dilution was plated on LB agar containing ampicillin (100 µg/mL) to determine the number of colony-forming units ensuring at least a 1000× coverage. The remainder of the recovered cells was transferred to 50 mL LB containing ampicillin (100 µg/mL) and grown for 16 h at 37 °C with shaking at 270 rpm. Plasmid DNA library was isolated from the expanded culture using a ZymoPURE II Plasmid Midiprep Kit (Zymo Research, Irvine, CA; D4201) and used as the DHFR–P166A-SecM library.

### Library transformation, expression, and sorting

The functional SecM and P166A-SecM DHFR libraries were independently transformed into T7 Express *lysY/I*^*q*^ chemically competent *E. coli* (New England Biolabs, Ipswich, MA; C3013I) in three biological replicates using the manufacturer’s heat-shock protocol. Briefly, 500 ng of each plasmid library was added to 50 µL of competent cells, incubated on ice for 30 min, heat-shocked at 42 °C for 10 s, and returned to ice for 5 min. Cells were recovered in 950 µL SOC medium for 1 h at 37 °C with shaking at 250 rpm. Serial dilutions of each recovery culture were plated on LB agar containing ampicillin (100 µg/mL) to determine transformation efficiency and to rule out genetic bottlenecking. Transformations typically achieved 20-40 million colony forming units. The remainder of each recovery culture was transferred to 50 mL LB containing ampicillin (100 µg/mL) and grown for approximately 16 h at 37 °C with shaking at 250 rpm. Overnight cultures were diluted 1:100 into fresh LB containing ampicillin (100 µg/mL) and grown at 37 °C with shaking at 250 rpm until the OD_600_ reached 0.4-0.6. Reporter expression was then induced with 1 mM IPTG (Thermo Fisher Scientific, Waltham, MA) for 2 h under the same conditions, while non-induced controls were handled identically without IPTG. For library sorting, cells were collected by centrifugation at 2,500 × g for 3 min at 4 °C, washed twice with 1× PBS and twice with FACS buffer consisting of 2% fetal bovine serum in 1× PBS, resuspended in FACS buffer, and filtered through Flowmi strainers (SP Bel-Art, Wayne, NJ; H13680-0040). Samples were diluted as needed to maintain acquisition rates below 30,000 events/s and sorted using a Bigfoot Cell Sorter (Thermo Fisher Scientific, Waltham, MA). Events were gated sequentially by forward and side scatter, singlets were isolated using FSC-H and FSC-A, and mKate-positive cells were identified relative to the corresponding non-induced control. mKate-positive singlets were separated into four approximately equally populated bins based on their GFP:mKate fluorescence ratio, and 2 × 10^6^ cells were collected per bin. The mean GFP:mKate fluorescence ratio of each bin was recorded for downstream calculation of variant scores. Sorted cells were pelleted at 4,000 × g for 20 min, resuspended in 5 mL LB media containing ampicillin (100 µg/mL), and grown at 37 °C with shaking at 250 rpm until the OD_600_ reached 0.6-0.8. Cells were then pelleted and stored at −20 °C. Plasmid DNA was isolated from each sorted population using a Zymo plasmid miniprep kit (Zymo Research, Irvine, CA; D4210) and eluted in 10 µL nuclease-free water.

### Preparation of next-generation sequencing libraries

To quantify the relative abundance of DHFR variants within each sorted population, plasmid DNA purified from the expanded post-sort fractions was used to prepare sequencing libraries using a two-step PCR workflow. The entire DHFR coding region was first amplified as a single contiguous amplicon using locus-specific primers containing 5′ and 3′ universal adapter sequences for subsequent indexing. An initial round of PCR reactions was carried out in a final volume of 20 µL containing 100 ng of plasmid DNA, 1 µL each of 10 µM forward and reverse primers, and 10µL of 2× Platinum™ Hot Start PCR Master Mix (Thermo Fisher Scientific, Waltham, MA). Reactions were initially denatured at 95 °C for 5 min, followed by 10 cycles of 98 °C for 20 s, 42 °C for 15 s, and 72 °C for 30 s, and a final extension at 72 °C for 10 min. Amplicons were purified using 0.8 × solid phase reversible immobilization (SPRI) bead cleanup (Beckman Coulter, Brea, CA) and resuspended in 10 µL nuclease free water.

Purified first-round amplicons were then used as templates for a secondary PCR to install sample-specific i5 and i7 combinatorial index pair sequences. Indexing reactions were carried out in a final volume of 20 µL containing 8 µL of purified product from first-round PCR reaction, 2 µL of a 5 µM combined i5/i7 indexing primer mixture, and 10 µL of 2× Platinum™ Hot Start PCR Master Mix (Thermo Fisher Scientific, Waltham, MA). Reactions were initially denatured at 95 °C for 5 min, followed by 12 cycles of 98 °C for 20 s, 47 °C for 15 s, and 72 °C for 30 s, and a final extension at 72 °C for 10 min. The final indexed libraries were purified using a 0.8 × SPRI bead cleanup and resuspended in 20 µL nuclease free water.

Purified indexed libraries were quantified using Qubit™ 1× dsDNA High Sensitivity Assay (Thermo Fisher Scientific, Invitrogen™) and pooled in equal mass. The resulting pool was quantified using the NEBNext^®^ Library Quant Kit for Illumina on an Applied Biosystems StepOne Real-Time PCR system, supplemented with 10% PhiX control library, and diluted to a final loading concentration of 18 pM. Libraries were sequenced on an AVITI System Sequencing Instrument (Element Biosciences, San Diego, CA) using 2×300 cycles on High Output Cloudbreak Freestyle™ sequencing cassette.

### Estimation of SecM readthrough scores from the deep sequencing data

The arrest readthrough scores for each DHFR variant in the pool were calculated from the sequencing reads as described previously.^37^ Briefly, the AVITI sequencing data from each sample were filtered to remove reads having more than one expected error. Reads with average quality score less than 30 and multiple codon errors were also removed. The remaining reads were rarefied to generate the same number of reads across all fractions of a sample. A weighted-average GFP: mKate fluorescence ratio for each variant was calculated as described previously. For ease of comparison the weighted-average scores for each variant was normalized against the weight-average scores for WT DHFR. Weighted-average scores derived from ≤ 50 reads across the four fractions were discarded.

## Acknowledgements

We thank Phillip J. SanMiguel for his technical assistance and acknowledge experimental support from the Bindley Bioscience Center Genomics Core Facility. We would also like to acknowledge Ashish R. Jhangiani for his assistance with the plasmid sample preparations. This work was supported by funding from the National Institute of General Medical Sciences (R35GM152086 to J.P.S.). This manuscript was edited with assistance from ChatGPT-5.6 Sol (Open AI, San Francisco, CA).

## Data and Code Availability

Code for the analysis of sequencing data for deep mutational scanning experiments can be found on the Schlebach Lab GitHub page (https://github.com/schlebachlab). Raw sequencing data are available as an NCBI BioProject (PRJNA1552148). DMS measurements are included in the Supplementary Materials. All other experimental data have been deposited in a Mendeley data directory (doi: 10.17632/vxknfr3j2g.1).

