## Supplemental Materials for "Elucidation of Sequence Constraints for Protein Folding on the Ribosome"

Contents:

-Table S1
-Table S2

**Table S1. Deep Mutational Scanning Measurements of Translationally Stalled DHFR Variants.**

| Variant ID | Average WT-SecM Score n = 3 | WT-SecM SD | Average P166A-SecM Score n = 3 | P166A-SecM SD |
|---|---|---|---|---|
| M1I | 0.750 | 0.033 | 0.634 | 0.054 |
| M1V | 1.051 | 0.082 | 0.902 | 0.072 |
| M1T | 0.779 | 0.088 | 0.447 | 0.078 |
| I2L | 0.885 | 0.061 | 0.737 | 0.082 |
| I2F | 0.662 | 0.060 | 0.784 | 0.050 |
| I2V | 0.765 | 0.105 | 0.721 | 0.130 |
| I2C | 0.348 | 0.030 | 0.428 | 0.051 |
| I2M | 0.614 | 0.016 | 0.758 | 0.130 |
| I2A | 0.331 | 0.042 | 0.383 | 0.056 |
| I2W | 0.449 | 0.031 | 0.437 | 0.084 |
| I2T | 0.524 | 0.050 | 0.794 | 0.087 |
| I2Y | 0.524 | 0.031 | 0.799 | 0.138 |
| I2G | 0.330 | 0.015 | 0.308 | 0.017 |
| I2S | 0.331 | 0.038 | 0.547 | 0.028 |
| I2N | 0.413 | 0.020 | 0.925 | 0.257 |
| I2H | 0.439 | 0.052 | 0.666 | 0.086 |
| I2P | 0.324 | 0.033 | 0.325 | 0.079 |
| I2Q | 0.412 | 0.039 | 0.658 | 0.200 |
| I2R | 0.374 | 0.049 | 0.430 | 0.080 |
| I2E | 0.373 | 0.034 | 0.880 | 0.065 |
| I2K | 0.415 | 0.044 | 1.025 | 0.090 |
| I2D | 0.380 | 0.024 | 0.506 | 0.028 |
| S3I | 0.534 | 0.010 | 0.493 | 0.192 |
| S3L | 0.365 | 0.049 | 0.296 | 0.060 |
| S3V | 0.530 | 0.039 | 0.596 | 0.074 |
| S3C | 0.872 | 0.084 | 0.616 | 0.073 |
| S3A | 0.723 | 0.050 | 0.408 | 0.041 |
| S3T | 0.562 | 0.041 | 0.360 | 0.042 |
| S3Y | 0.417 | 0.043 | 0.380 | 0.129 |
| S3G | 0.724 | 0.095 | 0.503 | 0.104 |
| S3N | 0.749 | 0.020 | 1.001 | 0.059 |
| S3H | 0.468 | 0.057 | 0.596 | 0.064 |
| S3P | 0.335 | 0.031 | 0.265 | 0.071 |
| S3Q | 0.382 | 0.045 | 0.514 | 0.122 |
| S3R | 0.449 | 0.033 | 0.524 | 0.038 |
| S3K | 0.441 | 0.065 | 0.588 | 0.222 |
| S3D | 0.401 | 0.043 | 0.604 | 0.105 |
| L4I | 1.298 | 0.119 | 1.153 | 0.154 |
| L4F | 0.648 | 0.035 | 1.091 | 0.163 |
| L4V | 1.051 | 0.058 | 0.655 | 0.171 |
| L4C | 1.128 | 0.068 | 1.371 | 0.128 |
| L4A | 0.538 | 0.036 | 0.483 | 0.069 |

| Variant ID | Average WT-SecM Score n = 3 | WT-SecM SD | Average P166A-SecM Score n = 3 | P166A-SecM SD |
|---|---|---|---|---|
| L4W | 0.453 | 0.033 | 0.851 | 0.074 |
| L4T | 0.445 | 0.018 | 0.838 | 0.076 |
| L4Y | 0.490 | 0.011 | 0.941 | 0.185 |
| L4G | 0.402 | 0.033 | 0.510 | 0.035 |
| L4S | 0.429 | 0.028 | 0.913 | 0.107 |
| L4N | 0.382 | 0.041 | 0.746 | 0.134 |
| L4H | 0.428 | 0.033 | 0.937 | 0.087 |
| L4P | 0.451 | 0.033 | 0.943 | 0.118 |
| L4Q | 0.487 | 0.045 | 0.800 | 0.137 |
| L4R | 0.392 | 0.038 | 0.717 | 0.100 |
| L4E | 0.391 | 0.067 | 0.605 | 0.047 |
| L4K | 0.373 | 0.051 | 0.474 | 0.134 |
| L4D | 0.337 | 0.031 | 0.594 | 0.111 |
| I5L | 1.063 | 0.049 | 0.945 | 0.148 |
| I5F | 0.969 | 0.066 | 0.997 | 0.271 |
| I5V | 1.010 | 0.081 | 0.750 | 0.077 |
| I5C | 0.855 | 0.059 | 1.065 | 0.213 |
| I5M | 1.062 | 0.098 | 0.839 | 0.110 |
| I5A | 0.459 | 0.040 | 0.441 | 0.062 |
| I5W | 0.457 | 0.033 | 0.261 | 0.053 |
| I5T | 0.650 | 0.050 | 0.788 | 0.052 |
| I5Y | 0.882 | 0.064 | 0.840 | 0.146 |
| I5G | 0.351 | 0.036 | 0.298 | 0.038 |
| I5S | 0.496 | 0.030 | 0.632 | 0.089 |
| I5N | 0.424 | 0.036 | 0.825 | 0.147 |
| I5H | 0.870 | 0.048 | 1.075 | 0.193 |
| I5P | 0.342 | 0.046 | 0.364 | 0.069 |
| I5Q | 0.465 | 0.037 | 0.560 | 0.097 |
| I5R | 0.392 | 0.035 | 0.323 | 0.050 |
| I5E | 0.383 | 0.022 | 0.495 | 0.043 |
| I5K | 0.443 | 0.025 | 0.602 | 0.103 |
| I5D | 0.369 | 0.048 | 0.565 | 0.132 |
| A6I | 0.813 | 0.038 | 0.678 | 0.008 |
| A6L | 0.740 | 0.037 | 0.536 | 0.073 |
| A6F | 0.778 | 0.053 | 0.624 | 0.057 |
| A6V | 1.003 | 0.046 | 0.921 | 0.030 |
| A6C | 1.342 | 0.083 | 1.087 | 0.114 |
| A6M | 0.838 | 0.086 | 0.879 | 0.091 |
| A6W | 0.637 | 0.023 | 0.671 | 0.034 |
| A6T | 1.205 | 0.052 | 1.025 | 0.140 |
| A6Y | 0.866 | 0.081 | 0.858 | 0.147 |
| A6G | 1.143 | 0.041 | 1.019 | 0.176 |
| A6S | 1.215 | 0.067 | 0.891 | 0.035 |

| Variant ID | Average WT-SecM Score n = 3 | WT-SecM SD | Average P166A-SecM Score n = 3 | P166A-SecM SD |
|---|---|---|---|---|
| A6N | 1.099 | 0.050 | 0.959 | 0.190 |
| A6H | 1.009 | 0.042 | 1.084 | 0.159 |
| A6P | 0.526 | 0.042 | 0.477 | 0.042 |
| A6Q | 1.069 | 0.077 | 1.096 | 0.100 |
| A6R | 0.822 | 0.028 | 1.012 | 0.065 |
| A6E | 0.764 | 0.040 | 0.844 | 0.250 |
| A6K | 0.934 | 0.073 | 1.293 | 0.376 |
| A6D | 0.759 | 0.025 | 1.012 | 0.229 |
| A7I | 1.405 | 0.113 | 1.210 | 0.029 |
| A7L | 1.333 | 0.063 | 1.083 | 0.159 |
| A7F | 1.016 | 0.069 | 0.922 | 0.078 |
| A7V | 1.325 | 0.069 | 1.111 | 0.125 |
| A7C | 1.493 | 0.076 | 1.281 | 0.174 |
| A7W | 1.241 | 0.132 | 0.877 | 0.361 |
| A7T | 1.370 | 0.083 | 0.987 | 0.029 |
| A7Y | 1.243 | 0.040 | 0.938 | 0.102 |
| A7G | 1.323 | 0.076 | 0.897 | 0.088 |
| A7S | 1.419 | 0.082 | 1.109 | 0.146 |
| A7N | 1.246 | 0.074 | 1.362 | 0.086 |
| A7H | 1.492 | 0.085 | 1.060 | 0.022 |
| A7P | 1.437 | 0.097 | 1.007 | 0.152 |
| A7Q | 1.412 | 0.107 | 1.011 | 0.101 |
| A7R | 1.477 | 0.081 | 0.903 | 0.096 |
| A7E | 1.036 | 0.075 | 1.177 | 0.145 |
| A7K | 1.397 | 0.096 | 0.943 | 0.098 |
| A7D | 0.927 | 0.033 | 1.065 | 0.116 |
| L8I | 1.311 | 0.052 | 1.046 | 0.147 |
| L8F | 1.356 | 0.083 | 1.029 | 0.146 |
| L8V | 1.194 | 0.077 | 1.021 | 0.100 |
| L8C | 1.451 | 0.054 | 1.183 | 0.198 |
| L8M | 1.362 | 0.067 | 0.790 | 0.132 |
| L8A | 1.263 | 0.098 | 1.180 | 0.089 |
| L8T | 1.314 | 0.081 | 1.298 | 0.168 |
| L8Y | 1.349 | 0.045 | 1.124 | 0.083 |
| L8G | 1.000 | 0.062 | 0.889 | 0.101 |
| L8S | 1.182 | 0.094 | 0.875 | 0.105 |
| L8N | 1.099 | 0.099 | 0.986 | 0.244 |
| L8H | 1.353 | 0.070 | 1.045 | 0.139 |
| L8P | 1.490 | 0.090 | 1.044 | 0.109 |
| L8Q | 1.280 | 0.056 | 1.180 | 0.013 |
| L8R | 1.081 | 0.070 | 0.938 | 0.111 |
| L8E | 0.947 | 0.045 | 1.190 | 0.197 |
| L8K | 1.160 | 0.050 | 0.966 | 0.077 |

| Variant ID | Average WT-SecM Score n = 3 | WT-SecM SD | Average P166A-SecM Score n = 3 | P166A-SecM SD |
|---|---|---|---|---|
| L8D | 0.761 | 0.054 | 1.045 | 0.144 |
| A9I | 0.761 | 0.016 | 0.631 | 0.095 |
| A9L | 0.978 | 0.057 | 0.881 | 0.098 |
| A9F | 1.372 | 0.112 | 0.873 | 0.059 |
| A9V | 1.016 | 0.047 | 1.042 | 0.082 |
| A9C | 1.421 | 0.103 | 1.020 | 0.306 |
| A9M | 1.183 | 0.096 | 0.954 | 0.189 |
| A9W | 1.306 | 0.178 | 1.133 | 0.238 |
| A9T | 1.401 | 0.064 | 0.976 | 0.162 |
| A9Y | 1.543 | 0.068 | 1.156 | 0.167 |
| A9G | 1.519 | 0.107 | 1.135 | 0.152 |
| A9S | 1.234 | 0.061 | 1.064 | 0.104 |
| A9N | 1.466 | 0.135 | 0.912 | 0.261 |
| A9H | 1.456 | 0.093 | 0.913 | 0.095 |
| A9P | 1.395 | 0.075 | 0.941 | 0.132 |
| A9Q | 1.233 | 0.115 | 1.122 | 0.072 |
| A9R | 1.212 | 0.070 | 1.084 | 0.180 |
| A9E | 0.949 | 0.046 | 0.840 | 0.052 |
| A9K | 1.126 | 0.043 | 0.986 | 0.182 |
| A9D | 1.228 | 0.111 | 1.067 | 0.124 |
| V10I | 1.311 | 0.048 | 1.028 | 0.163 |
| V10L | 1.458 | 0.069 | 1.033 | 0.152 |
| V10F | 1.481 | 0.068 | 1.018 | 0.043 |
| V10C | 1.506 | 0.084 | 1.020 | 0.056 |
| V10M | 1.395 | 0.053 | 0.788 | 0.103 |
| V10A | 1.287 | 0.074 | 0.980 | 0.056 |
| V10W | 1.570 | 0.090 | 1.308 | 0.132 |
| V10T | 1.297 | 0.062 | 0.790 | 0.107 |
| V10Y | 1.503 | 0.041 | 0.998 | 0.157 |
| V10G | 1.548 | 0.055 | 1.020 | 0.152 |
| V10S | 1.557 | 0.084 | 1.052 | 0.160 |
| V10N | 1.628 | 0.085 | 1.148 | 0.067 |
| V10H | 1.610 | 0.077 | 1.155 | 0.150 |
| V10P | 1.532 | 0.055 | 1.198 | 0.152 |
| V10Q | 1.592 | 0.049 | 1.080 | 0.078 |
| V10R | 1.544 | 0.076 | 1.090 | 0.215 |
| V10E | 1.422 | 0.102 | 0.936 | 0.112 |
| V10K | 1.554 | 0.083 | 1.149 | 0.191 |
| V10D | 1.427 | 0.090 | 1.107 | 0.280 |
| D11I | 0.522 | 0.031 | 0.392 | 0.062 |
| D11L | 0.661 | 0.025 | 0.647 | 0.056 |
| D11F | 0.728 | 0.050 | 0.601 | 0.126 |
| D11V | 0.576 | 0.012 | 0.613 | 0.097 |

| Variant ID | Average WT-SecM Score n = 3 | WT-SecM SD | Average P166A-SecM Score n = 3 | P166A-SecM SD |
|---|---|---|---|---|
| D11C | 0.965 | 0.044 | 0.909 | 0.143 |
| D11M | 0.730 | 0.016 | 0.881 | 0.273 |
| D11A | 0.891 | 0.039 | 1.108 | 0.117 |
| D11W | 0.641 | 0.044 | 0.552 | 0.089 |
| D11T | 1.003 | 0.052 | 1.103 | 0.129 |
| D11Y | 0.930 | 0.053 | 0.857 | 0.090 |
| D11G | 1.128 | 0.082 | 0.901 | 0.109 |
| D11S | 1.049 | 0.058 | 1.020 | 0.108 |
| D11N | 1.130 | 0.088 | 1.094 | 0.077 |
| D11H | 1.227 | 0.057 | 1.058 | 0.177 |
| D11P | 1.175 | 0.070 | 0.951 | 0.156 |
| D11Q | 1.074 | 0.073 | 1.031 | 0.289 |
| D11R | 1.434 | 0.103 | 1.107 | 0.126 |
| D11E | 1.100 | 0.083 | 1.146 | 0.050 |
| D11K | 1.317 | 0.070 | 0.894 | 0.242 |
| R12I | 0.730 | 0.053 | 0.352 | 0.044 |
| R12L | 0.911 | 0.043 | 0.698 | 0.091 |
| R12F | 0.972 | 0.070 | 0.800 | 0.066 |
| R12V | 0.883 | 0.044 | 0.735 | 0.069 |
| R12C | 1.046 | 0.076 | 0.965 | 0.040 |
| R12A | 1.264 | 0.068 | 0.971 | 0.015 |
| R12T | 1.122 | 0.103 | 0.814 | 0.036 |
| R12Y | 1.046 | 0.095 | 0.826 | 0.128 |
| R12G | 1.344 | 0.070 | 0.924 | 0.041 |
| R12S | 1.269 | 0.051 | 1.089 | 0.084 |
| R12N | 1.316 | 0.095 | 1.195 | 0.416 |
| R12H | 1.097 | 0.049 | 0.996 | 0.024 |
| R12P | 0.910 | 0.057 | 0.781 | 0.088 |
| R12Q | 1.261 | 0.163 | 0.922 | 0.241 |
| R12E | 1.107 | 0.008 | 1.019 | 0.195 |
| R12K | 1.413 | 0.085 | 1.036 | 0.116 |
| R12D | 1.258 | 0.012 | 0.998 | 0.234 |
| V13I | 1.212 | 0.055 | 0.976 | 0.077 |
| V13L | 1.297 | 0.059 | 0.928 | 0.086 |
| V13F | 1.228 | 0.059 | 0.897 | 0.090 |
| V13C | 1.475 | 0.068 | 1.278 | 0.320 |
| V13M | 1.277 | 0.139 | 1.107 | 0.181 |
| V13A | 1.193 | 0.057 | 1.006 | 0.103 |
| V13W | 1.183 | 0.025 | 0.962 | 0.115 |
| V13T | 1.349 | 0.114 | 0.892 | 0.193 |
| V13Y | 1.196 | 0.109 | 0.976 | 0.285 |
| V13G | 1.219 | 0.046 | 0.972 | 0.148 |
| V13S | 1.267 | 0.043 | 0.920 | 0.129 |

| Variant ID | Average WT-SecM Score n = 3 | WT-SecM SD | Average P166A-SecM Score n = 3 | P166A-SecM SD |
|---|---|---|---|---|
| V13N | 1.259 | 0.081 | 0.725 | 0.191 |
| V13H | 1.248 | 0.045 | 1.116 | 0.025 |
| V13P | 0.910 | 0.048 | 0.971 | 0.120 |
| V13Q | 1.299 | 0.007 | 0.999 | 0.202 |
| V13R | 1.489 | 0.070 | 0.934 | 0.122 |
| V13E | 1.102 | 0.067 | 1.108 | 0.148 |
| V13K | 1.480 | 0.054 | 0.657 | 0.058 |
| V13D | 0.994 | 0.030 | 0.583 | 0.084 |
| I14L | 1.364 | 0.061 | 0.961 | 0.075 |
| I14F | 1.205 | 0.066 | 0.929 | 0.016 |
| I14V | 1.295 | 0.075 | 0.815 | 0.039 |
| I14C | 1.297 | 0.110 | 0.694 | 0.181 |
| I14M | 0.969 | 0.010 | 0.934 | 0.012 |
| I14A | 1.080 | 0.061 | 1.039 | 0.069 |
| I14T | 1.252 | 0.077 | 0.843 | 0.107 |
| I14Y | 1.104 | 0.053 | 0.956 | 0.142 |
| I14G | 1.125 | 0.087 | 0.778 | 0.138 |
| I14S | 1.245 | 0.086 | 0.764 | 0.110 |
| I14N | 1.053 | 0.089 | 1.123 | 0.167 |
| I14H | 1.120 | 0.050 | 0.854 | 0.115 |
| I14P | 0.997 | 0.053 | 0.965 | 0.075 |
| I14Q | 1.087 | 0.079 | 0.801 | 0.075 |
| I14R | 1.132 | 0.047 | 0.961 | 0.170 |
| I14E | 1.008 | 0.075 | 1.029 | 0.080 |
| I14K | 1.143 | 0.101 | 0.806 | 0.162 |
| I14D | 1.037 | 0.039 | 0.813 | 0.184 |
| G15I | 1.342 | 0.066 | 1.024 | 0.180 |
| G15L | 1.328 | 0.061 | 0.950 | 0.089 |
| G15F | 1.354 | 0.111 | 1.031 | 0.089 |
| G15V | 1.398 | 0.070 | 1.003 | 0.195 |
| G15C | 1.137 | 0.055 | 0.957 | 0.052 |
| G15M | 1.366 | 0.103 | 0.972 | 0.085 |
| G15A | 1.624 | 0.102 | 1.287 | 0.129 |
| G15W | 1.334 | 0.088 | 0.992 | 0.126 |
| G15T | 1.495 | 0.055 | 1.098 | 0.158 |
| G15Y | 1.462 | 0.105 | 1.056 | 0.107 |
| G15S | 1.080 | 0.057 | 0.800 | 0.085 |
| G15N | 1.491 | 0.079 | 1.169 | 0.151 |
| G15H | 1.433 | 0.084 | 1.037 | 0.126 |
| G15P | 1.497 | 0.105 | 0.986 | 0.041 |
| G15Q | 1.432 | 0.110 | 1.012 | 0.093 |
| G15R | 1.265 | 0.087 | 1.016 | 0.079 |
| G15E | 1.467 | 0.081 | 1.189 | 0.191 |

| Variant ID | Average WT-SecM Score n = 3 | WT-SecM SD | Average P166A-SecM Score n = 3 | P166A-SecM SD |
|---|---|---|---|---|
| G15K | 1.499 | 0.088 | 0.830 | 0.251 |
| G15D | 1.304 | 0.048 | 0.996 | 0.070 |
| M16I | 1.012 | 0.020 | 1.008 | 0.033 |
| M16L | 1.525 | 0.068 | 1.087 | 0.172 |
| M16F | 1.523 | 0.068 | 1.207 | 0.155 |
| M16V | 1.481 | 0.089 | 1.127 | 0.192 |
| M16C | 1.556 | 0.046 | 1.177 | 0.124 |
| M16A | 1.599 | 0.068 | 1.282 | 0.218 |
| M16W | 1.321 | 0.090 | 1.008 | 0.109 |
| M16T | 1.363 | 0.075 | 1.047 | 0.121 |
| M16Y | 1.544 | 0.074 | 1.052 | 0.167 |
| M16G | 1.524 | 0.052 | 0.890 | 0.133 |
| M16S | 1.535 | 0.086 | 1.141 | 0.126 |
| M16N | 1.507 | 0.064 | 1.312 | 0.086 |
| M16H | 1.556 | 0.062 | 1.037 | 0.067 |
| M16P | 1.583 | 0.077 | 1.038 | 0.105 |
| M16Q | 1.519 | 0.046 | 1.253 | 0.260 |
| M16R | 1.573 | 0.065 | 1.228 | 0.135 |
| M16E | 1.594 | 0.064 | 1.264 | 0.180 |
| M16K | 1.473 | 0.108 | 1.217 | 0.042 |
| M16D | 1.629 | 0.078 | 1.367 | 0.209 |
| E17I | 1.015 | 0.062 | 0.770 | 0.100 |
| E17L | 1.059 | 0.044 | 0.798 | 0.055 |
| E17F | 1.134 | 0.081 | 0.888 | 0.107 |
| E17V | 1.052 | 0.089 | 0.726 | 0.077 |
| E17C | 1.247 | 0.016 | 0.963 | 0.070 |
| E17M | 1.188 | 0.061 | 0.864 | 0.058 |
| E17A | 1.184 | 0.099 | 0.781 | 0.170 |
| E17W | 1.043 | 0.022 | 0.723 | 0.061 |
| E17T | 1.096 | 0.066 | 0.561 | 0.031 |
| E17Y | 1.307 | 0.029 | 0.883 | 0.128 |
| E17G | 1.294 | 0.079 | 0.806 | 0.163 |
| E17S | 1.301 | 0.049 | 0.655 | 0.053 |
| E17N | 1.388 | 0.081 | 1.027 | 0.033 |
| E17H | 1.186 | 0.006 | 0.590 | 0.105 |
| E17P | 1.253 | 0.072 | 0.695 | 0.067 |
| E17Q | 1.408 | 0.069 | 0.632 | 0.106 |
| E17R | 1.348 | 0.072 | 0.772 | 0.118 |
| E17K | 1.260 | 0.085 | 0.967 | 0.191 |
| E17D | 1.416 | 0.087 | 1.141 | 0.134 |
| N18I | 1.126 | 0.061 | 0.799 | 0.120 |
| N18L | 1.179 | 0.038 | 1.023 | 0.204 |
| N18F | 1.130 | 0.090 | 1.061 | 0.136 |

| Variant ID | Average WT-SecM Score n = 3 | WT-SecM SD | Average P166A-SecM Score n = 3 | P166A-SecM SD |
|---|---|---|---|---|
| N18V | 1.227 | 0.066 | 1.009 | 0.050 |
| N18C | 1.391 | 0.074 | 0.969 | 0.050 |
| N18A | 1.368 | 0.070 | 0.891 | 0.088 |
| N18T | 1.353 | 0.092 | 1.057 | 0.050 |
| N18Y | 1.265 | 0.051 | 0.957 | 0.116 |
| N18G | 1.360 | 0.083 | 0.741 | 0.008 |
| N18S | 1.297 | 0.059 | 0.935 | 0.115 |
| N18H | 1.342 | 0.085 | 1.014 | 0.257 |
| N18P | 1.290 | 0.073 | 1.012 | 0.054 |
| N18Q | 1.411 | 0.133 | 0.927 | 0.145 |
| N18R | 1.335 | 0.097 | 0.974 | 0.091 |
| N18E | 1.357 | 0.086 | 0.769 | 0.135 |
| N18K | 1.322 | 0.105 | 0.910 | 0.231 |
| N18D | 1.188 | 0.053 | 0.868 | 0.049 |
| A19I | 1.281 | 0.051 | 1.031 | 0.229 |
| A19L | 1.315 | 0.060 | 0.844 | 0.124 |
| A19F | 1.270 | 0.131 | 0.809 | 0.056 |
| A19V | 1.214 | 0.068 | 0.902 | 0.108 |
| A19C | 1.391 | 0.039 | 0.737 | 0.221 |
| A19W | 1.267 | 0.062 | 0.967 | 0.118 |
| A19T | 1.228 | 0.054 | 0.746 | 0.138 |
| A19Y | 1.298 | 0.169 | 0.682 | 0.034 |
| A19G | 1.432 | 0.063 | 1.044 | 0.072 |
| A19S | 1.378 | 0.094 | 1.109 | 0.044 |
| A19H | 1.409 | 0.102 | 0.511 | 0.146 |
| A19P | 1.323 | 0.078 | 0.804 | 0.047 |
| A19Q | 1.427 | 0.113 | 0.954 | 0.168 |
| A19R | 1.412 | 0.089 | 0.956 | 0.155 |
| A19E | 1.448 | 0.076 | 1.110 | 0.127 |
| A19K | 1.361 | 0.144 | 0.988 | 0.067 |
| A19D | 1.255 | 0.062 | 0.893 | 0.197 |
| M20I | 1.209 | 0.038 | 0.978 | 0.098 |
| M20L | 1.391 | 0.065 | 1.050 | 0.031 |
| M20F | 1.406 | 0.093 | 0.867 | 0.076 |
| M20V | 1.249 | 0.058 | 0.855 | 0.092 |
| M20C | 1.498 | 0.112 | 1.046 | 0.187 |
| M20A | 1.542 | 0.083 | 1.135 | 0.026 |
| M20W | 1.395 | 0.031 | 0.946 | 0.360 |
| M20T | 1.252 | 0.075 | 1.057 | 0.022 |
| M20Y | 1.428 | 0.108 | 0.907 | 0.172 |
| M20G | 1.427 | 0.095 | 1.002 | 0.077 |
| M20S | 1.461 | 0.073 | 1.125 | 0.206 |
| M20N | 1.433 | 0.098 | 0.897 | 0.064 |

| Variant ID | Average WT-SecM Score n = 3 | WT-SecM SD | Average P166A-SecM Score n = 3 | P166A-SecM SD |
|---|---|---|---|---|
| M20H | 1.453 | 0.046 | 1.011 | 0.311 |
| M20P | 1.477 | 0.078 | 0.992 | 0.128 |
| M20Q | 1.470 | 0.077 | 0.878 | 0.137 |
| M20R | 1.469 | 0.088 | 1.036 | 0.267 |
| M20E | 1.449 | 0.118 | 1.170 | 0.084 |
| M20K | 1.351 | 0.086 | 0.803 | 0.070 |
| M20D | 1.445 | 0.092 | 0.958 | 0.080 |
| P21I | 1.400 | 0.077 | 1.157 | 0.135 |
| P21L | 1.259 | 0.081 | 0.892 | 0.102 |
| P21F | 1.346 | 0.138 | 0.989 | 0.217 |
| P21V | 1.377 | 0.062 | 1.221 | 0.192 |
| P21C | 1.391 | 0.022 | 1.021 | 0.068 |
| P21A | 1.318 | 0.085 | 0.939 | 0.026 |
| P21T | 1.283 | 0.077 | 1.029 | 0.105 |
| P21Y | 1.317 | 0.018 | 0.624 | 0.135 |
| P21G | 1.299 | 0.111 | 0.779 | 0.050 |
| P21S | 1.151 | 0.071 | 0.860 | 0.067 |
| P21N | 1.380 | 0.050 | 0.845 | 0.287 |
| P21H | 1.361 | 0.115 | 0.880 | 0.126 |
| P21Q | 1.317 | 0.057 | 1.237 | 0.016 |
| P21R | 1.415 | 0.093 | 1.096 | 0.081 |
| P21E | 1.205 | 0.044 | 0.752 | 0.244 |
| P21K | 1.386 | 0.028 | 0.913 | 0.522 |
| P21D | 1.215 | 0.036 | 0.903 | 0.127 |
| W22I | 1.349 | 0.095 | 0.664 | 0.186 |
| W22L | 1.278 | 0.068 | 0.853 | 0.024 |
| W22V | 1.316 | 0.123 | 1.070 | 0.139 |
| W22C | 1.236 | 0.043 | 0.826 | 0.121 |
| W22A | 1.237 | 0.043 | 1.002 | 0.069 |
| W22T | 1.221 | 0.104 | 0.721 | 0.233 |
| W22G | 1.230 | 0.063 | 0.743 | 0.207 |
| W22S | 1.256 | 0.076 | 0.873 | 0.085 |
| W22H | 1.313 | 0.118 | 0.960 | 0.189 |
| W22P | 1.201 | 0.088 | 0.728 | 0.170 |
| W22Q | 1.227 | 0.070 | 0.982 | 0.253 |
| W22R | 1.115 | 0.043 | 0.857 | 0.151 |
| W22E | 1.199 | 0.054 | 1.130 | 0.132 |
| N23I | 1.352 | 0.083 | 0.806 | 0.158 |
| N23L | 1.457 | 0.101 | 1.048 | 0.138 |
| N23F | 1.461 | 0.060 | 1.055 | 0.177 |
| N23V | 1.463 | 0.123 | 0.727 | 0.102 |
| N23A | 1.358 | 0.127 | 1.141 | 0.188 |
| N23T | 1.328 | 0.079 | 0.887 | 0.126 |

| Variant ID | Average WT-SecM Score n = 3 | WT-SecM SD | Average P166A-SecM Score n = 3 | P166A-SecM SD |
|---|---|---|---|---|
| N23Y | 1.435 | 0.075 | 1.173 | 0.093 |
| N23G | 1.284 | 0.078 | 1.027 | 0.194 |
| N23S | 1.278 | 0.071 | 1.081 | 0.078 |
| N23H | 1.431 | 0.069 | 1.016 | 0.128 |
| N23P | 1.264 | 0.099 | 0.914 | 0.199 |
| N23Q | 1.394 | 0.115 | 0.831 | 0.134 |
| N23R | 1.391 | 0.074 | 0.939 | 0.136 |
| N23E | 1.353 | 0.066 | 0.958 | 0.281 |
| N23K | 1.253 | 0.078 | 1.250 | 0.176 |
| N23D | 1.240 | 0.072 | 1.006 | 0.115 |
| L24I | 1.402 | 0.090 | 1.122 | 0.432 |
| L24F | 1.226 | 0.038 | 0.923 | 0.167 |
| L24V | 1.280 | 0.076 | 1.007 | 0.059 |
| L24C | 1.399 | 0.037 | 0.881 | 0.265 |
| L24M | 1.131 | 0.054 | 1.181 | 0.090 |
| L24A | 1.119 | 0.062 | 0.932 | 0.127 |
| L24T | 1.248 | 0.091 | 0.955 | 0.147 |
| L24Y | 1.301 | 0.086 | 0.915 | 0.072 |
| L24G | 1.251 | 0.085 | 0.958 | 0.163 |
| L24S | 1.325 | 0.048 | 1.075 | 0.043 |
| L24N | 1.378 | 0.065 | 1.022 | 0.311 |
| L24H | 1.227 | 0.034 | 0.893 | 0.195 |
| L24P | 0.992 | 0.060 | 0.956 | 0.136 |
| L24Q | 1.324 | 0.097 | 1.142 | 0.030 |
| L24R | 1.225 | 0.056 | 1.013 | 0.120 |
| L24K | 1.177 | 0.076 | 1.103 | 0.267 |
| L24D | 1.197 | 0.129 | 0.893 | 0.141 |
| P25I | 1.282 | 0.153 | 0.833 | 0.144 |
| P25L | 1.318 | 0.025 | 1.001 | 0.084 |
| P25F | 1.296 | 0.077 | 0.640 | 0.141 |
| P25V | 1.312 | 0.115 | 0.690 | 0.157 |
| P25A | 1.349 | 0.062 | 0.537 | 0.092 |
| P25T | 1.302 | 0.020 | 1.087 | 0.228 |
| P25Y | 1.343 | 0.042 | 1.521 | 0.280 |
| P25G | 1.299 | 0.097 | 1.239 | 0.120 |
| P25S | 1.231 | 0.030 | 0.894 | 0.071 |
| P25N | 1.437 | 0.079 | 1.261 | 0.287 |
| P25H | 1.288 | 0.117 | 0.994 | 0.189 |
| P25Q | 1.451 | 0.079 | 1.115 | 0.122 |
| P25R | 1.303 | 0.075 | 1.162 | 0.199 |
| P25E | 1.461 | 0.124 | 1.177 | 0.212 |
| P25D | 1.403 | 0.133 | 1.638 | 0.143 |
| A26I | 1.197 | 0.037 | 1.370 | 0.179 |

| Variant ID | Average WT-SecM Score n = 3 | WT-SecM SD | Average P166A-SecM Score n = 3 | P166A-SecM SD |
|---|---|---|---|---|
| A26L | 1.262 | 0.104 | 1.075 | 0.183 |
| A26F | 1.272 | 0.081 | 1.012 | 0.073 |
| A26V | 1.039 | 0.076 | 1.091 | 0.116 |
| A26C | 1.360 | 0.120 | 1.145 | 0.141 |
| A26W | 1.244 | 0.115 | 0.976 | 0.248 |
| A26T | 1.127 | 0.051 | 0.728 | 0.118 |
| A26Y | 1.181 | 0.106 | 1.245 | 0.082 |
| A26G | 1.298 | 0.091 | 1.218 | 0.194 |
| A26S | 1.288 | 0.063 | 0.794 | 0.138 |
| A26N | 1.424 | 0.079 | 1.193 | 0.272 |
| A26H | 1.408 | 0.124 | 1.180 | 0.155 |
| A26P | 1.212 | 0.051 | 0.998 | 0.185 |
| A26Q | 1.356 | 0.090 | 1.074 | 0.076 |
| A26R | 1.267 | 0.096 | 1.253 | 0.144 |
| A26E | 1.392 | 0.102 | 1.452 | 0.356 |
| A26K | 1.398 | 0.139 | 1.113 | 0.133 |
| A26D | 1.207 | 0.053 | 0.891 | 0.233 |
| D27I | 1.521 | 0.109 | 1.179 | 0.125 |
| D27L | 1.549 | 0.084 | 0.878 | 0.068 |
| D27F | 1.591 | 0.072 | 1.101 | 0.090 |
| D27V | 1.439 | 0.095 | 0.863 | 0.098 |
| D27C | 1.473 | 0.039 | 0.983 | 0.123 |
| D27M | 1.539 | 0.092 | 1.030 | 0.115 |
| D27A | 1.190 | 0.039 | 0.490 | 0.092 |
| D27T | 1.495 | 0.134 | 0.918 | 0.148 |
| D27Y | 1.443 | 0.119 | 0.846 | 0.220 |
| D27G | 1.341 | 0.042 | 0.988 | 0.033 |
| D27S | 1.456 | 0.071 | 1.043 | 0.177 |
| D27N | 1.274 | 0.084 | 0.969 | 0.116 |
| D27H | 1.498 | 0.077 | 0.963 | 0.119 |
| D27P | 1.488 | 0.069 | 1.008 | 0.078 |
| D27Q | 1.496 | 0.061 | 1.023 | 0.245 |
| D27R | 1.278 | 0.121 | 1.021 | 0.169 |
| D27E | 1.293 | 0.098 | 0.906 | 0.042 |
| D27K | 1.296 | 0.092 | 0.891 | 0.162 |
| L28I | 1.317 | 0.058 | 1.121 | 0.133 |
| L28F | 1.199 | 0.053 | 1.064 | 0.085 |
| L28V | 1.390 | 0.055 | 0.790 | 0.144 |
| L28C | 1.326 | 0.106 | 1.252 | 0.238 |
| L28A | 1.316 | 0.037 | 1.217 | 0.248 |
| L28W | 1.293 | 0.152 | 0.828 | 0.132 |
| L28T | 1.238 | 0.080 | 0.704 | 0.087 |
| L28Y | 1.351 | 0.080 | 0.882 | 0.098 |

| Variant ID | Average WT-SecM Score n = 3 | WT-SecM SD | Average P166A-SecM Score n = 3 | P166A-SecM SD |
|---|---|---|---|---|
| L28G | 1.312 | 0.054 | 1.019 | 0.095 |
| L28S | 0.842 | 0.042 | 0.408 | 0.075 |
| L28H | 1.350 | 0.089 | 1.372 | 0.119 |
| L28P | 1.019 | 0.065 | 1.020 | 0.114 |
| L28Q | 1.056 | 0.090 | 0.633 | 0.203 |
| L28R | 1.374 | 0.040 | 0.981 | 0.153 |
| L28E | 1.352 | 0.080 | 1.111 | 0.093 |
| L28K | 1.380 | 0.028 | 1.102 | 0.021 |
| L28D | 1.293 | 0.063 | 1.067 | 0.326 |
| A29I | 1.363 | 0.098 | 0.966 | 0.183 |
| A29L | 0.485 | 0.064 | 0.302 | 0.045 |
| A29F | 0.647 | 0.047 | 0.587 | 0.046 |
| A29V | 1.193 | 0.128 | 0.956 | 0.078 |
| A29C | 1.141 | 0.072 | 0.955 | 0.099 |
| A29M | 1.480 | 0.060 | 1.241 | 0.502 |
| A29W | 1.210 | 0.178 | 1.001 | 0.160 |
| A29T | 1.179 | 0.047 | 0.880 | 0.057 |
| A29Y | 1.311 | 0.102 | 0.798 | 0.129 |
| A29G | 1.271 | 0.091 | 0.992 | 0.101 |
| A29S | 0.947 | 0.083 | 0.695 | 0.073 |
| A29N | 1.398 | 0.081 | 1.016 | 0.230 |
| A29H | 0.726 | 0.080 | 0.483 | 0.091 |
| A29P | 0.563 | 0.043 | 0.471 | 0.072 |
| A29Q | 0.658 | 0.047 | 0.432 | 0.085 |
| A29R | 0.913 | 0.038 | 0.577 | 0.097 |
| A29E | 1.515 | 0.106 | 1.222 | 0.522 |
| A29K | 1.405 | 0.055 | 0.964 | 0.092 |
| A29D | 0.892 | 0.033 | 0.402 | 0.057 |
| W30I | 0.582 | 0.038 | 0.891 | 0.104 |
| W30L | 0.649 | 0.044 | 0.889 | 0.141 |
| W30F | 0.923 | 0.058 | 0.716 | 0.114 |
| W30V | 0.596 | 0.073 | 0.916 | 0.044 |
| W30C | 0.667 | 0.039 | 0.861 | 0.098 |
| W30M | 0.725 | 0.052 | 1.089 | 0.098 |
| W30A | 0.740 | 0.040 | 0.980 | 0.123 |
| W30T | 0.688 | 0.042 | 0.900 | 0.126 |
| W30Y | 0.926 | 0.034 | 1.083 | 0.102 |
| W30G | 0.663 | 0.057 | 0.754 | 0.116 |
| W30S | 0.754 | 0.052 | 0.895 | 0.066 |
| W30N | 0.883 | 0.077 | 1.094 | 0.280 |
| W30H | 0.960 | 0.039 | 1.118 | 0.158 |
| W30P | 0.566 | 0.037 | 0.651 | 0.059 |
| W30Q | 0.838 | 0.061 | 1.278 | 0.174 |

| Variant ID | Average WT-SecM Score n = 3 | WT-SecM SD | Average P166A-SecM Score n = 3 | P166A-SecM SD |
|---|---|---|---|---|
| W30R | 0.873 | 0.048 | 1.017 | 0.069 |
| W30E | 0.688 | 0.034 | 1.078 | 0.125 |
| W30K | 0.671 | 0.033 | 0.819 | 0.124 |
| W30D | 0.652 | 0.031 | 0.794 | 0.207 |
| F31I | 1.195 | 0.047 | 0.958 | 0.129 |
| F31L | 1.218 | 0.071 | 1.018 | 0.127 |
| F31V | 1.111 | 0.098 | 0.914 | 0.074 |
| F31C | 0.969 | 0.041 | 0.968 | 0.166 |
| F31A | 0.906 | 0.066 | 0.927 | 0.144 |
| F31T | 0.949 | 0.018 | 0.942 | 0.110 |
| F31Y | 0.924 | 0.008 | 1.152 | 0.315 |
| F31G | 0.784 | 0.017 | 1.100 | 0.079 |
| F31S | 0.910 | 0.026 | 1.000 | 0.140 |
| F31N | 0.887 | 0.043 | 1.127 | 0.265 |
| F31H | 0.995 | 0.084 | 1.350 | 0.193 |
| F31P | 0.553 | 0.039 | 0.833 | 0.082 |
| F31Q | 1.135 | 0.072 | 0.945 | 0.144 |
| F31R | 1.230 | 0.066 | 0.895 | 0.019 |
| F31E | 0.590 | 0.036 | 0.619 | 0.041 |
| F31K | 1.081 | 0.055 | 0.989 | 0.112 |
| F31D | 0.573 | 0.048 | 0.883 | 0.096 |
| K32I | 0.934 | 0.052 | 0.790 | 0.024 |
| K32L | 1.132 | 0.070 | 0.788 | 0.099 |
| K32F | 1.235 | 0.091 | 0.961 | 0.051 |
| K32V | 1.058 | 0.030 | 0.747 | 0.125 |
| K32C | 1.224 | 0.084 | 1.075 | 0.015 |
| K32M | 1.272 | 0.092 | 0.719 | 0.349 |
| K32A | 1.234 | 0.049 | 0.972 | 0.120 |
| K32W | 1.317 | 0.074 | 0.893 | 0.096 |
| K32T | 1.264 | 0.095 | 1.186 | 0.228 |
| K32Y | 1.320 | 0.096 | 0.940 | 0.064 |
| K32G | 1.175 | 0.069 | 0.929 | 0.184 |
| K32S | 1.301 | 0.075 | 0.954 | 0.136 |
| K32N | 1.319 | 0.080 | 0.809 | 0.114 |
| K32H | 1.344 | 0.095 | 0.990 | 0.176 |
| K32P | 0.849 | 0.055 | 1.046 | 0.104 |
| K32Q | 1.278 | 0.074 | 1.054 | 0.241 |
| K32R | 1.309 | 0.088 | 0.906 | 0.165 |
| K32E | 1.328 | 0.070 | 0.976 | 0.132 |
| K32D | 1.412 | 0.127 | 0.961 | 0.108 |
| R33I | 1.295 | 0.074 | 0.895 | 0.121 |
| R33L | 1.347 | 0.087 | 0.841 | 0.051 |
| R33F | 1.344 | 0.087 | 0.932 | 0.157 |

| Variant ID | Average WT-SecM Score n = 3 | WT-SecM SD | Average P166A-SecM Score n = 3 | P166A-SecM SD |
|---|---|---|---|---|
| R33V | 0.940 | 0.055 | 0.654 | 0.038 |
| R33C | 1.064 | 0.045 | 0.999 | 0.038 |
| R33M | 1.367 | 0.077 | 1.008 | 0.247 |
| R33A | 1.334 | 0.041 | 1.118 | 0.311 |
| R33T | 1.338 | 0.039 | 0.798 | 0.255 |
| R33Y | 1.332 | 0.052 | 0.922 | 0.117 |
| R33G | 1.044 | 0.063 | 0.854 | 0.019 |
| R33S | 1.338 | 0.078 | 1.164 | 0.115 |
| R33N | 1.253 | 0.047 | 0.693 | 0.288 |
| R33H | 1.143 | 0.032 | 1.002 | 0.082 |
| R33P | 0.660 | 0.020 | 0.778 | 0.124 |
| R33Q | 1.310 | 0.116 | 1.199 | 0.175 |
| R33E | 1.312 | 0.110 | 0.700 | 0.065 |
| R33K | 1.131 | 0.161 | 0.780 | 0.189 |
| R33D | 1.269 | 0.060 | 0.902 | 0.185 |
| N34I | 1.430 | 0.053 | 0.874 | 0.173 |
| N34L | 1.437 | 0.077 | 0.989 | 0.113 |
| N34F | 1.262 | 0.128 | 1.089 | 0.103 |
| N34V | 1.416 | 0.026 | 0.953 | 0.267 |
| N34C | 1.441 | 0.099 | 1.089 | 0.299 |
| N34M | 1.484 | 0.098 | 1.097 | 0.296 |
| N34A | 1.373 | 0.063 | 1.105 | 0.140 |
| N34W | 1.350 | 0.154 | 0.793 | 0.182 |
| N34T | 1.198 | 0.099 | 0.803 | 0.168 |
| N34Y | 1.209 | 0.068 | 0.792 | 0.181 |
| N34G | 0.800 | 0.057 | 1.134 | 0.205 |
| N34S | 1.063 | 0.077 | 0.830 | 0.089 |
| N34H | 1.235 | 0.080 | 1.018 | 0.112 |
| N34P | 0.559 | 0.037 | 0.660 | 0.076 |
| N34Q | 1.395 | 0.115 | 0.849 | 0.162 |
| N34R | 1.116 | 0.069 | 1.105 | 0.081 |
| N34E | 1.081 | 0.153 | 0.958 | 0.024 |
| N34K | 1.054 | 0.079 | 0.953 | 0.113 |
| N34D | 0.858 | 0.068 | 0.810 | 0.110 |
| T35I | 1.045 | 0.057 | 0.829 | 0.035 |
| T35L | 0.971 | 0.059 | 0.863 | 0.161 |
| T35F | 0.425 | 0.042 | 0.429 | 0.110 |
| T35V | 1.118 | 0.060 | 1.058 | 0.082 |
| T35C | 1.270 | 0.075 | 0.990 | 0.116 |
| T35M | 0.851 | 0.017 | 1.112 | 0.043 |
| T35A | 0.935 | 0.059 | 0.926 | 0.113 |
| T35Y | 0.495 | 0.024 | 0.622 | 0.238 |
| T35G | 0.700 | 0.066 | 0.753 | 0.116 |

| Variant ID | Average WT-SecM Score n = 3 | WT-SecM SD | Average P166A-SecM Score n = 3 | P166A-SecM SD |
|---|---|---|---|---|
| T35S | 0.843 | 0.060 | 0.866 | 0.160 |
| T35N | 0.623 | 0.038 | 0.853 | 0.122 |
| T35H | 0.578 | 0.067 | 0.916 | 0.176 |
| T35P | 0.505 | 0.041 | 0.870 | 0.136 |
| T35Q | 0.540 | 0.023 | 0.784 | 0.163 |
| T35R | 0.514 | 0.045 | 0.845 | 0.011 |
| T35E | 0.503 | 0.033 | 0.699 | 0.152 |
| T35K | 0.449 | 0.049 | 0.548 | 0.138 |
| T35D | 0.544 | 0.037 | 0.916 | 0.292 |
| L36I | 1.337 | 0.073 | 1.159 | 0.269 |
| L36F | 1.413 | 0.073 | 1.090 | 0.163 |
| L36V | 1.303 | 0.074 | 0.735 | 0.101 |
| L36C | 1.339 | 0.055 | 1.058 | 0.213 |
| L36A | 1.445 | 0.096 | 0.893 | 0.125 |
| L36T | 1.343 | 0.122 | 0.889 | 0.056 |
| L36Y | 1.382 | 0.115 | 1.187 | 0.030 |
| L36G | 1.200 | 0.056 | 0.877 | 0.246 |
| L36S | 1.374 | 0.074 | 1.116 | 0.120 |
| L36N | 1.352 | 0.180 | 1.012 | 0.048 |
| L36H | 1.526 | 0.094 | 1.146 | 0.207 |
| L36P | 0.800 | 0.039 | 1.114 | 0.113 |
| L36Q | 1.344 | 0.107 | 1.021 | 0.095 |
| L36R | 1.300 | 0.085 | 1.019 | 0.119 |
| L36E | 1.517 | 0.040 | 0.985 | 0.305 |
| L36K | 1.355 | 0.108 | 0.810 | 0.036 |
| L36D | 1.335 | 0.052 | 0.970 | 0.202 |
| N37I | 0.731 | 0.058 | 0.863 | 0.144 |
| N37L | 1.097 | 0.072 | 0.886 | 0.044 |
| N37F | 1.252 | 0.098 | 0.839 | 0.195 |
| N37V | 0.735 | 0.040 | 0.837 | 0.157 |
| N37C | 1.156 | 0.067 | 1.039 | 0.148 |
| N37A | 1.264 | 0.048 | 0.878 | 0.166 |
| N37T | 1.163 | 0.040 | 1.088 | 0.218 |
| N37Y | 1.154 | 0.028 | 1.192 | 0.143 |
| N37G | 1.401 | 0.097 | 0.911 | 0.279 |
| N37S | 1.242 | 0.097 | 0.964 | 0.167 |
| N37H | 1.297 | 0.070 | 1.172 | 0.110 |
| N37P | 0.812 | 0.059 | 1.028 | 0.218 |
| N37Q | 1.255 | 0.084 | 1.010 | 0.234 |
| N37R | 1.125 | 0.107 | 1.179 | 0.147 |
| N37E | 1.208 | 0.038 | 0.950 | 0.072 |
| N37K | 1.175 | 0.100 | 0.930 | 0.187 |
| N37D | 1.224 | 0.088 | 0.924 | 0.074 |

| Variant ID | Average WT-SecM Score n = 3 | WT-SecM SD | Average P166A-SecM Score n = 3 | P166A-SecM SD |
|---|---|---|---|---|
| K38I | 0.653 | 0.039 | 0.926 | 0.073 |
| K38L | 0.768 | 0.030 | 0.952 | 0.129 |
| K38F | 0.802 | 0.050 | 0.969 | 0.216 |
| K38V | 0.733 | 0.036 | 1.026 | 0.079 |
| K38C | 0.881 | 0.069 | 0.982 | 0.184 |
| K38M | 0.661 | 0.036 | 0.851 | 0.182 |
| K38A | 0.894 | 0.067 | 1.038 | 0.157 |
| K38W | 0.699 | 0.049 | 0.859 | 0.265 |
| K38T | 0.821 | 0.056 | 0.885 | 0.172 |
| K38Y | 0.870 | 0.047 | 0.943 | 0.151 |
| K38G | 0.821 | 0.026 | 0.935 | 0.203 |
| K38S | 0.805 | 0.046 | 0.881 | 0.114 |
| K38N | 0.875 | 0.050 | 1.126 | 0.140 |
| K38H | 0.915 | 0.050 | 0.911 | 0.180 |
| K38P | 0.644 | 0.041 | 0.916 | 0.109 |
| K38Q | 0.806 | 0.043 | 0.974 | 0.141 |
| K38R | 1.077 | 0.082 | 0.948 | 0.073 |
| K38E | 0.784 | 0.053 | 1.043 | 0.233 |
| K38D | 0.619 | 0.030 | 0.795 | 0.153 |
| P39L | 0.825 | 0.096 | 0.822 | 0.080 |
| P39A | 1.110 | 0.085 | 1.048 | 0.374 |
| P39S | 0.979 | 0.018 | 0.954 | 0.218 |
| P39R | 0.507 | 0.086 | 0.668 | 0.098 |
| V40I | 1.393 | 0.116 | 0.869 | 0.218 |
| V40L | 1.045 | 0.066 | 1.046 | 0.088 |
| V40F | 0.621 | 0.039 | 1.178 | 0.170 |
| V40C | 1.371 | 0.135 | 1.160 | 0.277 |
| V40M | 1.014 | 0.045 | 0.978 | 0.077 |
| V40A | 0.859 | 0.049 | 0.854 | 0.089 |
| V40W | 0.635 | 0.073 | 1.028 | 0.146 |
| V40T | 1.017 | 0.059 | 1.072 | 0.147 |
| V40Y | 0.489 | 0.043 | 1.010 | 0.159 |
| V40G | 0.432 | 0.036 | 0.606 | 0.070 |
| V40S | 0.611 | 0.049 | 0.992 | 0.153 |
| V40N | 0.513 | 0.027 | 0.839 | 0.239 |
| V40H | 0.450 | 0.026 | 0.926 | 0.139 |
| V40P | 0.365 | 0.038 | 0.445 | 0.057 |
| V40Q | 0.494 | 0.040 | 0.714 | 0.148 |
| V40R | 0.476 | 0.035 | 0.852 | 0.079 |
| V40E | 0.378 | 0.043 | 0.486 | 0.102 |
| V40K | 0.491 | 0.023 | 1.074 | 0.106 |
| V40D | 0.366 | 0.036 | 0.583 | 0.058 |
| I41L | 0.678 | 0.055 | 0.974 | 0.135 |

| Variant ID | Average WT-SecM Score n = 3 | WT-SecM SD | Average P166A-SecM Score n = 3 | P166A-SecM SD |
|---|---|---|---|---|
| I41F | 0.480 | 0.025 | 0.843 | 0.125 |
| I41V | 1.316 | 0.096 | 1.024 | 0.052 |
| I41C | 0.828 | 0.069 | 1.096 | 0.107 |
| I41M | 0.582 | 0.013 | 1.036 | 0.228 |
| I41A | 0.513 | 0.053 | 1.089 | 0.206 |
| I41W | 0.349 | 0.058 | 0.567 | 0.043 |
| I41T | 0.541 | 0.026 | 0.771 | 0.126 |
| I41Y | 0.380 | 0.064 | 0.528 | 0.101 |
| I41G | 0.353 | 0.054 | 0.362 | 0.063 |
| I41S | 0.365 | 0.033 | 0.470 | 0.072 |
| I41N | 0.348 | 0.037 | 0.494 | 0.219 |
| I41H | 0.351 | 0.057 | 0.587 | 0.028 |
| I41P | 0.342 | 0.056 | 0.340 | 0.047 |
| I41Q | 0.338 | 0.051 | 0.373 | 0.059 |
| I41R | 0.352 | 0.045 | 0.487 | 0.053 |
| I41E | 0.346 | 0.030 | 0.515 | 0.043 |
| I41K | 0.335 | 0.053 | 0.515 | 0.128 |
| M42I | 0.963 | 0.075 | 0.999 | 0.042 |
| M42L | 1.029 | 0.067 | 0.903 | 0.130 |
| M42F | 1.108 | 0.060 | 0.730 | 0.144 |
| M42V | 0.882 | 0.045 | 0.917 | 0.133 |
| M42C | 0.843 | 0.074 | 1.293 | 0.208 |
| M42A | 0.648 | 0.015 | 0.946 | 0.305 |
| M42W | 1.116 | 0.063 | 1.044 | 0.159 |
| M42T | 0.654 | 0.052 | 1.070 | 0.045 |
| M42Y | 1.011 | 0.043 | 1.051 | 0.187 |
| M42G | 0.393 | 0.040 | 0.565 | 0.051 |
| M42S | 0.526 | 0.027 | 1.017 | 0.069 |
| M42N | 0.370 | 0.029 | 0.439 | 0.052 |
| M42H | 0.442 | 0.062 | 0.504 | 0.140 |
| M42P | 0.350 | 0.037 | 0.365 | 0.066 |
| M42Q | 0.780 | 0.052 | 0.844 | 0.214 |
| M42R | 0.380 | 0.033 | 0.423 | 0.048 |
| M42E | 0.371 | 0.039 | 0.617 | 0.131 |
| M42K | 0.355 | 0.045 | 0.416 | 0.042 |
| M42D | 0.340 | 0.044 | 0.490 | 0.072 |
| G43I | 0.411 | 0.047 | 0.610 | 0.049 |
| G43L | 0.429 | 0.034 | 0.675 | 0.085 |
| G43F | 0.410 | 0.053 | 0.596 | 0.006 |
| G43V | 0.469 | 0.045 | 0.765 | 0.081 |
| G43C | 1.013 | 0.050 | 0.832 | 0.106 |
| G43A | 1.045 | 0.035 | 0.954 | 0.240 |
| G43W | 0.449 | 0.010 | 0.607 | 0.181 |

| Variant ID | Average<br>WT-SecM Score<br>n = 3 | WT-SecM<br>SD | Average<br>P166A-SecM Score<br>n = 3 | P166A-SecM<br>SD |
|---|---|---|---|---|
| G43T | 0.656 | 0.060 | 0.880 | 0.152 |
| G43Y | 0.475 | 0.042 | 0.819 | 0.253 |
| G43S | 1.140 | 0.077 | 0.824 | 0.122 |
| G43H | 0.755 | 0.034 | 0.602 | 0.207 |
| G43P | 0.594 | 0.028 | 0.777 | 0.124 |
| G43Q | 0.528 | 0.035 | 0.567 | 0.078 |
| G43R | 0.410 | 0.044 | 0.760 | 0.097 |
| G43E | 0.809 | 0.039 | 1.063 | 0.254 |
| G43D | 1.078 | 0.034 | 0.976 | 0.100 |
| R44I | 1.154 | 0.093 | 1.012 | 0.068 |
| R44L | 0.980 | 0.018 | 0.950 | 0.018 |
| R44F | 1.198 | 0.069 | 1.036 | 0.084 |
| R44V | 1.232 | 0.057 | 1.029 | 0.140 |
| R44C | 1.083 | 0.044 | 0.900 | 0.070 |
| R44A | 1.163 | 0.095 | 0.812 | 0.099 |
| R44T | 0.962 | 0.075 | 1.004 | 0.246 |
| R44Y | 1.216 | 0.047 | 0.879 | 0.021 |
| R44G | 1.091 | 0.042 | 1.026 | 0.278 |
| R44S | 1.210 | 0.046 | 1.014 | 0.107 |
| R44N | 1.220 | 0.106 | 0.987 | 0.229 |
| R44H | 0.976 | 0.018 | 0.957 | 0.012 |
| R44P | 1.006 | 0.005 | 0.966 | 0.032 |
| R44Q | 1.054 | 0.088 | 0.941 | 0.082 |
| R44E | 1.191 | 0.077 | 0.880 | 0.353 |
| R44D | 1.181 | 0.043 | 1.139 | 0.131 |
| H45I | 1.251 | 0.096 | 1.071 | 0.441 |
| H45L | 1.316 | 0.076 | 0.923 | 0.118 |
| H45V | 1.438 | 0.098 | 1.273 | 0.133 |
| H45C | 1.384 | 0.114 | 1.051 | 0.174 |
| H45A | 1.371 | 0.069 | 0.984 | 0.175 |
| H45T | 1.429 | 0.076 | 0.924 | 0.253 |
| H45Y | 1.010 | 0.039 | 1.052 | 0.054 |
| H45G | 1.366 | 0.030 | 0.849 | 0.157 |
| H45S | 1.338 | 0.103 | 1.004 | 0.312 |
| H45N | 1.084 | 0.028 | 1.044 | 0.141 |
| H45P | 1.177 | 0.072 | 0.916 | 0.100 |
| H45R | 1.171 | 0.065 | 0.806 | 0.092 |
| H45E | 1.452 | 0.014 | 0.723 | 0.122 |
| T46I | 1.094 | 0.040 | 0.914 | 0.174 |
| T46L | 1.206 | 0.098 | 0.758 | 0.019 |
| T46F | 1.191 | 0.056 | 1.286 | 0.417 |
| T46V | 1.222 | 0.130 | 0.818 | 0.270 |
| T46C | 1.294 | 0.172 | 0.759 | 0.131 |

| Variant ID | Average WT-SecM Score n = 3 | WT-SecM SD | Average P166A-SecM Score n = 3 | P166A-SecM SD |
|---|---|---|---|---|
| T46A | 1.107 | 0.054 | 1.066 | 0.049 |
| T46W | 1.411 | 0.091 | 1.253 | 0.024 |
| T46G | 0.792 | 0.046 | 0.927 | 0.211 |
| T46S | 1.113 | 0.046 | 1.115 | 0.239 |
| T46N | 1.178 | 0.072 | 1.612 | 0.203 |
| T46H | 1.345 | 0.048 | 1.153 | 0.189 |
| T46P | 0.630 | 0.040 | 1.038 | 0.123 |
| T46Q | 1.163 | 0.044 | 1.003 | 0.248 |
| T46R | 0.904 | 0.025 | 0.809 | 0.210 |
| T46E | 1.243 | 0.093 | 1.003 | 0.091 |
| T46K | 1.034 | 0.072 | 0.907 | 0.220 |
| T46D | 0.848 | 0.106 | 0.790 | 0.146 |
| W47I | 0.609 | 0.032 | 0.876 | 0.151 |
| W47L | 1.143 | 0.073 | 0.913 | 0.105 |
| W47F | 1.245 | 0.074 | 1.208 | 0.276 |
| W47V | 0.464 | 0.027 | 0.932 | 0.128 |
| W47C | 0.571 | 0.032 | 0.928 | 0.096 |
| W47A | 0.710 | 0.014 | 1.037 | 0.212 |
| W47T | 0.394 | 0.036 | 0.601 | 0.084 |
| W47Y | 1.242 | 0.054 | 0.941 | 0.104 |
| W47G | 0.502 | 0.048 | 0.851 | 0.078 |
| W47S | 0.518 | 0.019 | 0.887 | 0.088 |
| W47H | 0.899 | 0.040 | 1.372 | 0.125 |
| W47P | 0.354 | 0.048 | 0.498 | 0.143 |
| W47Q | 0.564 | 0.056 | 0.862 | 0.134 |
| W47R | 0.751 | 0.052 | 0.889 | 0.054 |
| W47D | 0.362 | 0.042 | 0.383 | 0.015 |
| E48I | 1.068 | 0.062 | 1.331 | 0.065 |
| E48L | 1.172 | 0.036 | 0.932 | 0.087 |
| E48F | 0.964 | 0.081 | 1.086 | 0.204 |
| E48V | 1.165 | 0.077 | 1.068 | 0.048 |
| E48C | 1.110 | 0.100 | 1.255 | 0.179 |
| E48A | 1.047 | 0.063 | 0.816 | 0.044 |
| E48T | 1.052 | 0.055 | 1.160 | 0.160 |
| E48Y | 1.121 | 0.128 | 1.324 | 0.443 |
| E48G | 0.956 | 0.089 | 0.941 | 0.052 |
| E48S | 1.059 | 0.071 | 0.948 | 0.089 |
| E48H | 1.159 | 0.043 | 0.915 | 0.134 |
| E48P | 0.425 | 0.029 | 0.857 | 0.152 |
| E48Q | 1.189 | 0.048 | 1.337 | 0.167 |
| E48R | 1.118 | 0.060 | 0.935 | 0.044 |
| E48K | 1.001 | 0.040 | 0.922 | 0.141 |
| S49I | 1.297 | 0.055 | 0.737 | 0.071 |

| Variant ID | Average WT-SecM Score n = 3 | WT-SecM SD | Average P166A-SecM Score n = 3 | P166A-SecM SD |
|---|---|---|---|---|
| S49L | 1.223 | 0.080 | 0.809 | 0.062 |
| S49V | 1.251 | 0.042 | 0.691 | 0.231 |
| S49C | 1.294 | 0.078 | 0.667 | 0.132 |
| S49A | 1.381 | 0.018 | 0.862 | 0.174 |
| S49T | 1.257 | 0.057 | 1.008 | 0.173 |
| S49G | 1.334 | 0.089 | 1.087 | 0.339 |
| S49P | 0.763 | 0.076 | 0.827 | 0.141 |
| S49R | 1.373 | 0.066 | 1.094 | 0.078 |
| S49D | 1.245 | 0.082 | 0.688 | 0.155 |
| I50L | 1.346 | 0.038 | 0.865 | 0.234 |
| I50F | 1.333 | 0.104 | 1.181 | 0.421 |
| I50V | 1.122 | 0.104 | 0.771 | 0.079 |
| I50C | 1.221 | 0.049 | 1.037 | 0.173 |
| I50M | 1.340 | 0.184 | 0.757 | 0.118 |
| I50A | 1.202 | 0.085 | 0.830 | 0.410 |
| I50T | 1.097 | 0.064 | 0.941 | 0.096 |
| I50G | 1.010 | 0.087 | 0.849 | 0.255 |
| I50S | 1.288 | 0.060 | 0.906 | 0.388 |
| I50P | 0.488 | 0.030 | 0.862 | 0.084 |
| I50Q | 1.273 | 0.122 | 1.003 | 0.113 |
| I50R | 1.324 | 0.115 | 1.187 | 0.145 |
| G51I | 0.743 | 0.094 | 1.139 | 0.239 |
| G51L | 1.057 | 0.028 | 0.980 | 0.225 |
| G51F | 0.731 | 0.056 | 0.980 | 0.241 |
| G51V | 0.819 | 0.024 | 1.228 | 0.049 |
| G51C | 0.898 | 0.023 | 0.990 | 0.131 |
| G51A | 1.114 | 0.089 | 1.290 | 0.264 |
| G51T | 0.995 | 0.062 | 0.901 | 0.037 |
| G51Y | 0.785 | 0.100 | 0.862 | 0.160 |
| G51S | 1.086 | 0.057 | 1.025 | 0.036 |
| G51P | 1.002 | 0.034 | 1.113 | 0.233 |
| G51R | 1.117 | 0.092 | 1.338 | 0.056 |
| G51D | 1.010 | 0.005 | 0.927 | 0.107 |
| R52I | 1.226 | 0.052 | 1.259 | 0.060 |
| R52L | 1.121 | 0.090 | 1.050 | 0.107 |
| R52F | 1.177 | 0.051 | 0.821 | 0.102 |
| R52V | 1.343 | 0.053 | 1.055 | 0.175 |
| R52C | 1.142 | 0.065 | 1.075 | 0.081 |
| R52A | 1.367 | 0.070 | 0.920 | 0.132 |
| R52W | 1.139 | 0.066 | 1.182 | 0.256 |
| R52T | 1.302 | 0.103 | 1.045 | 0.174 |
| R52Y | 1.194 | 0.061 | 0.842 | 0.118 |
| R52G | 1.361 | 0.095 | 1.121 | 0.192 |

| Variant ID | Average<br>WT-SecM Score<br>n = 3 | WT-SecM<br>SD | Average<br>P166A-SecM Score<br>n = 3 | P166A-SecM<br>SD |
|---|---|---|---|---|
| R52S | 1.289 | 0.052 | 0.961 | 0.086 |
| R52N | 1.328 | 0.054 | 0.858 | 0.090 |
| R52H | 1.137 | 0.040 | 0.981 | 0.094 |
| R52P | 0.773 | 0.064 | 0.969 | 0.154 |
| R52Q | 1.285 | 0.041 | 0.963 | 0.065 |
| R52E | 1.418 | 0.117 | 1.328 | 0.175 |
| R52K | 1.410 | 0.056 | 0.959 | 0.152 |
| R52D | 1.313 | 0.062 | 0.854 | 0.066 |
| P53I | 1.283 | 0.089 | 0.792 | 0.106 |
| P53L | 1.194 | 0.075 | 0.828 | 0.174 |
| P53F | 1.275 | 0.070 | 0.714 | 0.161 |
| P53V | 1.267 | 0.069 | 0.816 | 0.190 |
| P53C | 1.272 | 0.065 | 1.169 | 0.156 |
| P53M | 1.314 | 0.055 | 0.800 | 0.194 |
| P53A | 1.255 | 0.047 | 0.931 | 0.049 |
| P53W | 1.237 | 0.138 | 1.173 | 0.260 |
| P53T | 1.099 | 0.082 | 0.993 | 0.089 |
| P53Y | 1.262 | 0.075 | 0.789 | 0.024 |
| P53G | 1.192 | 0.091 | 0.984 | 0.094 |
| P53S | 1.143 | 0.043 | 0.907 | 0.064 |
| P53N | 1.190 | 0.033 | 0.967 | 0.198 |
| P53H | 1.243 | 0.095 | 0.939 | 0.201 |
| P53Q | 1.220 | 0.062 | 1.058 | 0.073 |
| P53R | 1.222 | 0.077 | 0.908 | 0.118 |
| P53E | 1.262 | 0.078 | 0.934 | 0.237 |
| P53K | 1.205 | 0.082 | 0.963 | 0.219 |
| P53D | 1.287 | 0.077 | 0.791 | 0.076 |
| L54I | 1.177 | 0.087 | 0.904 | 0.081 |
| L54F | 1.163 | 0.112 | 1.163 | 0.026 |
| L54V | 0.993 | 0.041 | 0.936 | 0.102 |
| L54C | 1.006 | 0.107 | 1.131 | 0.226 |
| L54A | 0.825 | 0.045 | 1.006 | 0.053 |
| L54W | 0.926 | 0.052 | 0.736 | 0.153 |
| L54T | 1.114 | 0.062 | 0.907 | 0.089 |
| L54Y | 1.194 | 0.069 | 0.781 | 0.093 |
| L54G | 0.708 | 0.029 | 0.875 | 0.064 |
| L54S | 0.872 | 0.045 | 0.985 | 0.055 |
| L54N | 0.724 | 0.059 | 1.236 | 0.214 |
| L54P | 0.642 | 0.052 | 1.101 | 0.234 |
| L54Q | 0.798 | 0.047 | 1.116 | 0.119 |
| L54R | 0.854 | 0.056 | 1.023 | 0.201 |
| L54E | 0.727 | 0.034 | 1.053 | 0.047 |
| L54K | 0.738 | 0.024 | 0.666 | 0.143 |

| Variant ID | Average WT-SecM Score n = 3 | WT-SecM SD | Average P166A-SecM Score n = 3 | P166A-SecM SD |
|---|---|---|---|---|
| L54D | 0.670 | 0.046 | 0.877 | 0.159 |
| P55I | 1.295 | 0.038 | 0.793 | 0.077 |
| P55L | 1.227 | 0.056 | 0.898 | 0.159 |
| P55F | 1.324 | 0.015 | 0.997 | 0.362 |
| P55V | 1.313 | 0.120 | 0.709 | 0.071 |
| P55C | 1.282 | 0.068 | 1.155 | 0.115 |
| P55A | 1.197 | 0.032 | 0.858 | 0.168 |
| P55T | 1.154 | 0.068 | 0.899 | 0.220 |
| P55G | 1.151 | 0.078 | 1.281 | 0.179 |
| P55S | 1.147 | 0.074 | 0.917 | 0.099 |
| P55H | 1.142 | 0.065 | 0.995 | 0.212 |
| P55Q | 1.043 | 0.087 | 1.075 | 0.113 |
| P55R | 1.090 | 0.039 | 0.825 | 0.097 |
| P55E | 1.239 | 0.038 | 0.940 | 0.037 |
| P55K | 1.067 | 0.131 | 0.733 | 0.141 |
| P55D | 1.169 | 0.017 | 1.304 | 0.593 |
| G56I | 0.662 | 0.036 | 0.873 | 0.076 |
| G56L | 0.914 | 0.065 | 0.949 | 0.085 |
| G56F | 1.066 | 0.045 | 1.030 | 0.113 |
| G56V | 0.863 | 0.033 | 0.867 | 0.072 |
| G56C | 1.160 | 0.087 | 1.021 | 0.072 |
| G56M | 0.858 | 0.066 | 1.014 | 0.248 |
| G56A | 1.165 | 0.060 | 0.903 | 0.063 |
| G56T | 0.961 | 0.048 | 0.911 | 0.175 |
| G56Y | 1.096 | 0.084 | 1.044 | 0.039 |
| G56S | 1.252 | 0.076 | 1.018 | 0.152 |
| G56N | 1.162 | 0.090 | 1.222 | 0.107 |
| G56H | 1.122 | 0.090 | 0.923 | 0.217 |
| G56P | 0.547 | 0.035 | 1.007 | 0.078 |
| G56Q | 1.014 | 0.054 | 1.362 | 0.159 |
| G56R | 0.944 | 0.041 | 0.994 | 0.009 |
| G56E | 1.037 | 0.004 | 0.971 | 0.111 |
| G56K | 0.901 | 0.067 | 1.201 | 0.211 |
| G56D | 1.084 | 0.030 | 1.049 | 0.219 |
| R57I | 1.374 | 0.069 | 0.992 | 0.214 |
| R57L | 1.148 | 0.071 | 0.932 | 0.050 |
| R57F | 1.008 | 0.110 | 0.934 | 0.326 |
| R57V | 1.293 | 0.098 | 1.014 | 0.203 |
| R57C | 1.142 | 0.075 | 0.997 | 0.048 |
| R57M | 1.174 | 0.128 | 1.037 | 0.357 |
| R57A | 1.291 | 0.080 | 0.866 | 0.124 |
| R57T | 1.314 | 0.081 | 0.916 | 0.116 |
| R57Y | 1.100 | 0.093 | 1.226 | 0.226 |

| Variant ID | Average WT-SecM Score n = 3 | WT-SecM SD | Average P166A-SecM Score n = 3 | P166A-SecM SD |
|---|---|---|---|---|
| R57G | 1.220 | 0.095 | 1.036 | 0.011 |
| R57S | 1.407 | 0.081 | 0.812 | 0.115 |
| R57N | 1.365 | 0.092 | 0.994 | 0.413 |
| R57H | 1.053 | 0.072 | 0.965 | 0.033 |
| R57P | 1.047 | 0.087 | 0.773 | 0.069 |
| R57Q | 1.161 | 0.075 | 0.942 | 0.022 |
| R57E | 1.235 | 0.009 | 0.651 | 0.106 |
| R57K | 1.213 | 0.134 | 0.770 | 0.217 |
| R57D | 1.221 | 0.144 | 1.325 | 0.079 |
| K58I | 1.296 | 0.098 | 0.938 | 0.127 |
| K58L | 1.362 | 0.090 | 1.071 | 0.077 |
| K58F | 1.194 | 0.092 | 0.985 | 0.063 |
| K58V | 1.349 | 0.090 | 1.033 | 0.141 |
| K58C | 1.284 | 0.120 | 0.926 | 0.240 |
| K58A | 1.230 | 0.111 | 1.092 | 0.080 |
| K58T | 1.328 | 0.078 | 1.098 | 0.048 |
| K58Y | 1.109 | 0.072 | 1.120 | 0.291 |
| K58G | 0.859 | 0.026 | 0.830 | 0.139 |
| K58S | 1.213 | 0.077 | 1.059 | 0.085 |
| K58N | 1.171 | 0.066 | 0.981 | 0.112 |
| K58H | 1.332 | 0.109 | 1.249 | 0.396 |
| K58P | 1.267 | 0.053 | 1.148 | 0.317 |
| K58Q | 0.984 | 0.023 | 0.905 | 0.052 |
| K58R | 1.229 | 0.098 | 0.961 | 0.144 |
| K58E | 1.108 | 0.109 | 0.890 | 0.072 |
| K58D | 1.071 | 0.073 | 1.100 | 0.123 |
| N59I | 1.276 | 0.119 | 1.201 | 0.249 |
| N59L | 1.287 | 0.083 | 1.111 | 0.101 |
| N59F | 1.363 | 0.048 | 1.026 | 0.207 |
| N59V | 1.299 | 0.061 | 1.177 | 0.255 |
| N59C | 1.366 | 0.072 | 1.132 | 0.204 |
| N59M | 1.294 | 0.054 | 0.663 | 0.197 |
| N59A | 1.144 | 0.087 | 1.070 | 0.090 |
| N59W | 1.088 | 0.092 | 0.912 | 0.184 |
| N59T | 1.336 | 0.038 | 0.879 | 0.122 |
| N59Y | 1.192 | 0.088 | 0.866 | 0.088 |
| N59G | 0.797 | 0.061 | 0.943 | 0.061 |
| N59S | 1.125 | 0.037 | 0.998 | 0.126 |
| N59H | 1.168 | 0.008 | 1.006 | 0.134 |
| N59P | 1.029 | 0.097 | 0.937 | 0.121 |
| N59Q | 1.117 | 0.053 | 0.992 | 0.196 |
| N59R | 0.670 | 0.041 | 0.845 | 0.143 |
| N59E | 0.732 | 0.110 | 0.971 | 0.047 |

| Variant ID | Average WT-SecM Score n = 3 | WT-SecM SD | Average P166A-SecM Score n = 3 | P166A-SecM SD |
|---|---|---|---|---|
| N59K | 0.795 | 0.056 | 1.131 | 0.215 |
| N59D | 0.827 | 0.090 | 0.707 | 0.138 |
| I60L | 0.842 | 0.071 | 1.031 | 0.140 |
| I60F | 0.727 | 0.027 | 0.880 | 0.124 |
| I60V | 0.928 | 0.087 | 1.017 | 0.209 |
| I60C | 0.484 | 0.038 | 1.144 | 0.111 |
| I60M | 0.757 | 0.044 | 0.995 | 0.143 |
| I60A | 0.361 | 0.028 | 0.488 | 0.188 |
| I60W | 0.357 | 0.023 | 0.420 | 0.080 |
| I60T | 0.483 | 0.038 | 0.770 | 0.031 |
| I60Y | 0.660 | 0.042 | 0.775 | 0.139 |
| I60G | 0.345 | 0.037 | 0.416 | 0.052 |
| I60S | 0.374 | 0.033 | 0.519 | 0.032 |
| I60N | 0.357 | 0.039 | 0.561 | 0.170 |
| I60H | 0.377 | 0.050 | 0.721 | 0.065 |
| I60P | 0.344 | 0.035 | 0.407 | 0.030 |
| I60Q | 0.356 | 0.039 | 0.475 | 0.062 |
| I60R | 0.337 | 0.045 | 0.417 | 0.069 |
| I60E | 0.343 | 0.040 | 0.309 | 0.063 |
| I60K | 0.343 | 0.042 | 0.479 | 0.044 |
| I61L | 0.726 | 0.033 | 0.941 | 0.078 |
| I61F | 0.368 | 0.047 | 0.513 | 0.062 |
| I61V | 1.210 | 0.095 | 0.915 | 0.128 |
| I61C | 0.959 | 0.036 | 1.206 | 0.149 |
| I61M | 0.553 | 0.023 | 0.642 | 0.198 |
| I61A | 0.487 | 0.051 | 0.973 | 0.018 |
| I61W | 0.347 | 0.041 | 0.606 | 0.084 |
| I61T | 0.592 | 0.041 | 0.813 | 0.081 |
| I61Y | 0.368 | 0.036 | 0.548 | 0.143 |
| I61G | 0.416 | 0.031 | 0.692 | 0.171 |
| I61S | 0.392 | 0.031 | 0.606 | 0.072 |
| I61N | 0.375 | 0.034 | 0.537 | 0.099 |
| I61H | 0.349 | 0.020 | 0.544 | 0.056 |
| I61P | 0.345 | 0.026 | 0.494 | 0.125 |
| I61Q | 0.350 | 0.042 | 0.405 | 0.015 |
| I61R | 0.343 | 0.039 | 0.420 | 0.082 |
| I61E | 0.326 | 0.033 | 0.337 | 0.145 |
| I61K | 0.345 | 0.063 | 0.586 | 0.025 |
| I61D | 0.348 | 0.039 | 0.439 | 0.202 |
| L62I | 1.240 | 0.099 | 0.920 | 0.118 |
| L62F | 1.075 | 0.077 | 0.902 | 0.082 |
| L62V | 0.963 | 0.064 | 0.874 | 0.094 |
| L62C | 0.592 | 0.053 | 0.790 | 0.127 |

| Variant ID | Average WT-SecM Score n = 3 | WT-SecM SD | Average P166A-SecM Score n = 3 | P166A-SecM SD |
|---|---|---|---|---|
| L62A | 0.455 | 0.018 | 0.891 | 0.164 |
| L62W | 0.767 | 0.080 | 0.814 | 0.399 |
| L62T | 0.469 | 0.051 | 0.854 | 0.175 |
| L62Y | 0.969 | 0.061 | 1.143 | 0.200 |
| L62G | 0.342 | 0.047 | 0.447 | 0.046 |
| L62S | 0.370 | 0.044 | 0.406 | 0.041 |
| L62N | 0.349 | 0.046 | 0.432 | 0.073 |
| L62H | 0.553 | 0.021 | 0.826 | 0.192 |
| L62P | 0.544 | 0.034 | 0.651 | 0.113 |
| L62Q | 0.476 | 0.041 | 0.859 | 0.241 |
| L62R | 0.376 | 0.034 | 0.555 | 0.076 |
| L62E | 0.357 | 0.038 | 0.431 | 0.076 |
| L62K | 0.407 | 0.031 | 0.732 | 0.143 |
| L62D | 0.372 | 0.020 | 0.373 | 0.047 |
| S63I | 0.542 | 0.056 | 0.959 | 0.053 |
| S63L | 0.411 | 0.048 | 0.543 | 0.045 |
| S63F | 0.530 | 0.028 | 0.798 | 0.023 |
| S63V | 0.664 | 0.028 | 0.991 | 0.130 |
| S63C | 1.216 | 0.070 | 0.944 | 0.106 |
| S63M | 0.539 | 0.049 | 1.108 | 0.094 |
| S63A | 1.256 | 0.066 | 0.774 | 0.082 |
| S63W | 0.622 | 0.044 | 1.115 | 0.121 |
| S63T | 1.383 | 0.093 | 0.778 | 0.107 |
| S63Y | 0.611 | 0.065 | 1.116 | 0.056 |
| S63G | 1.031 | 0.074 | 0.978 | 0.064 |
| S63N | 0.862 | 0.038 | 1.064 | 0.073 |
| S63H | 0.872 | 0.013 | 1.030 | 0.131 |
| S63P | 0.357 | 0.033 | 0.408 | 0.054 |
| S63Q | 0.696 | 0.082 | 0.861 | 0.043 |
| S63R | 0.507 | 0.052 | 0.884 | 0.033 |
| S63E | 0.705 | 0.070 | 0.966 | 0.021 |
| S63K | 0.435 | 0.025 | 0.873 | 0.112 |
| S63D | 1.270 | 0.128 | 1.005 | 0.070 |
| S64I | 1.097 | 0.077 | 0.874 | 0.056 |
| S64L | 1.212 | 0.073 | 0.970 | 0.127 |
| S64F | 1.053 | 0.023 | 1.216 | 0.137 |
| S64V | 1.163 | 0.075 | 0.890 | 0.096 |
| S64C | 1.174 | 0.072 | 0.889 | 0.202 |
| S64M | 1.278 | 0.168 | 0.951 | 0.400 |
| S64A | 1.285 | 0.075 | 1.087 | 0.175 |
| S64W | 0.967 | 0.045 | 0.812 | 0.275 |
| S64T | 1.308 | 0.057 | 0.920 | 0.067 |
| S64Y | 1.094 | 0.083 | 1.253 | 0.426 |

| Variant ID | Average WT-SecM Score n = 3 | WT-SecM SD | Average P166A-SecM Score n = 3 | P166A-SecM SD |
|---|---|---|---|---|
| S64G | 1.119 | 0.079 | 0.889 | 0.050 |
| S64N | 1.140 | 0.060 | 1.037 | 0.079 |
| S64H | 1.259 | 0.100 | 1.249 | 0.160 |
| S64P | 1.163 | 0.058 | 1.178 | 0.141 |
| S64Q | 1.322 | 0.076 | 1.310 | 0.155 |
| S64R | 1.169 | 0.068 | 1.003 | 0.101 |
| S64E | 1.379 | 0.091 | 0.845 | 0.217 |
| S64K | 1.318 | 0.085 | 0.728 | 0.121 |
| S64D | 1.317 | 0.081 | 0.838 | 0.146 |
| Q65I | 1.292 | 0.068 | 1.105 | 0.078 |
| Q65L | 1.227 | 0.054 | 1.108 | 0.081 |
| Q65F | 1.217 | 0.059 | 1.155 | 0.310 |
| Q65V | 1.309 | 0.103 | 0.774 | 0.106 |
| Q65C | 1.140 | 0.079 | 0.768 | 0.390 |
| Q65M | 1.246 | 0.098 | 0.925 | 0.226 |
| Q65A | 1.303 | 0.067 | 1.008 | 0.079 |
| Q65T | 1.359 | 0.066 | 1.239 | 0.082 |
| Q65Y | 1.311 | 0.068 | 0.853 | 0.100 |
| Q65G | 1.173 | 0.027 | 0.979 | 0.304 |
| Q65S | 1.272 | 0.099 | 0.890 | 0.115 |
| Q65N | 1.396 | 0.111 | 0.853 | 0.324 |
| Q65H | 1.181 | 0.102 | 1.131 | 0.099 |
| Q65P | 0.853 | 0.033 | 0.988 | 0.131 |
| Q65R | 1.160 | 0.067 | 0.953 | 0.034 |
| Q65E | 1.342 | 0.082 | 1.214 | 0.176 |
| Q65K | 1.334 | 0.035 | 0.906 | 0.139 |
| Q65D | 1.342 | 0.144 | 0.845 | 0.269 |
| P66I | 1.343 | 0.109 | 0.929 | 0.183 |
| P66L | 1.139 | 0.060 | 0.957 | 0.153 |
| P66F | 1.254 | 0.042 | 0.771 | 0.191 |
| P66V | 1.293 | 0.115 | 1.278 | 0.058 |
| P66C | 1.303 | 0.060 | 1.056 | 0.283 |
| P66M | 1.279 | 0.072 | 1.024 | 0.278 |
| P66A | 1.307 | 0.096 | 1.022 | 0.105 |
| P66W | 0.986 | 0.068 | 0.603 | 0.064 |
| P66T | 1.318 | 0.063 | 0.872 | 0.167 |
| P66Y | 1.137 | 0.102 | 1.181 | 0.114 |
| P66G | 1.165 | 0.165 | 0.959 | 0.103 |
| P66S | 1.188 | 0.058 | 1.168 | 0.017 |
| P66N | 1.256 | 0.056 | 1.248 | 0.298 |
| P66H | 1.250 | 0.069 | 1.253 | 0.175 |
| P66Q | 1.265 | 0.014 | 1.059 | 0.278 |
| P66R | 1.239 | 0.057 | 0.962 | 0.143 |

| Variant ID | Average WT-SecM Score n = 3 | WT-SecM SD | Average P166A-SecM Score n = 3 | P166A-SecM SD |
|---|---|---|---|---|
| P66E | 1.358 | 0.157 | 1.032 | 0.111 |
| P66K | 1.366 | 0.090 | 1.408 | 0.378 |
| P66D | 1.361 | 0.067 | 1.344 | 0.262 |
| G67I | 1.239 | 0.094 | 0.728 | 0.146 |
| G67L | 1.242 | 0.087 | 0.932 | 0.178 |
| G67F | 1.043 | 0.057 | 1.223 | 0.099 |
| G67V | 1.198 | 0.049 | 0.896 | 0.062 |
| G67C | 1.183 | 0.037 | 0.953 | 0.164 |
| G67M | 1.184 | 0.071 | 1.086 | 0.149 |
| G67A | 1.135 | 0.056 | 1.151 | 0.236 |
| G67W | 0.873 | 0.034 | 1.012 | 0.299 |
| G67T | 1.145 | 0.067 | 1.043 | 0.131 |
| G67Y | 1.022 | 0.059 | 1.305 | 0.202 |
| G67S | 1.153 | 0.087 | 1.062 | 0.205 |
| G67N | 1.093 | 0.037 | 0.871 | 0.083 |
| G67H | 1.070 | 0.083 | 1.393 | 0.080 |
| G67P | 1.296 | 0.081 | 1.057 | 0.177 |
| G67Q | 1.232 | 0.085 | 1.046 | 0.406 |
| G67R | 1.021 | 0.050 | 1.067 | 0.070 |
| G67K | 1.070 | 0.087 | 1.304 | 0.259 |
| G67D | 1.055 | 0.066 | 0.940 | 0.056 |
| T68I | 1.240 | 0.023 | 1.109 | 0.105 |
| T68L | 1.304 | 0.058 | 1.131 | 0.107 |
| T68F | 1.210 | 0.098 | 1.183 | 0.047 |
| T68V | 1.301 | 0.056 | 1.069 | 0.079 |
| T68C | 1.244 | 0.042 | 0.896 | 0.183 |
| T68M | 1.083 | 0.045 | 0.940 | 0.023 |
| T68A | 1.206 | 0.076 | 1.055 | 0.096 |
| T68Y | 1.281 | 0.021 | 1.129 | 0.398 |
| T68G | 1.295 | 0.058 | 1.034 | 0.173 |
| T68S | 1.343 | 0.084 | 1.074 | 0.058 |
| T68N | 1.369 | 0.129 | 0.829 | 0.175 |
| T68H | 1.296 | 0.116 | 1.108 | 0.218 |
| T68P | 1.205 | 0.029 | 0.971 | 0.068 |
| T68Q | 0.917 | 0.052 | 0.729 | 0.054 |
| T68R | 1.243 | 0.032 | 0.952 | 0.049 |
| T68E | 1.385 | 0.087 | 1.206 | 0.126 |
| T68K | 1.259 | 0.128 | 0.969 | 0.155 |
| T68D | 1.360 | 0.017 | 1.009 | 0.307 |
| D69I | 0.668 | 0.072 | 0.818 | 0.099 |
| D69L | 0.681 | 0.014 | 0.912 | 0.105 |
| D69F | 0.814 | 0.040 | 1.017 | 0.151 |
| D69V | 0.670 | 0.068 | 0.842 | 0.042 |

| Variant ID | Average WT-SecM Score n = 3 | WT-SecM SD | Average P166A-SecM Score n = 3 | P166A-SecM SD |
|---|---|---|---|---|
| D69C | 1.113 | 0.121 | 0.824 | 0.173 |
| D69M | 0.698 | 0.063 | 0.944 | 0.176 |
| D69A | 0.799 | 0.053 | 1.011 | 0.256 |
| D69W | 0.694 | 0.011 | 0.983 | 0.082 |
| D69T | 0.885 | 0.037 | 1.067 | 0.175 |
| D69Y | 0.902 | 0.050 | 0.846 | 0.110 |
| D69G | 0.870 | 0.051 | 0.876 | 0.041 |
| D69S | 1.048 | 0.063 | 1.023 | 0.106 |
| D69N | 0.948 | 0.035 | 0.925 | 0.040 |
| D69H | 0.896 | 0.037 | 0.759 | 0.126 |
| D69P | 0.642 | 0.028 | 0.855 | 0.193 |
| D69Q | 0.756 | 0.066 | 0.948 | 0.206 |
| D69R | 0.579 | 0.046 | 1.078 | 0.111 |
| D69E | 1.000 | 0.063 | 0.931 | 0.199 |
| D69K | 0.636 | 0.020 | 1.196 | 0.201 |
| D70I | 1.203 | 0.077 | 1.115 | 0.310 |
| D70L | 1.253 | 0.052 | 0.981 | 0.117 |
| D70F | 1.058 | 0.078 | 1.108 | 0.185 |
| D70V | 1.217 | 0.095 | 1.096 | 0.242 |
| D70C | 1.304 | 0.104 | 0.701 | 0.119 |
| D70M | 1.210 | 0.078 | 1.252 | 0.185 |
| D70A | 1.285 | 0.053 | 1.126 | 0.104 |
| D70T | 1.361 | 0.036 | 1.089 | 0.201 |
| D70Y | 1.198 | 0.034 | 0.997 | 0.040 |
| D70G | 1.096 | 0.063 | 0.992 | 0.050 |
| D70S | 1.401 | 0.053 | 1.081 | 0.099 |
| D70N | 1.146 | 0.058 | 0.932 | 0.054 |
| D70H | 1.292 | 0.039 | 1.000 | 0.061 |
| D70P | 1.366 | 0.077 | 1.080 | 0.155 |
| D70Q | 1.351 | 0.060 | 1.013 | 0.183 |
| D70R | 1.208 | 0.127 | 1.018 | 0.172 |
| D70E | 1.231 | 0.106 | 0.906 | 0.245 |
| D70K | 1.327 | 0.080 | 0.926 | 0.178 |
| R71I | 0.921 | 0.062 | 1.198 | 0.019 |
| R71L | 0.958 | 0.022 | 0.838 | 0.052 |
| R71F | 0.951 | 0.089 | 1.064 | 0.378 |
| R71V | 0.944 | 0.047 | 1.180 | 0.097 |
| R71C | 1.096 | 0.054 | 0.986 | 0.128 |
| R71A | 1.159 | 0.068 | 0.901 | 0.034 |
| R71T | 1.271 | 0.126 | 0.919 | 0.276 |
| R71Y | 0.938 | 0.101 | 1.290 | 0.195 |
| R71G | 1.199 | 0.078 | 1.230 | 0.108 |
| R71S | 1.265 | 0.065 | 0.919 | 0.128 |

| Variant ID | Average WT-SecM Score n = 3 | WT-SecM SD | Average P166A-SecM Score n = 3 | P166A-SecM SD |
|---|---|---|---|---|
| R71H | 1.034 | 0.048 | 0.997 | 0.074 |
| R71P | 0.824 | 0.025 | 1.008 | 0.100 |
| R71Q | 1.198 | 0.121 | 0.832 | 0.177 |
| R71E | 1.159 | 0.059 | 1.097 | 0.275 |
| R71K | 1.192 | 0.153 | 0.905 | 0.089 |
| V72I | 0.993 | 0.012 | 1.009 | 0.024 |
| V72L | 0.917 | 0.035 | 1.068 | 0.108 |
| V72F | 1.035 | 0.042 | 0.870 | 0.059 |
| V72C | 1.178 | 0.087 | 0.892 | 0.085 |
| V72A | 1.062 | 0.092 | 0.937 | 0.121 |
| V72T | 0.699 | 0.067 | 1.123 | 0.323 |
| V72Y | 0.865 | 0.074 | 0.790 | 0.189 |
| V72G | 0.863 | 0.017 | 0.881 | 0.157 |
| V72S | 0.642 | 0.051 | 1.060 | 0.049 |
| V72N | 0.581 | 0.049 | 1.104 | 0.362 |
| V72H | 0.553 | 0.029 | 1.161 | 0.347 |
| V72P | 0.757 | 0.035 | 1.105 | 0.095 |
| V72Q | 0.636 | 0.068 | 1.226 | 0.182 |
| V72R | 0.391 | 0.030 | 0.654 | 0.103 |
| V72E | 0.508 | 0.027 | 0.958 | 0.273 |
| V72K | 0.409 | 0.017 | 0.754 | 0.140 |
| V72D | 0.464 | 0.030 | 1.117 | 0.129 |
| T73L | 1.218 | 0.030 | 0.779 | 0.139 |
| T73F | 1.205 | 0.093 | 1.527 | 0.150 |
| T73V | 1.351 | 0.064 | 1.267 | 0.195 |
| T73C | 1.254 | 0.103 | 0.795 | 0.081 |
| T73M | 1.068 | 0.053 | 1.048 | 0.021 |
| T73A | 1.068 | 0.059 | 1.080 | 0.079 |
| T73W | 1.063 | 0.078 | 1.201 | 0.287 |
| T73Y | 1.235 | 0.075 | 0.693 | 0.103 |
| T73G | 0.631 | 0.028 | 0.872 | 0.026 |
| T73S | 1.109 | 0.035 | 1.048 | 0.169 |
| T73N | 0.993 | 0.083 | 0.998 | 0.212 |
| T73H | 1.259 | 0.033 | 1.170 | 0.110 |
| T73P | 0.526 | 0.015 | 0.902 | 0.129 |
| T73Q | 1.148 | 0.095 | 1.094 | 0.192 |
| T73R | 0.982 | 0.046 | 0.878 | 0.127 |
| T73E | 1.244 | 0.014 | 0.752 | 0.094 |
| T73K | 1.107 | 0.101 | 0.879 | 0.082 |
| T73D | 1.033 | 0.107 | 1.068 | 0.187 |
| W74I | 0.684 | 0.048 | 0.816 | 0.245 |
| W74L | 0.666 | 0.016 | 0.969 | 0.111 |
| W74F | 1.172 | 0.057 | 0.794 | 0.041 |

| Variant ID | Average WT-SecM Score n = 3 | WT-SecM SD | Average P166A-SecM Score n = 3 | P166A-SecM SD |
|---|---|---|---|---|
| W74V | 0.699 | 0.031 | 1.098 | 0.134 |
| W74C | 0.669 | 0.018 | 1.030 | 0.100 |
| W74A | 0.473 | 0.011 | 0.979 | 0.133 |
| W74T | 0.659 | 0.014 | 0.989 | 0.190 |
| W74Y | 1.228 | 0.059 | 1.209 | 0.485 |
| W74G | 0.507 | 0.002 | 0.661 | 0.125 |
| W74S | 0.519 | 0.025 | 0.913 | 0.141 |
| W74N | 0.501 | 0.029 | 1.014 | 0.223 |
| W74H | 0.704 | 0.005 | 1.350 | 0.283 |
| W74P | 0.392 | 0.040 | 0.785 | 0.131 |
| W74Q | 0.510 | 0.014 | 1.166 | 0.087 |
| W74R | 0.657 | 0.034 | 0.989 | 0.037 |
| W74E | 0.416 | 0.042 | 0.734 | 0.102 |
| W74K | 0.411 | 0.021 | 1.006 | 0.104 |
| W74D | 0.372 | 0.043 | 0.672 | 0.068 |
| V75I | 0.662 | 0.053 | 0.829 | 0.130 |
| V75L | 0.562 | 0.015 | 0.899 | 0.209 |
| V75F | 0.466 | 0.040 | 0.894 | 0.149 |
| V75C | 1.322 | 0.055 | 1.656 | 0.188 |
| V75M | 0.865 | 0.044 | 1.007 | 0.024 |
| V75A | 1.127 | 0.062 | 1.011 | 0.011 |
| V75T | 1.150 | 0.094 | 0.915 | 0.078 |
| V75Y | 0.458 | 0.043 | 1.026 | 0.273 |
| V75G | 0.545 | 0.040 | 0.920 | 0.159 |
| V75S | 1.011 | 0.067 | 1.065 | 0.145 |
| V75N | 0.414 | 0.035 | 0.764 | 0.188 |
| V75H | 0.517 | 0.035 | 1.220 | 0.100 |
| V75P | 0.381 | 0.034 | 0.619 | 0.072 |
| V75Q | 0.526 | 0.033 | 1.027 | 0.144 |
| V75R | 0.673 | 0.019 | 0.918 | 0.248 |
| V75K | 0.618 | 0.021 | 0.797 | 0.120 |
| V75D | 0.343 | 0.029 | 0.662 | 0.113 |
| K76I | 0.948 | 0.047 | 1.049 | 0.135 |
| K76L | 1.090 | 0.064 | 1.016 | 0.036 |
| K76F | 0.995 | 0.102 | 1.129 | 0.160 |
| K76V | 1.104 | 0.053 | 1.053 | 0.092 |
| K76C | 1.246 | 0.095 | 0.814 | 0.132 |
| K76A | 1.288 | 0.042 | 0.995 | 0.094 |
| K76T | 1.265 | 0.051 | 0.933 | 0.082 |
| K76Y | 1.147 | 0.078 | 0.904 | 0.069 |
| K76G | 1.237 | 0.081 | 1.284 | 0.062 |
| K76S | 1.381 | 0.062 | 1.057 | 0.116 |
| K76N | 1.206 | 0.090 | 0.844 | 0.051 |

| Variant ID | Average WT-SecM Score n = 3 | WT-SecM SD | Average P166A-SecM Score n = 3 | P166A-SecM SD |
|---|---|---|---|---|
| K76H | 1.244 | 0.070 | 1.247 | 0.257 |
| K76P | 1.017 | 0.084 | 1.067 | 0.310 |
| K76R | 1.210 | 0.079 | 0.917 | 0.127 |
| K76E | 1.134 | 0.083 | 0.943 | 0.032 |
| K76D | 1.165 | 0.047 | 0.704 | 0.060 |
| S77I | 0.549 | 0.040 | 0.972 | 0.180 |
| S77L | 0.714 | 0.028 | 0.901 | 0.053 |
| S77F | 0.770 | 0.042 | 1.030 | 0.084 |
| S77V | 0.632 | 0.030 | 1.222 | 0.129 |
| S77C | 1.164 | 0.052 | 0.837 | 0.109 |
| S77M | 0.715 | 0.033 | 1.168 | 0.193 |
| S77A | 0.934 | 0.051 | 1.004 | 0.092 |
| S77W | 0.610 | 0.028 | 0.974 | 0.303 |
| S77T | 1.308 | 0.069 | 1.021 | 0.125 |
| S77Y | 0.835 | 0.054 | 0.800 | 0.035 |
| S77G | 1.167 | 0.082 | 0.918 | 0.100 |
| S77N | 1.259 | 0.091 | 1.046 | 0.168 |
| S77H | 1.053 | 0.071 | 1.023 | 0.316 |
| S77P | 0.453 | 0.022 | 0.642 | 0.071 |
| S77Q | 0.892 | 0.036 | 1.039 | 0.264 |
| S77R | 0.898 | 0.056 | 1.095 | 0.088 |
| S77E | 1.073 | 0.066 | 1.194 | 0.274 |
| S77D | 1.249 | 0.100 | 0.792 | 0.168 |
| V78I | 1.229 | 0.092 | 0.975 | 0.135 |
| V78L | 1.006 | 0.063 | 0.966 | 0.142 |
| V78F | 0.797 | 0.042 | 1.108 | 0.206 |
| V78C | 1.244 | 0.089 | 1.267 | 0.074 |
| V78M | 0.982 | 0.028 | 0.968 | 0.034 |
| V78A | 1.130 | 0.073 | 1.024 | 0.089 |
| V78W | 0.621 | 0.085 | 0.713 | 0.146 |
| V78T | 1.168 | 0.056 | 0.983 | 0.075 |
| V78Y | 0.995 | 0.032 | 0.770 | 0.195 |
| V78G | 0.819 | 0.024 | 0.861 | 0.205 |
| V78S | 1.010 | 0.065 | 0.794 | 0.198 |
| V78N | 0.712 | 0.053 | 1.096 | 0.162 |
| V78H | 0.809 | 0.026 | 0.872 | 0.196 |
| V78P | 1.250 | 0.088 | 1.024 | 0.264 |
| V78Q | 0.898 | 0.041 | 0.983 | 0.039 |
| V78R | 0.981 | 0.056 | 1.128 | 0.044 |
| V78E | 0.982 | 0.012 | 0.843 | 0.277 |
| V78K | 0.777 | 0.048 | 1.230 | 0.240 |
| V78D | 0.798 | 0.057 | 1.073 | 0.211 |
| D79I | 0.755 | 0.022 | 0.828 | 0.093 |

| Variant ID | Average WT-SecM Score n = 3 | WT-SecM SD | Average P166A-SecM Score n = 3 | P166A-SecM SD |
|---|---|---|---|---|
| D79L | 0.765 | 0.040 | 1.022 | 0.146 |
| D79F | 0.785 | 0.064 | 0.920 | 0.025 |
| D79V | 0.874 | 0.055 | 1.045 | 0.156 |
| D79C | 0.879 | 0.089 | 0.946 | 0.066 |
| D79M | 0.845 | 0.087 | 0.902 | 0.369 |
| D79A | 1.136 | 0.055 | 1.064 | 0.209 |
| D79W | 0.760 | 0.055 | 0.757 | 0.262 |
| D79T | 1.095 | 0.041 | 0.871 | 0.238 |
| D79Y | 0.871 | 0.027 | 1.166 | 0.062 |
| D79G | 1.014 | 0.089 | 1.013 | 0.011 |
| D79S | 1.124 | 0.062 | 1.065 | 0.130 |
| D79N | 1.065 | 0.006 | 0.885 | 0.042 |
| D79H | 1.050 | 0.100 | 1.082 | 0.131 |
| D79P | 1.121 | 0.069 | 1.111 | 0.065 |
| D79Q | 1.102 | 0.092 | 0.697 | 0.147 |
| D79R | 1.069 | 0.076 | 1.274 | 0.110 |
| D79E | 1.224 | 0.048 | 1.080 | 0.212 |
| D79K | 1.021 | 0.042 | 0.498 | 0.113 |
| E80I | 0.602 | 0.057 | 1.191 | 0.040 |
| E80L | 0.872 | 0.048 | 0.907 | 0.133 |
| E80F | 0.617 | 0.058 | 1.044 | 0.164 |
| E80V | 0.654 | 0.041 | 0.785 | 0.103 |
| E80C | 0.864 | 0.023 | 1.140 | 0.097 |
| E80A | 1.168 | 0.048 | 1.074 | 0.114 |
| E80W | 0.518 | 0.052 | 1.136 | 0.125 |
| E80T | 0.834 | 0.038 | 1.050 | 0.142 |
| E80Y | 0.647 | 0.031 | 0.986 | 0.101 |
| E80G | 0.990 | 0.067 | 0.965 | 0.083 |
| E80S | 0.973 | 0.033 | 0.982 | 0.132 |
| E80N | 0.836 | 0.039 | 1.114 | 0.125 |
| E80H | 1.012 | 0.082 | 1.006 | 0.064 |
| E80P | 0.394 | 0.036 | 0.534 | 0.008 |
| E80Q | 1.127 | 0.086 | 0.838 | 0.176 |
| E80R | 0.838 | 0.074 | 1.034 | 0.082 |
| E80K | 0.970 | 0.045 | 1.180 | 0.104 |
| E80D | 1.185 | 0.051 | 1.091 | 0.081 |
| A81I | 0.392 | 0.033 | 0.773 | 0.120 |
| A81L | 0.403 | 0.052 | 0.754 | 0.060 |
| A81F | 0.361 | 0.046 | 0.426 | 0.055 |
| A81V | 0.594 | 0.004 | 1.047 | 0.133 |
| A81C | 0.764 | 0.059 | 0.946 | 0.142 |
| A81W | 0.366 | 0.044 | 0.315 | 0.087 |
| A81T | 0.514 | 0.022 | 0.864 | 0.122 |

| Variant ID | Average WT-SecM Score n = 3 | WT-SecM SD | Average P166A-SecM Score n = 3 | P166A-SecM SD |
|---|---|---|---|---|
| A81Y | 0.359 | 0.035 | 0.508 | 0.073 |
| A81G | 0.911 | 0.040 | 0.955 | 0.064 |
| A81S | 0.845 | 0.046 | 1.116 | 0.088 |
| A81N | 0.352 | 0.051 | 0.283 | 0.058 |
| A81H | 0.339 | 0.041 | 0.406 | 0.029 |
| A81P | 0.346 | 0.037 | 0.399 | 0.046 |
| A81Q | 0.343 | 0.047 | 0.463 | 0.023 |
| A81R | 0.336 | 0.031 | 0.400 | 0.034 |
| A81E | 0.355 | 0.043 | 0.434 | 0.127 |
| A81K | 0.347 | 0.038 | 0.339 | 0.048 |
| A81D | 0.372 | 0.047 | 0.481 | 0.101 |
| I82L | 1.163 | 0.089 | 1.054 | 0.082 |
| I82F | 0.720 | 0.038 | 0.853 | 0.109 |
| I82V | 1.112 | 0.050 | 0.999 | 0.136 |
| I82C | 0.944 | 0.059 | 1.091 | 0.125 |
| I82M | 1.066 | 0.092 | 0.939 | 0.135 |
| I82A | 0.568 | 0.029 | 1.091 | 0.145 |
| I82W | 0.503 | 0.054 | 1.053 | 0.078 |
| I82T | 0.804 | 0.057 | 1.026 | 0.145 |
| I82Y | 0.695 | 0.062 | 1.079 | 0.292 |
| I82G | 0.410 | 0.031 | 0.734 | 0.032 |
| I82S | 0.510 | 0.016 | 0.996 | 0.175 |
| I82N | 0.461 | 0.045 | 0.705 | 0.099 |
| I82H | 0.531 | 0.033 | 1.089 | 0.134 |
| I82P | 0.366 | 0.048 | 0.407 | 0.094 |
| I82Q | 0.669 | 0.029 | 0.818 | 0.254 |
| I82R | 0.571 | 0.036 | 0.964 | 0.138 |
| I82E | 0.784 | 0.077 | 0.864 | 0.105 |
| I82K | 0.578 | 0.031 | 1.240 | 0.125 |
| I82D | 0.413 | 0.051 | 0.642 | 0.056 |
| A83I | 1.257 | 0.090 | 0.926 | 0.207 |
| A83L | 1.302 | 0.084 | 1.083 | 0.137 |
| A83F | 1.185 | 0.080 | 1.282 | 0.211 |
| A83V | 1.159 | 0.058 | 0.956 | 0.081 |
| A83C | 1.206 | 0.084 | 1.181 | 0.264 |
| A83M | 1.353 | 0.019 | 0.845 | 0.091 |
| A83W | 1.145 | 0.079 | 0.697 | 0.148 |
| A83T | 1.241 | 0.021 | 0.908 | 0.085 |
| A83Y | 1.250 | 0.104 | 1.016 | 0.196 |
| A83G | 1.161 | 0.098 | 0.979 | 0.148 |
| A83S | 1.263 | 0.045 | 0.950 | 0.102 |
| A83N | 1.273 | 0.067 | 1.151 | 0.063 |
| A83H | 1.254 | 0.090 | 1.119 | 0.229 |

| Variant ID | Average WT-SecM Score n = 3 | WT-SecM SD | Average P166A-SecM Score n = 3 | P166A-SecM SD |
|---|---|---|---|---|
| A83P | 0.483 | 0.027 | 0.969 | 0.211 |
| A83Q | 1.346 | 0.072 | 0.868 | 0.147 |
| A83R | 1.278 | 0.082 | 1.010 | 0.149 |
| A83E | 1.245 | 0.037 | 1.194 | 0.139 |
| A83K | 1.248 | 0.064 | 1.316 | 0.366 |
| A83D | 1.284 | 0.044 | 0.869 | 0.262 |
| A84I | 1.067 | 0.075 | 1.075 | 0.020 |
| A84L | 0.962 | 0.042 | 0.915 | 0.017 |
| A84F | 0.903 | 0.054 | 0.897 | 0.030 |
| A84V | 1.195 | 0.068 | 1.002 | 0.024 |
| A84C | 1.173 | 0.096 | 1.023 | 0.054 |
| A84M | 1.175 | 0.029 | 1.147 | 0.230 |
| A84T | 1.027 | 0.062 | 1.137 | 0.079 |
| A84Y | 0.877 | 0.080 | 1.214 | 0.284 |
| A84G | 0.706 | 0.062 | 1.107 | 0.051 |
| A84S | 1.154 | 0.058 | 0.980 | 0.067 |
| A84N | 0.950 | 0.049 | 1.048 | 0.143 |
| A84H | 1.042 | 0.082 | 1.240 | 0.042 |
| A84P | 0.357 | 0.035 | 0.449 | 0.042 |
| A84Q | 1.173 | 0.057 | 1.227 | 0.204 |
| A84R | 0.985 | 0.068 | 1.021 | 0.200 |
| A84E | 1.229 | 0.097 | 1.021 | 0.108 |
| A84K | 1.064 | 0.111 | 0.879 | 0.164 |
| A84D | 0.693 | 0.036 | 1.046 | 0.059 |
| A85I | 0.707 | 0.017 | 0.921 | 0.257 |
| A85L | 0.606 | 0.030 | 0.816 | 0.154 |
| A85F | 0.505 | 0.011 | 1.108 | 0.150 |
| A85V | 0.823 | 0.048 | 0.757 | 0.038 |
| A85C | 1.235 | 0.075 | 1.123 | 0.173 |
| A85M | 0.555 | 0.040 | 0.629 | 0.119 |
| A85W | 0.492 | 0.027 | 0.968 | 0.006 |
| A85T | 0.762 | 0.036 | 0.951 | 0.046 |
| A85Y | 0.456 | 0.024 | 1.000 | 0.127 |
| A85G | 1.038 | 0.050 | 1.014 | 0.093 |
| A85S | 1.032 | 0.046 | 0.952 | 0.103 |
| A85N | 0.439 | 0.053 | 0.708 | 0.157 |
| A85H | 0.425 | 0.030 | 0.899 | 0.089 |
| A85P | 0.504 | 0.031 | 1.060 | 0.087 |
| A85Q | 0.416 | 0.030 | 0.670 | 0.095 |
| A85R | 0.388 | 0.044 | 0.530 | 0.071 |
| A85E | 0.468 | 0.057 | 0.721 | 0.048 |
| A85K | 0.399 | 0.035 | 0.644 | 0.058 |
| A85D | 0.365 | 0.019 | 0.601 | 0.055 |

| Variant ID | Average WT-SecM Score n = 3 | WT-SecM SD | Average P166A-SecM Score n = 3 | P166A-SecM SD |
|---|---|---|---|---|
| G86I | 0.547 | 0.026 | 0.913 | 0.096 |
| G86L | 0.606 | 0.029 | 1.065 | 0.122 |
| G86F | 0.583 | 0.046 | 1.377 | 0.296 |
| G86V | 0.580 | 0.009 | 1.219 | 0.168 |
| G86C | 0.748 | 0.068 | 1.001 | 0.035 |
| G86M | 0.716 | 0.068 | 1.097 | 0.132 |
| G86A | 0.865 | 0.044 | 1.004 | 0.064 |
| G86W | 0.561 | 0.060 | 0.999 | 0.298 |
| G86T | 0.652 | 0.038 | 0.999 | 0.171 |
| G86Y | 0.587 | 0.070 | 0.873 | 0.033 |
| G86S | 0.851 | 0.061 | 1.036 | 0.045 |
| G86N | 0.734 | 0.028 | 0.915 | 0.196 |
| G86H | 0.718 | 0.006 | 1.124 | 0.038 |
| G86P | 0.650 | 0.025 | 1.121 | 0.052 |
| G86Q | 0.779 | 0.076 | 0.816 | 0.105 |
| G86R | 0.741 | 0.041 | 1.105 | 0.021 |
| G86E | 0.908 | 0.088 | 0.825 | 0.197 |
| G86K | 0.751 | 0.058 | 1.292 | 0.197 |
| G86D | 0.913 | 0.017 | 0.920 | 0.160 |
| D87I | 1.051 | 0.053 | 1.197 | 0.069 |
| D87L | 1.042 | 0.051 | 0.870 | 0.051 |
| D87F | 0.954 | 0.061 | 1.233 | 0.244 |
| D87V | 1.111 | 0.030 | 1.038 | 0.151 |
| D87C | 1.183 | 0.084 | 1.095 | 0.056 |
| D87M | 1.014 | 0.094 | 0.618 | 0.104 |
| D87A | 1.182 | 0.097 | 1.007 | 0.086 |
| D87W | 0.782 | 0.063 | 1.171 | 0.181 |
| D87T | 1.116 | 0.060 | 0.915 | 0.140 |
| D87Y | 0.976 | 0.085 | 0.999 | 0.094 |
| D87G | 1.014 | 0.054 | 1.010 | 0.141 |
| D87S | 1.213 | 0.074 | 1.077 | 0.204 |
| D87N | 1.221 | 0.044 | 0.873 | 0.023 |
| D87H | 1.190 | 0.021 | 1.057 | 0.126 |
| D87P | 1.077 | 0.088 | 1.142 | 0.203 |
| D87Q | 1.162 | 0.043 | 1.058 | 0.178 |
| D87R | 1.152 | 0.079 | 1.009 | 0.169 |
| D87E | 1.164 | 0.074 | 1.101 | 0.224 |
| D87K | 1.185 | 0.082 | 0.962 | 0.105 |
| V88I | 0.896 | 0.018 | 0.929 | 0.083 |
| V88L | 0.872 | 0.076 | 0.977 | 0.122 |
| V88F | 1.081 | 0.056 | 0.854 | 0.102 |
| V88C | 1.329 | 0.073 | 0.994 | 0.125 |
| V88M | 1.111 | 0.118 | 0.721 | 0.255 |

| Variant ID | Average WT-SecM Score n = 3 | WT-SecM SD | Average P166A-SecM Score n = 3 | P166A-SecM SD |
|---|---|---|---|---|
| V88A | 1.239 | 0.069 | 1.079 | 0.070 |
| V88W | 0.830 | 0.060 | 1.021 | 0.353 |
| V88T | 1.173 | 0.082 | 0.815 | 0.087 |
| V88Y | 1.074 | 0.071 | 0.960 | 0.088 |
| V88G | 1.127 | 0.034 | 0.921 | 0.080 |
| V88S | 1.308 | 0.089 | 0.942 | 0.066 |
| V88N | 1.066 | 0.119 | 0.994 | 0.151 |
| V88H | 1.217 | 0.076 | 1.012 | 0.073 |
| V88P | 0.915 | 0.052 | 1.095 | 0.181 |
| V88Q | 1.144 | 0.021 | 0.852 | 0.065 |
| V88R | 0.941 | 0.104 | 1.251 | 0.202 |
| V88E | 1.376 | 0.065 | 0.818 | 0.157 |
| V88K | 0.946 | 0.047 | 0.766 | 0.120 |
| V88D | 1.190 | 0.090 | 1.069 | 0.270 |
| P89I | 0.995 | 0.072 | 0.795 | 0.093 |
| P89L | 0.978 | 0.053 | 0.971 | 0.134 |
| P89F | 0.769 | 0.035 | 0.820 | 0.189 |
| P89V | 1.094 | 0.063 | 0.901 | 0.149 |
| P89C | 1.054 | 0.091 | 0.972 | 0.140 |
| P89A | 1.203 | 0.059 | 0.755 | 0.164 |
| P89W | 0.772 | 0.097 | 1.189 | 0.186 |
| P89T | 1.095 | 0.080 | 0.889 | 0.107 |
| P89Y | 0.924 | 0.142 | 0.886 | 0.116 |
| P89G | 1.139 | 0.064 | 0.921 | 0.247 |
| P89S | 1.104 | 0.062 | 0.947 | 0.027 |
| P89N | 1.142 | 0.113 | 1.036 | 0.333 |
| P89H | 1.101 | 0.089 | 1.098 | 0.155 |
| P89Q | 1.167 | 0.132 | 1.004 | 0.039 |
| P89R | 1.095 | 0.079 | 1.146 | 0.206 |
| P89E | 1.345 | 0.118 | 1.097 | 0.255 |
| P89K | 1.044 | 0.074 | 0.926 | 0.030 |
| P89D | 1.315 | 0.113 | 1.090 | 0.155 |
| E90I | 0.419 | 0.037 | 0.691 | 0.092 |
| E90L | 0.420 | 0.033 | 0.737 | 0.078 |
| E90F | 0.400 | 0.038 | 0.714 | 0.059 |
| E90V | 0.439 | 0.039 | 0.772 | 0.021 |
| E90C | 0.447 | 0.051 | 0.923 | 0.164 |
| E90M | 0.466 | 0.061 | 0.954 | 0.234 |
| E90A | 0.446 | 0.021 | 0.704 | 0.092 |
| E90W | 0.427 | 0.031 | 0.640 | 0.139 |
| E90T | 0.423 | 0.027 | 0.705 | 0.109 |
| E90Y | 0.411 | 0.040 | 0.632 | 0.090 |
| E90G | 0.488 | 0.042 | 0.753 | 0.069 |

| Variant ID | Average WT-SecM Score n = 3 | WT-SecM SD | Average P166A-SecM Score n = 3 | P166A-SecM SD |
|---|---|---|---|---|
| E90S | 0.437 | 0.027 | 0.696 | 0.091 |
| E90N | 0.520 | 0.014 | 0.855 | 0.093 |
| E90H | 0.497 | 0.037 | 0.866 | 0.129 |
| E90P | 0.366 | 0.031 | 0.607 | 0.054 |
| E90Q | 0.561 | 0.030 | 1.036 | 0.070 |
| E90R | 0.385 | 0.031 | 0.585 | 0.093 |
| E90K | 0.466 | 0.031 | 0.617 | 0.132 |
| E90D | 0.711 | 0.033 | 0.835 | 0.103 |
| I91L | 0.951 | 0.064 | 0.985 | 0.060 |
| I91F | 0.601 | 0.031 | 0.851 | 0.179 |
| I91V | 1.153 | 0.100 | 0.840 | 0.126 |
| I91C | 0.960 | 0.067 | 1.144 | 0.058 |
| I91M | 0.978 | 0.033 | 0.974 | 0.111 |
| I91A | 0.923 | 0.093 | 1.087 | 0.103 |
| I91W | 0.415 | 0.073 | 0.729 | 0.088 |
| I91T | 0.986 | 0.069 | 0.952 | 0.054 |
| I91Y | 0.528 | 0.048 | 0.884 | 0.186 |
| I91G | 0.468 | 0.040 | 0.655 | 0.054 |
| I91S | 0.638 | 0.038 | 1.027 | 0.121 |
| I91N | 0.386 | 0.029 | 0.566 | 0.075 |
| I91H | 0.407 | 0.029 | 0.840 | 0.088 |
| I91P | 0.387 | 0.037 | 0.455 | 0.037 |
| I91Q | 0.450 | 0.044 | 0.894 | 0.078 |
| I91R | 0.418 | 0.040 | 0.791 | 0.017 |
| I91E | 0.357 | 0.035 | 0.411 | 0.008 |
| I91K | 0.572 | 0.040 | 0.762 | 0.137 |
| I91D | 0.346 | 0.038 | 0.390 | 0.085 |
| M92I | 1.067 | 0.035 | 0.960 | 0.041 |
| M92L | 1.032 | 0.046 | 1.013 | 0.119 |
| M92F | 1.259 | 0.095 | 1.157 | 0.209 |
| M92V | 1.140 | 0.069 | 0.991 | 0.116 |
| M92C | 1.146 | 0.058 | 1.121 | 0.051 |
| M92A | 0.713 | 0.044 | 1.109 | 0.148 |
| M92W | 0.622 | 0.046 | 0.905 | 0.128 |
| M92T | 0.810 | 0.039 | 0.982 | 0.113 |
| M92Y | 0.836 | 0.059 | 1.016 | 0.173 |
| M92G | 0.396 | 0.044 | 0.516 | 0.071 |
| M92S | 0.552 | 0.041 | 0.955 | 0.164 |
| M92N | 0.436 | 0.033 | 0.767 | 0.149 |
| M92H | 0.526 | 0.032 | 0.897 | 0.039 |
| M92P | 0.380 | 0.017 | 0.464 | 0.033 |
| M92Q | 0.542 | 0.045 | 1.083 | 0.153 |
| M92R | 0.723 | 0.072 | 0.701 | 0.072 |

| Variant ID | Average WT-SecM Score n = 3 | WT-SecM SD | Average P166A-SecM Score n = 3 | P166A-SecM SD |
|---|---|---|---|---|
| M92E | 0.373 | 0.040 | 0.551 | 0.110 |
| M92K | 0.399 | 0.028 | 0.625 | 0.092 |
| M92D | 0.386 | 0.048 | 0.440 | 0.020 |
| V93I | 0.998 | 0.074 | 1.049 | 0.158 |
| V93L | 0.527 | 0.008 | 0.832 | 0.069 |
| V93F | 0.407 | 0.028 | 0.705 | 0.057 |
| V93C | 0.883 | 0.102 | 0.841 | 0.095 |
| V93M | 0.656 | 0.030 | 0.902 | 0.030 |
| V93A | 0.778 | 0.029 | 0.864 | 0.134 |
| V93T | 0.479 | 0.032 | 0.935 | 0.069 |
| V93Y | 0.347 | 0.025 | 0.673 | 0.093 |
| V93G | 0.739 | 0.069 | 0.763 | 0.060 |
| V93S | 0.397 | 0.036 | 0.667 | 0.065 |
| V93N | 0.349 | 0.047 | 0.458 | 0.052 |
| V93H | 0.349 | 0.050 | 0.412 | 0.111 |
| V93P | 0.343 | 0.040 | 0.383 | 0.031 |
| V93Q | 0.340 | 0.028 | 0.497 | 0.143 |
| V93R | 0.344 | 0.028 | 0.436 | 0.082 |
| V93E | 0.356 | 0.034 | 0.393 | 0.122 |
| V93K | 0.353 | 0.047 | 0.385 | 0.032 |
| V93D | 0.335 | 0.026 | 0.442 | 0.112 |
| I94L | 1.121 | 0.064 | 0.959 | 0.084 |
| I94F | 1.135 | 0.111 | 0.775 | 0.210 |
| I94V | 1.124 | 0.091 | 0.948 | 0.066 |
| I94C | 1.039 | 0.118 | 1.077 | 0.165 |
| I94M | 1.005 | 0.083 | 0.924 | 0.128 |
| I94A | 0.957 | 0.087 | 1.136 | 0.161 |
| I94T | 0.717 | 0.043 | 0.866 | 0.089 |
| I94Y | 0.662 | 0.068 | 1.064 | 0.158 |
| I94G | 0.520 | 0.057 | 0.990 | 0.098 |
| I94S | 0.621 | 0.038 | 0.989 | 0.059 |
| I94N | 0.536 | 0.002 | 0.898 | 0.161 |
| I94H | 0.727 | 0.044 | 1.046 | 0.301 |
| I94P | 0.350 | 0.045 | 0.520 | 0.021 |
| I94Q | 0.401 | 0.036 | 0.683 | 0.114 |
| I94R | 0.430 | 0.051 | 0.712 | 0.049 |
| I94E | 0.360 | 0.038 | 0.527 | 0.062 |
| I94K | 0.405 | 0.039 | 0.779 | 0.144 |
| I94D | 0.355 | 0.047 | 0.432 | 0.075 |
| G95I | 0.453 | 0.032 | 0.851 | 0.094 |
| G95L | 0.449 | 0.040 | 0.782 | 0.051 |
| G95F | 0.421 | 0.034 | 0.787 | 0.185 |
| G95V | 0.606 | 0.029 | 0.909 | 0.089 |

| Variant ID | Average WT-SecM Score n = 3 | WT-SecM SD | Average P166A-SecM Score n = 3 | P166A-SecM SD |
|---|---|---|---|---|
| G95C | 0.792 | 0.062 | 0.897 | 0.059 |
| G95A | 0.584 | 0.065 | 0.773 | 0.105 |
| G95W | 0.394 | 0.046 | 0.747 | 0.047 |
| G95T | 0.418 | 0.059 | 0.792 | 0.061 |
| G95Y | 0.447 | 0.050 | 0.682 | 0.049 |
| G95S | 0.574 | 0.026 | 0.665 | 0.046 |
| G95N | 0.372 | 0.032 | 0.605 | 0.163 |
| G95H | 0.387 | 0.043 | 0.589 | 0.092 |
| G95P | 0.388 | 0.060 | 0.582 | 0.083 |
| G95Q | 0.370 | 0.034 | 0.417 | 0.084 |
| G95R | 0.424 | 0.017 | 0.568 | 0.064 |
| G95E | 0.424 | 0.011 | 0.601 | 0.151 |
| G95K | 0.355 | 0.015 | 0.363 | 0.100 |
| G95D | 0.632 | 0.034 | 0.747 | 0.051 |
| G96I | 0.559 | 0.045 | 0.941 | 0.192 |
| G96L | 0.570 | 0.029 | 0.922 | 0.105 |
| G96F | 0.715 | 0.032 | 1.018 | 0.089 |
| G96V | 0.704 | 0.057 | 0.884 | 0.103 |
| G96C | 0.937 | 0.057 | 0.998 | 0.099 |
| G96A | 0.920 | 0.037 | 0.927 | 0.041 |
| G96W | 0.619 | 0.031 | 0.666 | 0.063 |
| G96T | 0.784 | 0.042 | 0.848 | 0.085 |
| G96Y | 0.641 | 0.052 | 0.777 | 0.104 |
| G96S | 1.030 | 0.036 | 0.891 | 0.096 |
| G96N | 0.949 | 0.065 | 0.930 | 0.267 |
| G96H | 0.905 | 0.060 | 1.122 | 0.241 |
| G96P | 1.066 | 0.077 | 0.832 | 0.193 |
| G96Q | 0.716 | 0.054 | 0.931 | 0.193 |
| G96R | 0.664 | 0.032 | 0.832 | 0.035 |
| G96K | 0.695 | 0.034 | 0.720 | 0.124 |
| G96D | 1.129 | 0.058 | 0.989 | 0.096 |
| G97I | 1.242 | 0.043 | 1.040 | 0.034 |
| G97L | 1.243 | 0.072 | 1.008 | 0.031 |
| G97F | 1.181 | 0.104 | 0.735 | 0.086 |
| G97V | 1.202 | 0.028 | 0.659 | 0.110 |
| G97C | 1.182 | 0.054 | 0.913 | 0.193 |
| G97A | 1.285 | 0.023 | 1.090 | 0.054 |
| G97T | 1.231 | 0.042 | 0.913 | 0.067 |
| G97Y | 1.158 | 0.041 | 0.855 | 0.074 |
| G97S | 1.195 | 0.024 | 0.933 | 0.055 |
| G97N | 1.277 | 0.097 | 1.074 | 0.299 |
| G97H | 1.302 | 0.155 | 0.679 | 0.237 |
| G97P | 1.312 | 0.040 | 0.883 | 0.096 |

| Variant ID | Average WT-SecM Score n = 3 | WT-SecM SD | Average P166A-SecM Score n = 3 | P166A-SecM SD |
|---|---|---|---|---|
| G97Q | 1.350 | 0.113 | 1.024 | 0.202 |
| G97R | 1.288 | 0.025 | 0.997 | 0.015 |
| G97K | 1.360 | 0.063 | 0.718 | 0.145 |
| G97D | 1.046 | 0.050 | 1.084 | 0.027 |
| R98I | 1.301 | 0.097 | 0.820 | 0.066 |
| R98L | 1.210 | 0.077 | 0.922 | 0.070 |
| R98V | 1.299 | 0.050 | 1.291 | 0.169 |
| R98C | 1.102 | 0.042 | 1.011 | 0.059 |
| R98A | 1.381 | 0.070 | 1.136 | 0.142 |
| R98T | 1.344 | 0.068 | 1.016 | 0.297 |
| R98Y | 1.307 | 0.130 | 1.255 | 0.154 |
| R98G | 1.258 | 0.045 | 1.154 | 0.178 |
| R98S | 1.300 | 0.069 | 0.990 | 0.141 |
| R98H | 1.132 | 0.020 | 1.019 | 0.061 |
| R98P | 1.224 | 0.090 | 0.727 | 0.133 |
| R98K | 1.350 | 0.115 | 0.878 | 0.125 |
| V99I | 1.176 | 0.077 | 0.931 | 0.147 |
| V99L | 1.284 | 0.060 | 1.054 | 0.135 |
| V99F | 0.457 | 0.025 | 0.575 | 0.081 |
| V99C | 1.109 | 0.047 | 0.826 | 0.056 |
| V99A | 0.957 | 0.063 | 0.991 | 0.091 |
| V99W | 0.527 | 0.054 | 1.002 | 0.211 |
| V99T | 1.240 | 0.082 | 0.900 | 0.108 |
| V99Y | 0.496 | 0.063 | 0.833 | 0.134 |
| V99G | 0.654 | 0.013 | 0.776 | 0.052 |
| V99S | 0.746 | 0.047 | 1.064 | 0.182 |
| V99H | 0.464 | 0.046 | 0.778 | 0.013 |
| V99P | 0.516 | 0.037 | 0.897 | 0.114 |
| V99Q | 0.678 | 0.043 | 1.085 | 0.216 |
| V99R | 0.419 | 0.017 | 0.557 | 0.056 |
| V99E | 1.052 | 0.056 | 1.117 | 0.155 |
| V99K | 0.397 | 0.058 | 0.522 | 0.067 |
| V99D | 0.605 | 0.018 | 0.718 | 0.164 |
| Y100I | 0.802 | 0.049 | 0.716 | 0.072 |
| Y100L | 1.171 | 0.061 | 1.159 | 0.103 |
| Y100V | 0.661 | 0.050 | 0.846 | 0.071 |
| Y100C | 0.884 | 0.084 | 0.910 | 0.067 |
| Y100A | 0.829 | 0.040 | 0.962 | 0.246 |
| Y100W | 1.225 | 0.070 | 0.676 | 0.142 |
| Y100T | 0.874 | 0.042 | 0.855 | 0.113 |
| Y100G | 0.665 | 0.044 | 0.849 | 0.064 |
| Y100S | 0.685 | 0.030 | 0.923 | 0.142 |
| Y100N | 0.664 | 0.032 | 0.720 | 0.188 |

| Variant ID | Average WT-SecM Score n = 3 | WT-SecM SD | Average P166A-SecM Score n = 3 | P166A-SecM SD |
|---|---|---|---|---|
| Y100H | 0.940 | 0.075 | 0.952 | 0.144 |
| Y100P | 0.427 | 0.037 | 0.667 | 0.030 |
| Y100Q | 0.633 | 0.021 | 0.563 | 0.362 |
| Y100R | 0.477 | 0.047 | 0.700 | 0.184 |
| Y100E | 0.493 | 0.027 | 0.537 | 0.125 |
| Y100K | 0.382 | 0.048 | 0.573 | 0.096 |
| Y100D | 0.582 | 0.055 | 0.737 | 0.099 |
| E101I | 0.694 | 0.079 | 0.998 | 0.498 |
| E101L | 0.777 | 0.054 | 0.977 | 0.235 |
| E101V | 0.786 | 0.044 | 1.004 | 0.121 |
| E101C | 0.863 | 0.042 | 1.245 | 0.266 |
| E101A | 1.058 | 0.002 | 1.065 | 0.073 |
| E101Y | 0.963 | 0.117 | 1.036 | 0.097 |
| E101G | 0.982 | 0.055 | 0.981 | 0.027 |
| E101S | 1.207 | 0.053 | 1.029 | 0.207 |
| E101H | 1.108 | 0.074 | 1.161 | 0.190 |
| E101P | 0.674 | 0.024 | 1.109 | 0.064 |
| E101Q | 1.161 | 0.051 | 1.046 | 0.197 |
| E101R | 1.175 | 0.064 | 1.194 | 0.162 |
| E101K | 1.026 | 0.021 | 1.069 | 0.072 |
| E101D | 1.019 | 0.052 | 0.944 | 0.081 |
| Q102I | 0.879 | 0.082 | 0.669 | 0.067 |
| Q102L | 1.213 | 0.079 | 1.033 | 0.181 |
| Q102F | 0.769 | 0.069 | 0.961 | 0.064 |
| Q102V | 0.892 | 0.088 | 0.953 | 0.140 |
| Q102C | 1.049 | 0.087 | 1.223 | 0.279 |
| Q102M | 1.238 | 0.128 | 1.135 | 0.254 |
| Q102A | 1.370 | 0.067 | 1.163 | 0.193 |
| Q102W | 1.155 | 0.013 | 0.903 | 0.142 |
| Q102T | 1.207 | 0.069 | 0.953 | 0.079 |
| Q102Y | 1.094 | 0.084 | 1.125 | 0.135 |
| Q102G | 0.968 | 0.072 | 0.849 | 0.166 |
| Q102S | 1.122 | 0.063 | 0.809 | 0.147 |
| Q102N | 1.106 | 0.054 | 1.026 | 0.215 |
| Q102H | 1.121 | 0.094 | 1.035 | 0.072 |
| Q102P | 0.675 | 0.006 | 0.906 | 0.087 |
| Q102R | 1.054 | 0.059 | 0.921 | 0.155 |
| Q102E | 1.301 | 0.065 | 1.016 | 0.030 |
| Q102K | 1.067 | 0.057 | 0.992 | 0.096 |
| Q102D | 1.297 | 0.048 | 1.114 | 0.348 |
| F103I | 0.678 | 0.026 | 0.948 | 0.081 |
| F103L | 0.902 | 0.038 | 0.981 | 0.114 |
| F103V | 0.699 | 0.015 | 1.088 | 0.079 |

| Variant ID | Average WT-SecM Score n = 3 | WT-SecM SD | Average P166A-SecM Score n = 3 | P166A-SecM SD |
|---|---|---|---|---|
| F103C | 0.972 | 0.034 | 0.980 | 0.060 |
| F103M | 0.873 | 0.063 | 1.115 | 0.155 |
| F103A | 0.599 | 0.039 | 0.923 | 0.103 |
| F103W | 0.780 | 0.032 | 1.171 | 0.164 |
| F103T | 0.658 | 0.030 | 1.000 | 0.174 |
| F103Y | 0.958 | 0.022 | 0.966 | 0.036 |
| F103G | 0.384 | 0.041 | 0.589 | 0.100 |
| F103S | 0.633 | 0.027 | 0.836 | 0.090 |
| F103N | 0.383 | 0.054 | 0.590 | 0.138 |
| F103H | 0.394 | 0.035 | 0.526 | 0.039 |
| F103P | 0.352 | 0.032 | 0.454 | 0.078 |
| F103Q | 0.364 | 0.038 | 0.478 | 0.127 |
| F103R | 0.357 | 0.030 | 0.589 | 0.043 |
| F103E | 0.371 | 0.030 | 0.464 | 0.026 |
| F103K | 0.364 | 0.033 | 0.577 | 0.136 |
| F103D | 0.345 | 0.030 | 0.506 | 0.081 |
| L104I | 1.173 | 0.064 | 1.196 | 0.142 |
| L104F | 1.207 | 0.015 | 1.041 | 0.054 |
| L104V | 0.879 | 0.039 | 0.927 | 0.085 |
| L104C | 0.734 | 0.030 | 0.889 | 0.114 |
| L104M | 1.168 | 0.066 | 0.982 | 0.173 |
| L104A | 0.954 | 0.075 | 0.987 | 0.143 |
| L104W | 1.010 | 0.057 | 0.886 | 0.134 |
| L104T | 0.592 | 0.040 | 1.008 | 0.142 |
| L104Y | 1.360 | 0.092 | 0.933 | 0.043 |
| L104G | 0.597 | 0.031 | 1.113 | 0.167 |
| L104S | 0.747 | 0.027 | 0.945 | 0.110 |
| L104N | 0.445 | 0.049 | 0.802 | 0.095 |
| L104H | 0.758 | 0.056 | 1.191 | 0.101 |
| L104P | 0.358 | 0.032 | 0.574 | 0.092 |
| L104Q | 0.717 | 0.051 | 1.025 | 0.177 |
| L104R | 0.563 | 0.018 | 0.735 | 0.082 |
| L104E | 0.780 | 0.080 | 1.266 | 0.028 |
| L104K | 0.691 | 0.018 | 0.972 | 0.107 |
| L104D | 0.399 | 0.026 | 0.695 | 0.066 |
| P105I | 0.884 | 0.072 | 1.038 | 0.237 |
| P105L | 0.933 | 0.037 | 1.122 | 0.082 |
| P105F | 0.888 | 0.071 | 1.245 | 0.149 |
| P105V | 0.990 | 0.089 | 0.967 | 0.208 |
| P105C | 1.139 | 0.112 | 1.052 | 0.128 |
| P105A | 1.136 | 0.066 | 1.084 | 0.277 |
| P105W | 0.871 | 0.067 | 0.870 | 0.181 |
| P105T | 1.052 | 0.115 | 0.995 | 0.096 |

| Variant ID | Average WT-SecM Score n = 3 | WT-SecM SD | Average P166A-SecM Score n = 3 | P166A-SecM SD |
|---|---|---|---|---|
| P105Y | 0.926 | 0.063 | 1.200 | 0.123 |
| P105G | 1.138 | 0.058 | 1.027 | 0.199 |
| P105S | 1.097 | 0.059 | 0.946 | 0.087 |
| P105N | 1.209 | 0.095 | 0.737 | 0.284 |
| P105H | 1.113 | 0.112 | 0.967 | 0.118 |
| P105Q | 1.065 | 0.079 | 1.019 | 0.038 |
| P105R | 0.984 | 0.059 | 0.958 | 0.149 |
| P105E | 1.240 | 0.070 | 1.145 | 0.289 |
| P105K | 0.992 | 0.068 | 0.676 | 0.115 |
| P105D | 1.243 | 0.177 | 1.254 | 0.241 |
| K106I | 1.029 | 0.007 | 0.948 | 0.010 |
| K106L | 1.444 | 0.066 | 1.183 | 0.062 |
| K106F | 1.276 | 0.056 | 0.985 | 0.152 |
| K106V | 1.158 | 0.072 | 1.036 | 0.046 |
| K106C | 0.897 | 0.078 | 1.012 | 0.101 |
| K106A | 0.846 | 0.038 | 1.000 | 0.228 |
| K106W | 1.304 | 0.050 | 0.648 | 0.166 |
| K106T | 0.995 | 0.006 | 0.982 | 0.060 |
| K106Y | 1.434 | 0.065 | 1.216 | 0.049 |
| K106G | 0.664 | 0.051 | 1.078 | 0.068 |
| K106S | 0.982 | 0.074 | 1.122 | 0.142 |
| K106N | 1.026 | 0.054 | 0.904 | 0.096 |
| K106H | 1.376 | 0.108 | 1.196 | 0.218 |
| K106P | 0.538 | 0.042 | 1.111 | 0.099 |
| K106Q | 1.055 | 0.024 | 0.958 | 0.119 |
| K106R | 1.228 | 0.082 | 0.998 | 0.096 |
| K106E | 1.019 | 0.064 | 0.924 | 0.034 |
| K106D | 0.853 | 0.052 | 0.816 | 0.217 |
| A107I | 0.659 | 0.081 | 0.915 | 0.163 |
| A107L | 0.446 | 0.028 | 0.719 | 0.058 |
| A107F | 0.385 | 0.035 | 0.761 | 0.013 |
| A107V | 0.868 | 0.027 | 0.947 | 0.126 |
| A107C | 1.179 | 0.038 | 1.231 | 0.193 |
| A107M | 0.434 | 0.036 | 0.734 | 0.032 |
| A107W | 0.377 | 0.042 | 0.496 | 0.224 |
| A107T | 0.855 | 0.042 | 1.044 | 0.038 |
| A107Y | 0.372 | 0.028 | 0.697 | 0.113 |
| A107G | 0.598 | 0.048 | 1.075 | 0.042 |
| A107S | 1.033 | 0.066 | 1.109 | 0.133 |
| A107N | 0.359 | 0.039 | 0.519 | 0.170 |
| A107H | 0.345 | 0.055 | 0.411 | 0.043 |
| A107P | 0.374 | 0.030 | 0.562 | 0.078 |
| A107Q | 0.358 | 0.043 | 0.616 | 0.157 |

| Variant ID | Average WT-SecM Score n = 3 | WT-SecM SD | Average P166A-SecM Score n = 3 | P166A-SecM SD |
|---|---|---|---|---|
| A107R | 0.359 | 0.033 | 0.539 | 0.093 |
| A107E | 0.530 | 0.030 | 0.708 | 0.075 |
| A107K | 0.362 | 0.029 | 0.568 | 0.113 |
| A107D | 0.352 | 0.063 | 0.698 | 0.205 |
| Q108I | 0.773 | 0.055 | 1.001 | 0.037 |
| Q108L | 0.844 | 0.041 | 1.058 | 0.129 |
| Q108F | 0.859 | 0.033 | 0.932 | 0.159 |
| Q108V | 0.960 | 0.052 | 0.993 | 0.143 |
| Q108C | 1.113 | 0.071 | 1.036 | 0.280 |
| Q108A | 1.185 | 0.053 | 1.157 | 0.036 |
| Q108W | 0.782 | 0.062 | 1.405 | 0.089 |
| Q108T | 1.374 | 0.087 | 0.917 | 0.091 |
| Q108Y | 0.885 | 0.055 | 0.973 | 0.089 |
| Q108G | 1.263 | 0.101 | 1.243 | 0.099 |
| Q108S | 1.338 | 0.041 | 1.133 | 0.206 |
| Q108N | 1.293 | 0.086 | 1.455 | 0.048 |
| Q108H | 1.041 | 0.052 | 0.926 | 0.083 |
| Q108P | 0.567 | 0.033 | 0.723 | 0.123 |
| Q108R | 1.149 | 0.065 | 0.976 | 0.081 |
| Q108E | 1.264 | 0.063 | 0.820 | 0.080 |
| Q108K | 1.129 | 0.018 | 0.906 | 0.171 |
| Q108D | 1.342 | 0.081 | 0.638 | 0.002 |
| K109I | 0.896 | 0.033 | 0.988 | 0.100 |
| K109L | 0.971 | 0.024 | 1.048 | 0.151 |
| K109F | 0.908 | 0.089 | 1.008 | 0.056 |
| K109V | 0.938 | 0.086 | 1.056 | 0.329 |
| K109C | 0.956 | 0.063 | 0.935 | 0.071 |
| K109A | 0.750 | 0.046 | 0.993 | 0.211 |
| K109W | 0.707 | 0.031 | 1.240 | 0.482 |
| K109T | 0.920 | 0.022 | 0.874 | 0.100 |
| K109Y | 0.639 | 0.038 | 0.986 | 0.157 |
| K109G | 0.499 | 0.057 | 1.018 | 0.193 |
| K109S | 0.711 | 0.030 | 0.998 | 0.107 |
| K109N | 0.867 | 0.052 | 0.894 | 0.100 |
| K109H | 0.647 | 0.045 | 1.008 | 0.180 |
| K109P | 0.360 | 0.037 | 0.604 | 0.046 |
| K109Q | 0.976 | 0.049 | 0.964 | 0.078 |
| K109R | 1.290 | 0.062 | 1.077 | 0.163 |
| K109E | 0.568 | 0.083 | 0.786 | 0.209 |
| K109D | 0.393 | 0.050 | 0.578 | 0.090 |
| L110I | 1.352 | 0.103 | 0.910 | 0.085 |
| L110F | 0.809 | 0.019 | 0.876 | 0.149 |
| L110V | 0.962 | 0.055 | 1.022 | 0.017 |

| Variant ID | Average WT-SecM Score n = 3 | WT-SecM SD | Average P166A-SecM Score n = 3 | P166A-SecM SD |
|---|---|---|---|---|
| L110C | 0.761 | 0.017 | 0.945 | 0.144 |
| L110M | 1.094 | 0.067 | 0.940 | 0.244 |
| L110A | 0.560 | 0.036 | 0.994 | 0.115 |
| L110W | 0.490 | 0.046 | 0.805 | 0.067 |
| L110T | 0.440 | 0.028 | 0.756 | 0.119 |
| L110Y | 0.418 | 0.027 | 0.600 | 0.048 |
| L110G | 0.371 | 0.020 | 0.464 | 0.088 |
| L110S | 0.402 | 0.039 | 0.594 | 0.078 |
| L110N | 0.375 | 0.032 | 0.652 | 0.135 |
| L110H | 0.381 | 0.032 | 0.564 | 0.052 |
| L110P | 0.487 | 0.040 | 0.686 | 0.091 |
| L110Q | 0.639 | 0.044 | 1.040 | 0.156 |
| L110R | 0.431 | 0.016 | 0.579 | 0.043 |
| L110E | 0.399 | 0.023 | 0.536 | 0.055 |
| L110K | 0.373 | 0.031 | 0.548 | 0.128 |
| L110D | 0.351 | 0.032 | 0.521 | 0.100 |
| Y111I | 1.038 | 0.021 | 1.237 | 0.037 |
| Y111L | 0.747 | 0.070 | 1.035 | 0.243 |
| Y111F | 0.906 | 0.049 | 0.960 | 0.065 |
| Y111V | 0.846 | 0.074 | 1.114 | 0.089 |
| Y111C | 0.653 | 0.023 | 0.858 | 0.075 |
| Y111M | 0.637 | 0.034 | 0.809 | 0.255 |
| Y111A | 0.401 | 0.033 | 0.535 | 0.069 |
| Y111W | 0.467 | 0.027 | 0.766 | 0.156 |
| Y111T | 0.405 | 0.042 | 0.726 | 0.043 |
| Y111G | 0.345 | 0.043 | 0.591 | 0.068 |
| Y111S | 0.384 | 0.045 | 0.510 | 0.066 |
| Y111N | 0.433 | 0.016 | 0.584 | 0.063 |
| Y111H | 0.690 | 0.026 | 0.812 | 0.094 |
| Y111P | 0.346 | 0.031 | 0.566 | 0.081 |
| Y111Q | 0.372 | 0.040 | 0.510 | 0.155 |
| Y111R | 0.363 | 0.043 | 0.554 | 0.063 |
| Y111E | 0.362 | 0.030 | 0.561 | 0.028 |
| Y111K | 0.370 | 0.050 | 0.432 | 0.104 |
| Y111D | 0.883 | 0.067 | 0.824 | 0.100 |
| L112F | 1.197 | 0.074 | 0.963 | 0.239 |
| L112V | 1.043 | 0.065 | 1.027 | 0.150 |
| L112C | 0.673 | 0.068 | 0.942 | 0.207 |
| L112M | 0.965 | 0.032 | 0.819 | 0.075 |
| L112A | 0.489 | 0.033 | 0.776 | 0.062 |
| L112W | 0.552 | 0.050 | 0.843 | 0.079 |
| L112T | 0.518 | 0.049 | 0.809 | 0.146 |
| L112Y | 0.542 | 0.052 | 0.871 | 0.064 |

| Variant ID | Average WT-SecM Score n = 3 | WT-SecM SD | Average P166A-SecM Score n = 3 | P166A-SecM SD |
|---|---|---|---|---|
| L112G | 0.372 | 0.043 | 0.550 | 0.082 |
| L112S | 0.385 | 0.027 | 0.671 | 0.039 |
| L112N | 0.401 | 0.040 | 0.580 | 0.167 |
| L112H | 0.486 | 0.044 | 0.773 | 0.029 |
| L112P | 0.498 | 0.062 | 0.648 | 0.123 |
| L112Q | 0.519 | 0.031 | 0.772 | 0.152 |
| L112R | 0.495 | 0.013 | 0.638 | 0.028 |
| L112E | 0.445 | 0.061 | 0.527 | 0.083 |
| L112K | 0.458 | 0.049 | 0.557 | 0.191 |
| L112D | 0.372 | 0.068 | 0.626 | 0.259 |
| T113I | 1.391 | 0.101 | 1.003 | 0.184 |
| T113L | 0.893 | 0.080 | 0.595 | 0.120 |
| T113F | 0.348 | 0.045 | 0.226 | 0.043 |
| T113V | 1.313 | 0.088 | 0.869 | 0.132 |
| T113C | 1.157 | 0.073 | 0.718 | 0.056 |
| T113M | 1.162 | 0.006 | 0.914 | 0.099 |
| T113A | 1.058 | 0.072 | 0.892 | 0.044 |
| T113Y | 0.718 | 0.039 | 0.468 | 0.084 |
| T113G | 0.532 | 0.023 | 0.887 | 0.112 |
| T113S | 0.994 | 0.049 | 0.863 | 0.178 |
| T113N | 0.817 | 0.013 | 1.056 | 0.080 |
| T113H | 0.927 | 0.108 | 1.034 | 0.052 |
| T113P | 0.482 | 0.006 | 0.665 | 0.093 |
| T113Q | 1.093 | 0.051 | 0.552 | 0.115 |
| T113R | 0.904 | 0.042 | 0.756 | 0.107 |
| T113E | 0.543 | 0.051 | 0.846 | 0.117 |
| T113K | 1.051 | 0.092 | 1.092 | 0.037 |
| T113D | 0.444 | 0.032 | 0.942 | 0.106 |
| H114I | 1.170 | 0.125 | 1.100 | 0.069 |
| H114L | 1.177 | 0.070 | 1.159 | 0.095 |
| H114F | 1.258 | 0.119 | 0.995 | 0.240 |
| H114V | 1.116 | 0.055 | 0.894 | 0.243 |
| H114C | 1.070 | 0.037 | 1.242 | 0.405 |
| H114A | 0.878 | 0.032 | 1.062 | 0.165 |
| H114T | 1.150 | 0.030 | 1.071 | 0.222 |
| H114Y | 1.126 | 0.085 | 1.010 | 0.103 |
| H114G | 0.657 | 0.008 | 0.989 | 0.059 |
| H114S | 1.023 | 0.087 | 1.041 | 0.180 |
| H114N | 1.042 | 0.056 | 0.870 | 0.072 |
| H114P | 0.529 | 0.025 | 0.760 | 0.076 |
| H114Q | 1.062 | 0.131 | 1.077 | 0.160 |
| H114R | 1.214 | 0.079 | 0.984 | 0.157 |
| H114E | 0.817 | 0.014 | 0.663 | 0.004 |

| Variant ID | Average WT-SecM Score n = 3 | WT-SecM SD | Average P166A-SecM Score n = 3 | P166A-SecM SD |
|---|---|---|---|---|
| H114K | 1.413 | 0.099 | 1.211 | 0.124 |
| H114D | 0.699 | 0.018 | 0.833 | 0.124 |
| I115L | 1.058 | 0.076 | 0.940 | 0.067 |
| I115F | 0.944 | 0.063 | 0.986 | 0.183 |
| I115V | 1.232 | 0.110 | 0.922 | 0.041 |
| I115C | 1.045 | 0.036 | 1.076 | 0.132 |
| I115A | 0.764 | 0.039 | 0.981 | 0.065 |
| I115T | 0.950 | 0.047 | 0.972 | 0.106 |
| I115Y | 1.105 | 0.126 | 0.929 | 0.311 |
| I115G | 0.671 | 0.037 | 1.008 | 0.134 |
| I115S | 0.843 | 0.048 | 0.851 | 0.109 |
| I115N | 0.741 | 0.003 | 0.747 | 0.144 |
| I115H | 1.124 | 0.101 | 0.884 | 0.119 |
| I115P | 0.561 | 0.039 | 0.853 | 0.073 |
| I115Q | 0.636 | 0.053 | 0.819 | 0.142 |
| I115R | 0.752 | 0.029 | 0.747 | 0.108 |
| I115E | 0.500 | 0.056 | 0.745 | 0.026 |
| I115K | 0.702 | 0.034 | 1.002 | 0.156 |
| I115D | 0.522 | 0.024 | 0.755 | 0.222 |
| D116I | 1.104 | 0.083 | 0.959 | 0.102 |
| D116L | 1.159 | 0.076 | 1.283 | 0.223 |
| D116F | 1.216 | 0.098 | 1.088 | 0.163 |
| D116V | 1.154 | 0.078 | 1.066 | 0.173 |
| D116C | 1.288 | 0.082 | 1.009 | 0.124 |
| D116M | 1.218 | 0.118 | 1.366 | 0.131 |
| D116A | 1.182 | 0.034 | 1.085 | 0.090 |
| D116T | 1.161 | 0.075 | 0.956 | 0.023 |
| D116Y | 1.257 | 0.047 | 1.127 | 0.139 |
| D116G | 1.120 | 0.086 | 1.029 | 0.090 |
| D116S | 1.182 | 0.085 | 1.063 | 0.034 |
| D116N | 1.188 | 0.044 | 1.011 | 0.076 |
| D116H | 1.281 | 0.117 | 0.987 | 0.011 |
| D116P | 1.068 | 0.077 | 1.088 | 0.119 |
| D116Q | 1.308 | 0.095 | 1.221 | 0.167 |
| D116R | 1.191 | 0.048 | 1.138 | 0.138 |
| D116E | 1.219 | 0.091 | 1.058 | 0.145 |
| D116K | 1.294 | 0.100 | 1.052 | 0.093 |
| A117I | 1.271 | 0.056 | 0.791 | 0.117 |
| A117L | 1.237 | 0.055 | 0.982 | 0.107 |
| A117F | 0.875 | 0.035 | 0.740 | 0.087 |
| A117V | 1.180 | 0.037 | 0.946 | 0.116 |
| A117C | 1.394 | 0.125 | 1.067 | 0.065 |
| A117M | 1.274 | 0.095 | 0.913 | 0.359 |

| Variant ID | Average WT-SecM Score n = 3 | WT-SecM SD | Average P166A-SecM Score n = 3 | P166A-SecM SD |
|---|---|---|---|---|
| A117T | 1.090 | 0.035 | 1.091 | 0.021 |
| A117Y | 1.183 | 0.109 | 0.888 | 0.258 |
| A117G | 1.195 | 0.059 | 1.024 | 0.187 |
| A117S | 1.201 | 0.070 | 0.900 | 0.028 |
| A117N | 1.072 | 0.074 | 1.068 | 0.160 |
| A117H | 1.229 | 0.147 | 0.846 | 0.113 |
| A117P | 0.857 | 0.054 | 0.931 | 0.096 |
| A117Q | 1.222 | 0.072 | 1.145 | 0.171 |
| A117R | 1.175 | 0.088 | 1.063 | 0.004 |
| A117E | 1.024 | 0.062 | 0.856 | 0.062 |
| A117K | 1.214 | 0.122 | 1.136 | 0.028 |
| A117D | 1.116 | 0.057 | 0.997 | 0.139 |
| E118I | 1.226 | 0.042 | 0.893 | 0.112 |
| E118L | 1.234 | 0.073 | 0.781 | 0.114 |
| E118F | 1.103 | 0.041 | 0.887 | 0.140 |
| E118V | 1.269 | 0.063 | 0.830 | 0.140 |
| E118C | 1.267 | 0.027 | 0.918 | 0.136 |
| E118M | 1.191 | 0.101 | 1.007 | 0.038 |
| E118A | 1.229 | 0.080 | 0.898 | 0.029 |
| E118T | 1.292 | 0.085 | 1.060 | 0.200 |
| E118Y | 1.218 | 0.121 | 0.928 | 0.088 |
| E118G | 1.100 | 0.061 | 0.865 | 0.058 |
| E118S | 1.310 | 0.095 | 1.230 | 0.113 |
| E118N | 1.309 | 0.093 | 0.879 | 0.060 |
| E118H | 1.305 | 0.074 | 0.961 | 0.053 |
| E118P | 1.239 | 0.089 | 1.122 | 0.117 |
| E118Q | 1.183 | 0.060 | 1.129 | 0.205 |
| E118R | 1.308 | 0.075 | 0.915 | 0.216 |
| E118K | 1.180 | 0.065 | 1.055 | 0.041 |
| E118D | 1.229 | 0.040 | 0.933 | 0.054 |
| V119I | 1.425 | 0.105 | 1.098 | 0.173 |
| V119L | 1.317 | 0.082 | 1.103 | 0.048 |
| V119F | 1.271 | 0.189 | 1.216 | 0.423 |
| V119C | 1.351 | 0.086 | 0.939 | 0.183 |
| V119M | 1.141 | 0.037 | 1.153 | 0.123 |
| V119A | 1.147 | 0.095 | 1.058 | 0.033 |
| V119T | 1.273 | 0.058 | 0.938 | 0.186 |
| V119Y | 1.289 | 0.101 | 0.752 | 0.206 |
| V119G | 1.217 | 0.077 | 0.972 | 0.146 |
| V119S | 1.224 | 0.069 | 1.008 | 0.131 |
| V119N | 1.090 | 0.074 | 1.309 | 0.191 |
| V119H | 1.215 | 0.051 | 0.811 | 0.166 |
| V119P | 1.137 | 0.108 | 1.149 | 0.061 |

| Variant ID | Average WT-SecM Score n = 3 | WT-SecM SD | Average P166A-SecM Score n = 3 | P166A-SecM SD |
|---|---|---|---|---|
| V119Q | 1.202 | 0.088 | 0.684 | 0.028 |
| V119R | 1.113 | 0.066 | 1.056 | 0.171 |
| V119E | 1.088 | 0.102 | 1.234 | 0.062 |
| V119K | 1.044 | 0.006 | 0.856 | 0.274 |
| V119D | 1.003 | 0.076 | 0.886 | 0.061 |
| E120I | 1.260 | 0.085 | 0.920 | 0.046 |
| E120L | 1.276 | 0.060 | 0.907 | 0.092 |
| E120F | 1.202 | 0.102 | 0.920 | 0.415 |
| E120V | 1.254 | 0.092 | 0.943 | 0.120 |
| E120C | 1.339 | 0.070 | 1.037 | 0.276 |
| E120A | 1.262 | 0.015 | 1.027 | 0.025 |
| E120T | 1.266 | 0.079 | 0.929 | 0.184 |
| E120Y | 1.201 | 0.101 | 1.042 | 0.230 |
| E120G | 1.109 | 0.067 | 0.955 | 0.150 |
| E120S | 1.313 | 0.101 | 1.250 | 0.138 |
| E120N | 1.336 | 0.086 | 1.275 | 0.195 |
| E120H | 1.365 | 0.134 | 0.986 | 0.280 |
| E120P | 1.396 | 0.082 | 1.060 | 0.111 |
| E120Q | 1.165 | 0.077 | 0.964 | 0.151 |
| E120R | 1.281 | 0.059 | 0.841 | 0.163 |
| E120K | 1.160 | 0.061 | 0.932 | 0.132 |
| E120D | 1.137 | 0.067 | 0.999 | 0.062 |
| G121I | 1.159 | 0.101 | 1.262 | 0.308 |
| G121L | 1.184 | 0.118 | 0.837 | 0.122 |
| G121V | 1.063 | 0.071 | 0.923 | 0.114 |
| G121C | 1.077 | 0.046 | 0.960 | 0.103 |
| G121A | 1.240 | 0.079 | 0.763 | 0.144 |
| G121T | 1.203 | 0.111 | 0.826 | 0.121 |
| G121S | 1.199 | 0.110 | 1.063 | 0.130 |
| G121H | 1.180 | 0.133 | 1.002 | 0.215 |
| G121P | 1.112 | 0.079 | 0.934 | 0.030 |
| G121Q | 1.192 | 0.045 | 0.591 | 0.296 |
| G121R | 1.078 | 0.049 | 0.880 | 0.072 |
| G121E | 1.109 | 0.082 | 1.010 | 0.119 |
| G121K | 1.094 | 0.048 | 0.914 | 0.068 |
| G121D | 1.074 | 0.054 | 0.960 | 0.044 |
| D122I | 1.208 | 0.111 | 0.836 | 0.109 |
| D122L | 1.192 | 0.042 | 0.692 | 0.227 |
| D122V | 1.134 | 0.048 | 0.947 | 0.169 |
| D122A | 1.143 | 0.028 | 0.815 | 0.116 |
| D122T | 1.287 | 0.136 | 1.119 | 0.156 |
| D122Y | 1.057 | 0.068 | 0.948 | 0.081 |
| D122G | 1.106 | 0.053 | 0.957 | 0.066 |

| Variant ID | Average WT-SecM Score n = 3 | WT-SecM SD | Average P166A-SecM Score n = 3 | P166A-SecM SD |
|---|---|---|---|---|
| D122S | 1.293 | 0.068 | 0.967 | 0.143 |
| D122N | 1.092 | 0.073 | 1.018 | 0.044 |
| D122H | 1.229 | 0.101 | 0.754 | 0.114 |
| D122P | 1.159 | 0.072 | 0.879 | 0.011 |
| D122Q | 1.351 | 0.058 | 0.607 | 0.166 |
| D122R | 1.216 | 0.076 | 0.835 | 0.155 |
| D122E | 1.220 | 0.090 | 0.699 | 0.080 |
| T123I | 1.134 | 0.042 | 1.096 | 0.048 |
| T123L | 1.304 | 0.041 | 1.192 | 0.314 |
| T123V | 1.392 | 0.030 | 1.461 | 0.309 |
| T123A | 1.092 | 0.085 | 1.023 | 0.116 |
| T123G | 1.210 | 0.138 | 0.889 | 0.013 |
| T123S | 1.307 | 0.065 | 1.087 | 0.103 |
| T123N | 1.179 | 0.079 | 0.757 | 0.221 |
| T123P | 1.028 | 0.020 | 0.925 | 0.105 |
| T123Q | 1.348 | 0.063 | 0.541 | 0.111 |
| T123R | 1.216 | 0.068 | 1.147 | 0.045 |
| H124L | 1.296 | 0.138 | 1.140 | 0.205 |
| H124V | 1.410 | 0.097 | 1.113 | 0.134 |
| H124T | 1.379 | 0.068 | 1.014 | 0.454 |
| H124Y | 1.100 | 0.092 | 0.976 | 0.016 |
| H124S | 1.277 | 0.087 | 1.042 | 0.345 |
| H124N | 1.074 | 0.034 | 0.896 | 0.063 |
| H124P | 1.067 | 0.042 | 1.161 | 0.117 |
| H124Q | 1.098 | 0.027 | 0.884 | 0.179 |
| H124R | 1.206 | 0.051 | 1.128 | 0.135 |
| H124D | 1.073 | 0.026 | 0.849 | 0.074 |
| F125I | 0.771 | 0.069 | 0.936 | 0.059 |
| F125L | 0.953 | 0.047 | 0.774 | 0.062 |
| F125V | 0.676 | 0.049 | 0.755 | 0.150 |
| F125C | 0.821 | 0.057 | 0.830 | 0.077 |
| F125A | 0.671 | 0.051 | 0.690 | 0.084 |
| F125T | 0.557 | 0.023 | 1.000 | 0.035 |
| F125G | 0.647 | 0.022 | 0.852 | 0.082 |
| F125S | 0.769 | 0.058 | 0.790 | 0.140 |
| F125N | 0.664 | 0.044 | 0.642 | 0.263 |
| F125H | 0.671 | 0.089 | 0.467 | 0.276 |
| F125P | 0.539 | 0.030 | 0.596 | 0.109 |
| F125R | 0.550 | 0.046 | 0.758 | 0.119 |
| F125E | 0.510 | 0.078 | 0.468 | 0.036 |
| F125D | 0.545 | 0.011 | 0.595 | 0.204 |
| P126L | 1.069 | 0.052 | 0.914 | 0.057 |
| P126V | 1.148 | 0.106 | 0.924 | 0.048 |

| Variant ID | Average WT-SecM Score n = 3 | WT-SecM SD | Average P166A-SecM Score n = 3 | P166A-SecM SD |
|---|---|---|---|---|
| P126A | 1.009 | 0.035 | 1.021 | 0.075 |
| P126T | 0.983 | 0.046 | 0.930 | 0.125 |
| P126G | 0.814 | 0.072 | 0.996 | 0.304 |
| P126S | 0.878 | 0.021 | 1.043 | 0.024 |
| P126H | 0.761 | 0.087 | 0.945 | 0.260 |
| P126Q | 0.904 | 0.029 | 0.926 | 0.049 |
| P126R | 0.768 | 0.048 | 1.049 | 0.138 |
| P126K | 0.804 | 0.083 | 0.547 | 0.164 |
| P126D | 0.684 | 0.070 | 0.806 | 0.094 |
| D127I | 0.786 | 0.045 | 1.041 | 0.030 |
| D127L | 0.847 | 0.049 | 1.100 | 0.087 |
| D127F | 0.972 | 0.082 | 0.940 | 0.084 |
| D127V | 0.903 | 0.043 | 1.187 | 0.094 |
| D127C | 1.048 | 0.029 | 1.011 | 0.122 |
| D127M | 0.953 | 0.065 | 1.407 | 0.143 |
| D127A | 1.176 | 0.081 | 1.159 | 0.154 |
| D127T | 1.104 | 0.069 | 1.266 | 0.081 |
| D127Y | 1.040 | 0.070 | 0.956 | 0.137 |
| D127G | 1.161 | 0.058 | 1.096 | 0.023 |
| D127S | 1.140 | 0.070 | 1.228 | 0.119 |
| D127N | 1.087 | 0.027 | 0.987 | 0.085 |
| D127H | 1.008 | 0.066 | 0.906 | 0.115 |
| D127P | 1.256 | 0.075 | 1.196 | 0.132 |
| D127Q | 1.073 | 0.007 | 1.194 | 0.181 |
| D127R | 0.874 | 0.052 | 1.102 | 0.089 |
| D127E | 1.231 | 0.079 | 1.135 | 0.119 |
| D127K | 1.051 | 0.086 | 1.179 | 0.284 |
| Y128I | 1.428 | 0.060 | 1.131 | 0.217 |
| Y128L | 1.375 | 0.097 | 1.027 | 0.231 |
| Y128F | 1.425 | 0.096 | 1.115 | 0.272 |
| Y128V | 1.305 | 0.085 | 0.943 | 0.077 |
| Y128C | 1.137 | 0.105 | 0.825 | 0.080 |
| Y128M | 1.228 | 0.095 | 1.419 | 0.215 |
| Y128A | 1.002 | 0.049 | 0.947 | 0.085 |
| Y128W | 1.349 | 0.060 | 1.238 | 0.077 |
| Y128T | 1.245 | 0.088 | 0.969 | 0.169 |
| Y128G | 0.736 | 0.056 | 0.830 | 0.111 |
| Y128S | 0.899 | 0.050 | 0.891 | 0.126 |
| Y128N | 0.817 | 0.014 | 0.868 | 0.105 |
| Y128H | 1.041 | 0.080 | 1.009 | 0.047 |
| Y128P | 0.982 | 0.047 | 0.984 | 0.088 |
| Y128Q | 0.908 | 0.009 | 1.091 | 0.116 |
| Y128R | 0.857 | 0.030 | 1.170 | 0.013 |

| Variant ID | Average WT-SecM Score n = 3 | WT-SecM SD | Average P166A-SecM Score n = 3 | P166A-SecM SD |
|---|---|---|---|---|
| Y128E | 0.718 | 0.060 | 0.876 | 0.204 |
| Y128K | 0.758 | 0.006 | 0.874 | 0.121 |
| Y128D | 0.784 | 0.022 | 1.028 | 0.209 |
| E129I | 1.316 | 0.079 | 0.891 | 0.008 |
| E129L | 1.287 | 0.088 | 1.029 | 0.197 |
| E129F | 1.245 | 0.083 | 0.980 | 0.150 |
| E129V | 1.302 | 0.082 | 1.204 | 0.106 |
| E129C | 1.350 | 0.065 | 1.104 | 0.169 |
| E129M | 1.256 | 0.112 | 1.079 | 0.154 |
| E129A | 1.298 | 0.064 | 1.011 | 0.091 |
| E129W | 1.201 | 0.115 | 1.047 | 0.045 |
| E129T | 1.312 | 0.069 | 1.091 | 0.200 |
| E129Y | 1.265 | 0.016 | 1.029 | 0.185 |
| E129G | 1.135 | 0.060 | 1.064 | 0.053 |
| E129S | 1.347 | 0.110 | 1.231 | 0.071 |
| E129N | 1.358 | 0.097 | 1.158 | 0.142 |
| E129H | 1.341 | 0.044 | 0.910 | 0.152 |
| E129P | 1.283 | 0.089 | 1.177 | 0.119 |
| E129Q | 1.311 | 0.061 | 0.841 | 0.087 |
| E129R | 1.210 | 0.066 | 1.184 | 0.161 |
| E129K | 1.153 | 0.046 | 0.964 | 0.091 |
| E129D | 1.227 | 0.041 | 1.161 | 0.082 |
| P130I | 1.386 | 0.101 | 1.321 | 0.284 |
| P130L | 1.269 | 0.030 | 1.128 | 0.057 |
| P130F | 1.202 | 0.088 | 0.633 | 0.126 |
| P130V | 1.314 | 0.086 | 1.007 | 0.228 |
| P130C | 1.361 | 0.116 | 0.960 | 0.081 |
| P130A | 1.359 | 0.070 | 1.105 | 0.093 |
| P130T | 1.335 | 0.088 | 1.210 | 0.046 |
| P130Y | 1.292 | 0.076 | 0.813 | 0.245 |
| P130G | 1.420 | 0.048 | 1.085 | 0.087 |
| P130S | 1.291 | 0.064 | 0.988 | 0.037 |
| P130N | 1.396 | 0.101 | 1.122 | 0.130 |
| P130H | 1.248 | 0.114 | 0.832 | 0.012 |
| P130Q | 1.187 | 0.031 | 1.006 | 0.022 |
| P130R | 1.290 | 0.115 | 1.115 | 0.085 |
| P130E | 1.343 | 0.024 | 1.161 | 0.210 |
| P130K | 1.378 | 0.077 | 1.377 | 0.331 |
| P130D | 1.406 | 0.055 | 1.171 | 0.146 |
| D131I | 1.049 | 0.074 | 1.063 | 0.140 |
| D131L | 1.211 | 0.101 | 1.100 | 0.129 |
| D131F | 1.071 | 0.104 | 0.960 | 0.177 |
| D131V | 1.068 | 0.075 | 1.144 | 0.177 |

| Variant ID | Average WT-SecM Score n = 3 | WT-SecM SD | Average P166A-SecM Score n = 3 | P166A-SecM SD |
|---|---|---|---|---|
| D131C | 1.342 | 0.067 | 1.063 | 0.107 |
| D131M | 1.322 | 0.059 | 0.802 | 0.302 |
| D131A | 1.290 | 0.056 | 1.162 | 0.219 |
| D131T | 1.311 | 0.066 | 1.068 | 0.134 |
| D131Y | 1.051 | 0.024 | 0.931 | 0.040 |
| D131G | 1.218 | 0.065 | 0.967 | 0.141 |
| D131S | 1.308 | 0.117 | 1.011 | 0.045 |
| D131N | 1.121 | 0.030 | 0.998 | 0.073 |
| D131H | 1.124 | 0.020 | 1.126 | 0.108 |
| D131P | 0.901 | 0.100 | 0.886 | 0.091 |
| D131Q | 1.257 | 0.098 | 1.073 | 0.065 |
| D131R | 1.163 | 0.064 | 1.152 | 0.107 |
| D131E | 1.264 | 0.057 | 0.863 | 0.098 |
| D131K | 1.207 | 0.136 | 1.108 | 0.249 |
| D132I | 1.192 | 0.060 | 1.231 | 0.069 |
| D132L | 1.277 | 0.092 | 1.168 | 0.152 |
| D132F | 1.002 | 0.107 | 1.151 | 0.161 |
| D132V | 1.230 | 0.081 | 1.051 | 0.201 |
| D132C | 1.248 | 0.110 | 1.065 | 0.032 |
| D132M | 1.259 | 0.129 | 0.945 | 0.324 |
| D132A | 1.242 | 0.069 | 0.980 | 0.088 |
| D132W | 0.923 | 0.091 | 0.869 | 0.124 |
| D132T | 1.252 | 0.085 | 1.022 | 0.115 |
| D132Y | 1.012 | 0.006 | 0.920 | 0.058 |
| D132G | 1.130 | 0.071 | 1.010 | 0.100 |
| D132S | 1.243 | 0.073 | 0.966 | 0.093 |
| D132N | 1.199 | 0.081 | 1.105 | 0.112 |
| D132H | 1.284 | 0.075 | 1.038 | 0.184 |
| D132P | 0.888 | 0.084 | 0.909 | 0.028 |
| D132Q | 1.342 | 0.124 | 0.978 | 0.055 |
| D132R | 1.148 | 0.075 | 1.071 | 0.122 |
| D132E | 1.331 | 0.090 | 1.016 | 0.130 |
| D132K | 1.249 | 0.105 | 1.087 | 0.109 |
| W133I | 0.420 | 0.026 | 0.751 | 0.015 |
| W133L | 0.595 | 0.032 | 0.915 | 0.100 |
| W133F | 1.222 | 0.104 | 1.038 | 0.049 |
| W133V | 0.425 | 0.040 | 0.632 | 0.041 |
| W133C | 0.491 | 0.026 | 0.723 | 0.054 |
| W133M | 0.588 | 0.063 | 0.718 | 0.119 |
| W133A | 0.391 | 0.039 | 0.611 | 0.093 |
| W133T | 0.408 | 0.048 | 0.523 | 0.072 |
| W133Y | 0.948 | 0.034 | 0.873 | 0.144 |
| W133G | 0.425 | 0.015 | 0.633 | 0.073 |

| Variant ID | Average WT-SecM Score n = 3 | WT-SecM SD | Average P166A-SecM Score n = 3 | P166A-SecM SD |
|---|---|---|---|---|
| W133S | 0.389 | 0.043 | 0.651 | 0.094 |
| W133N | 0.376 | 0.034 | 0.666 | 0.102 |
| W133H | 0.447 | 0.027 | 0.681 | 0.025 |
| W133P | 0.381 | 0.047 | 0.682 | 0.133 |
| W133Q | 0.398 | 0.034 | 0.634 | 0.164 |
| W133R | 0.485 | 0.036 | 0.669 | 0.107 |
| W133E | 0.388 | 0.041 | 0.639 | 0.113 |
| W133K | 0.387 | 0.046 | 0.542 | 0.114 |
| W133D | 0.374 | 0.038 | 0.620 | 0.096 |
| E134I | 1.307 | 0.094 | 1.275 | 0.081 |
| E134L | 1.247 | 0.091 | 1.063 | 0.188 |
| E134F | 1.199 | 0.097 | 1.036 | 0.182 |
| E134V | 1.322 | 0.061 | 1.207 | 0.109 |
| E134C | 1.315 | 0.103 | 1.162 | 0.073 |
| E134M | 1.207 | 0.064 | 1.006 | 0.070 |
| E134A | 1.196 | 0.087 | 0.966 | 0.207 |
| E134W | 1.024 | 0.094 | 1.060 | 0.190 |
| E134T | 1.336 | 0.068 | 1.083 | 0.177 |
| E134Y | 1.223 | 0.084 | 1.060 | 0.222 |
| E134G | 1.066 | 0.083 | 1.063 | 0.093 |
| E134S | 1.098 | 0.068 | 1.011 | 0.150 |
| E134N | 1.343 | 0.056 | 1.085 | 0.269 |
| E134H | 1.253 | 0.144 | 1.318 | 0.147 |
| E134P | 0.805 | 0.043 | 1.085 | 0.165 |
| E134Q | 1.270 | 0.086 | 0.851 | 0.187 |
| E134R | 1.142 | 0.073 | 1.149 | 0.150 |
| E134K | 1.146 | 0.076 | 1.038 | 0.177 |
| E134D | 1.259 | 0.120 | 0.905 | 0.072 |
| S135I | 1.446 | 0.091 | 1.068 | 0.212 |
| S135L | 1.356 | 0.123 | 0.942 | 0.128 |
| S135F | 1.011 | 0.041 | 1.286 | 0.168 |
| S135V | 1.439 | 0.090 | 1.048 | 0.293 |
| S135C | 1.375 | 0.064 | 0.776 | 0.175 |
| S135A | 1.306 | 0.086 | 1.088 | 0.218 |
| S135W | 0.937 | 0.082 | 0.964 | 0.341 |
| S135T | 1.362 | 0.108 | 0.870 | 0.221 |
| S135Y | 1.078 | 0.059 | 0.663 | 0.098 |
| S135G | 1.108 | 0.080 | 1.067 | 0.329 |
| S135N | 1.313 | 0.092 | 1.280 | 0.152 |
| S135H | 1.249 | 0.097 | 0.996 | 0.186 |
| S135P | 1.284 | 0.064 | 0.938 | 0.169 |
| S135Q | 1.365 | 0.046 | 0.774 | 0.182 |
| S135R | 1.333 | 0.093 | 1.282 | 0.067 |

| Variant ID | Average WT-SecM Score n = 3 | WT-SecM SD | Average P166A-SecM Score n = 3 | P166A-SecM SD |
|---|---|---|---|---|
| S135E | 1.272 | 0.068 | 0.987 | 0.245 |
| S135K | 1.299 | 0.123 | 1.502 | 0.156 |
| S135D | 0.900 | 0.099 | 1.007 | 0.279 |
| V136I | 1.209 | 0.085 | 0.945 | 0.173 |
| V136L | 1.088 | 0.063 | 0.996 | 0.121 |
| V136F | 0.899 | 0.079 | 0.802 | 0.123 |
| V136C | 1.247 | 0.094 | 1.007 | 0.074 |
| V136A | 1.099 | 0.082 | 1.034 | 0.135 |
| V136W | 0.676 | 0.044 | 1.183 | 0.163 |
| V136T | 1.417 | 0.087 | 1.198 | 0.269 |
| V136Y | 0.809 | 0.079 | 0.989 | 0.113 |
| V136G | 0.881 | 0.044 | 1.154 | 0.050 |
| V136S | 1.180 | 0.076 | 1.102 | 0.102 |
| V136N | 0.978 | 0.110 | 1.075 | 0.332 |
| V136H | 1.017 | 0.069 | 0.863 | 0.159 |
| V136P | 0.647 | 0.045 | 0.905 | 0.047 |
| V136Q | 1.036 | 0.110 | 1.179 | 0.203 |
| V136R | 1.072 | 0.071 | 1.125 | 0.267 |
| V136E | 0.937 | 0.092 | 0.977 | 0.118 |
| V136K | 1.175 | 0.057 | 1.287 | 0.019 |
| V136D | 0.810 | 0.045 | 1.093 | 0.197 |
| F137I | 0.957 | 0.071 | 0.746 | 0.247 |
| F137L | 1.090 | 0.085 | 1.020 | 0.076 |
| F137V | 1.007 | 0.060 | 1.006 | 0.108 |
| F137C | 1.041 | 0.073 | 0.966 | 0.220 |
| F137M | 1.173 | 0.008 | 0.597 | 0.151 |
| F137A | 1.038 | 0.070 | 0.841 | 0.067 |
| F137W | 1.225 | 0.109 | 1.071 | 0.579 |
| F137T | 0.997 | 0.054 | 0.718 | 0.154 |
| F137Y | 1.326 | 0.123 | 0.945 | 0.131 |
| F137G | 0.871 | 0.026 | 0.931 | 0.178 |
| F137S | 1.072 | 0.078 | 0.987 | 0.155 |
| F137N | 0.863 | 0.047 | 1.198 | 0.207 |
| F137H | 1.358 | 0.096 | 1.062 | 0.218 |
| F137P | 0.487 | 0.018 | 0.842 | 0.151 |
| F137Q | 1.040 | 0.027 | 1.046 | 0.212 |
| F137R | 0.903 | 0.017 | 0.982 | 0.173 |
| F137E | 1.075 | 0.109 | 0.779 | 0.057 |
| F137K | 0.920 | 0.066 | 0.922 | 0.128 |
| F137D | 0.861 | 0.090 | 0.964 | 0.068 |
| S138I | 1.373 | 0.096 | 1.065 | 0.061 |
| S138L | 1.455 | 0.075 | 0.977 | 0.149 |
| S138F | 1.373 | 0.112 | 1.070 | 0.267 |

| Variant ID | Average WT-SecM Score n = 3 | WT-SecM SD | Average P166A-SecM Score n = 3 | P166A-SecM SD |
|---|---|---|---|---|
| S138V | 1.431 | 0.094 | 0.931 | 0.192 |
| S138C | 1.434 | 0.074 | 0.998 | 0.229 |
| S138A | 1.326 | 0.090 | 1.233 | 0.154 |
| S138W | 1.375 | 0.107 | 0.677 | 0.047 |
| S138T | 1.387 | 0.110 | 0.913 | 0.118 |
| S138Y | 1.409 | 0.110 | 0.733 | 0.162 |
| S138G | 1.089 | 0.044 | 0.903 | 0.121 |
| S138N | 1.170 | 0.053 | 1.085 | 0.024 |
| S138H | 1.458 | 0.117 | 0.917 | 0.047 |
| S138P | 0.479 | 0.037 | 0.854 | 0.181 |
| S138Q | 1.464 | 0.117 | 0.803 | 0.162 |
| S138R | 1.285 | 0.060 | 1.092 | 0.121 |
| S138E | 1.413 | 0.057 | 0.883 | 0.129 |
| S138K | 1.392 | 0.116 | 1.029 | 0.142 |
| S138D | 1.248 | 0.092 | 1.116 | 0.217 |
| E139I | 1.121 | 0.077 | 1.107 | 0.145 |
| E139L | 0.946 | 0.054 | 0.997 | 0.071 |
| E139F | 0.908 | 0.092 | 1.196 | 0.201 |
| E139V | 1.119 | 0.038 | 0.999 | 0.070 |
| E139C | 1.118 | 0.082 | 0.940 | 0.204 |
| E139M | 1.039 | 0.073 | 1.097 | 0.134 |
| E139A | 1.055 | 0.008 | 0.999 | 0.084 |
| E139W | 0.817 | 0.064 | 0.762 | 0.227 |
| E139T | 1.131 | 0.113 | 0.987 | 0.109 |
| E139Y | 0.918 | 0.056 | 0.954 | 0.189 |
| E139G | 0.895 | 0.056 | 0.989 | 0.087 |
| E139S | 1.249 | 0.088 | 1.001 | 0.161 |
| E139N | 0.907 | 0.061 | 0.920 | 0.107 |
| E139H | 0.891 | 0.078 | 1.033 | 0.179 |
| E139P | 0.447 | 0.046 | 0.648 | 0.130 |
| E139Q | 0.925 | 0.044 | 0.975 | 0.107 |
| E139R | 0.905 | 0.038 | 0.965 | 0.057 |
| E139K | 0.930 | 0.020 | 0.901 | 0.085 |
| E139D | 1.118 | 0.092 | 0.811 | 0.173 |
| F140I | 1.359 | 0.081 | 1.082 | 0.200 |
| F140L | 1.238 | 0.072 | 1.011 | 0.160 |
| F140V | 1.358 | 0.099 | 1.161 | 0.191 |
| F140C | 1.358 | 0.018 | 1.080 | 0.251 |
| F140M | 1.342 | 0.081 | 1.186 | 0.136 |
| F140A | 1.417 | 0.118 | 1.049 | 0.188 |
| F140W | 1.332 | 0.066 | 0.782 | 0.140 |
| F140T | 1.418 | 0.120 | 1.098 | 0.136 |
| F140Y | 1.400 | 0.054 | 1.117 | 0.290 |

| Variant ID | Average WT-SecM Score n = 3 | WT-SecM SD | Average P166A-SecM Score n = 3 | P166A-SecM SD |
|---|---|---|---|---|
| F140G | 1.324 | 0.079 | 1.146 | 0.081 |
| F140S | 1.375 | 0.068 | 1.037 | 0.166 |
| F140N | 1.401 | 0.119 | 1.077 | 0.112 |
| F140H | 1.462 | 0.132 | 0.828 | 0.206 |
| F140P | 1.352 | 0.102 | 1.251 | 0.067 |
| F140Q | 1.440 | 0.064 | 1.171 | 0.114 |
| F140R | 1.398 | 0.085 | 1.357 | 0.195 |
| F140E | 1.444 | 0.086 | 0.729 | 0.090 |
| F140K | 1.383 | 0.124 | 0.758 | 0.275 |
| F140D | 1.402 | 0.113 | 1.321 | 0.204 |
| H141I | 1.017 | 0.083 | 1.179 | 0.263 |
| H141L | 0.892 | 0.049 | 0.976 | 0.218 |
| H141F | 1.207 | 0.153 | 1.172 | 0.190 |
| H141V | 1.066 | 0.080 | 1.011 | 0.187 |
| H141C | 1.139 | 0.114 | 0.792 | 0.124 |
| H141M | 0.916 | 0.048 | 1.056 | 0.061 |
| H141A | 0.954 | 0.071 | 0.955 | 0.053 |
| H141W | 1.072 | 0.016 | 0.928 | 0.138 |
| H141T | 0.837 | 0.049 | 0.875 | 0.144 |
| H141Y | 1.146 | 0.043 | 0.888 | 0.012 |
| H141G | 0.971 | 0.040 | 1.115 | 0.080 |
| H141S | 0.963 | 0.049 | 0.883 | 0.131 |
| H141N | 1.105 | 0.032 | 0.990 | 0.082 |
| H141P | 0.890 | 0.032 | 0.979 | 0.038 |
| H141Q | 0.986 | 0.050 | 1.011 | 0.154 |
| H141R | 1.026 | 0.057 | 0.984 | 0.062 |
| H141E | 0.779 | 0.019 | 0.938 | 0.106 |
| H141K | 0.993 | 0.054 | 1.126 | 0.245 |
| H141D | 0.752 | 0.016 | 0.919 | 0.081 |
| D142I | 1.235 | 0.089 | 0.954 | 0.235 |
| D142L | 1.220 | 0.071 | 0.952 | 0.083 |
| D142F | 1.170 | 0.079 | 0.805 | 0.215 |
| D142V | 1.232 | 0.097 | 1.132 | 0.082 |
| D142C | 1.252 | 0.122 | 0.994 | 0.046 |
| D142M | 1.141 | 0.075 | 0.997 | 0.263 |
| D142A | 1.302 | 0.065 | 1.195 | 0.242 |
| D142W | 0.993 | 0.020 | 0.901 | 0.105 |
| D142T | 1.269 | 0.086 | 1.089 | 0.063 |
| D142Y | 1.091 | 0.050 | 1.056 | 0.014 |
| D142G | 1.186 | 0.056 | 0.933 | 0.147 |
| D142S | 1.312 | 0.093 | 1.086 | 0.068 |
| D142N | 1.173 | 0.052 | 0.847 | 0.042 |
| D142H | 1.287 | 0.065 | 1.144 | 0.178 |

| Variant ID | Average WT-SecM Score n = 3 | WT-SecM SD | Average P166A-SecM Score n = 3 | P166A-SecM SD |
|---|---|---|---|---|
| D142P | 1.393 | 0.054 | 1.171 | 0.264 |
| D142Q | 1.128 | 0.053 | 0.789 | 0.165 |
| D142R | 1.124 | 0.061 | 0.839 | 0.152 |
| D142E | 1.220 | 0.030 | 0.994 | 0.161 |
| D142K | 1.166 | 0.063 | 0.989 | 0.157 |
| A143I | 1.248 | 0.074 | 0.826 | 0.075 |
| A143L | 1.127 | 0.029 | 1.020 | 0.215 |
| A143F | 1.123 | 0.050 | 0.856 | 0.145 |
| A143V | 1.171 | 0.059 | 1.089 | 0.033 |
| A143C | 1.313 | 0.111 | 1.072 | 0.127 |
| A143W | 1.018 | 0.040 | 1.279 | 0.271 |
| A143T | 1.146 | 0.058 | 0.847 | 0.190 |
| A143Y | 1.051 | 0.082 | 1.078 | 0.227 |
| A143G | 1.243 | 0.062 | 1.046 | 0.238 |
| A143S | 1.322 | 0.100 | 1.003 | 0.053 |
| A143N | 1.192 | 0.104 | 0.905 | 0.210 |
| A143H | 1.200 | 0.091 | 1.072 | 0.189 |
| A143P | 1.313 | 0.088 | 1.075 | 0.136 |
| A143Q | 1.303 | 0.121 | 0.980 | 0.177 |
| A143R | 1.273 | 0.107 | 1.063 | 0.108 |
| A143E | 1.244 | 0.071 | 1.072 | 0.136 |
| A143K | 1.329 | 0.096 | 0.784 | 0.129 |
| A143D | 1.120 | 0.056 | 1.016 | 0.164 |
| D144I | 0.792 | 0.050 | 1.114 | 0.280 |
| D144L | 0.788 | 0.078 | 0.920 | 0.125 |
| D144F | 0.760 | 0.051 | 1.040 | 0.186 |
| D144V | 0.832 | 0.034 | 0.997 | 0.266 |
| D144C | 1.108 | 0.078 | 1.207 | 0.176 |
| D144M | 0.767 | 0.055 | 1.277 | 0.127 |
| D144A | 0.885 | 0.077 | 1.013 | 0.102 |
| D144W | 0.754 | 0.033 | 0.670 | 0.142 |
| D144T | 1.027 | 0.077 | 1.173 | 0.173 |
| D144Y | 0.767 | 0.048 | 1.066 | 0.134 |
| D144G | 1.035 | 0.051 | 1.063 | 0.162 |
| D144S | 1.070 | 0.077 | 1.035 | 0.253 |
| D144N | 1.054 | 0.054 | 1.044 | 0.173 |
| D144H | 0.882 | 0.066 | 0.760 | 0.126 |
| D144P | 0.846 | 0.051 | 1.028 | 0.169 |
| D144Q | 0.877 | 0.051 | 1.047 | 0.306 |
| D144R | 0.788 | 0.050 | 0.824 | 0.090 |
| D144E | 1.025 | 0.051 | 0.959 | 0.110 |
| D144K | 0.816 | 0.043 | 1.038 | 0.322 |
| A145I | 1.275 | 0.146 | 1.000 | 0.091 |

| Variant ID | Average WT-SecM Score n = 3 | WT-SecM SD | Average P166A-SecM Score n = 3 | P166A-SecM SD |
|---|---|---|---|---|
| A145L | 1.335 | 0.088 | 1.057 | 0.272 |
| A145F | 1.320 | 0.136 | 0.873 | 0.037 |
| A145V | 1.135 | 0.034 | 0.964 | 0.099 |
| A145C | 1.376 | 0.114 | 1.192 | 0.255 |
| A145M | 1.370 | 0.078 | 0.973 | 0.053 |
| A145W | 1.319 | 0.112 | 0.998 | 0.169 |
| A145T | 1.265 | 0.073 | 0.937 | 0.088 |
| A145Y | 1.292 | 0.132 | 0.867 | 0.119 |
| A145G | 1.324 | 0.064 | 0.816 | 0.245 |
| A145S | 1.350 | 0.092 | 1.094 | 0.085 |
| A145N | 1.390 | 0.126 | 1.109 | 0.172 |
| A145H | 1.285 | 0.091 | 1.074 | 0.077 |
| A145P | 1.303 | 0.095 | 1.258 | 0.084 |
| A145Q | 1.286 | 0.065 | 1.153 | 0.143 |
| A145R | 1.228 | 0.065 | 0.990 | 0.035 |
| A145E | 1.353 | 0.093 | 0.883 | 0.100 |
| A145K | 1.299 | 0.031 | 0.955 | 0.334 |
| A145D | 1.464 | 0.050 | 0.683 | 0.219 |
| Q146I | 1.372 | 0.078 | 0.991 | 0.121 |
| Q146L | 1.349 | 0.071 | 1.081 | 0.124 |
| Q146F | 1.385 | 0.063 | 0.910 | 0.177 |
| Q146V | 1.360 | 0.058 | 1.064 | 0.128 |
| Q146C | 1.358 | 0.078 | 1.198 | 0.183 |
| Q146M | 1.348 | 0.070 | 0.986 | 0.105 |
| Q146A | 1.367 | 0.092 | 1.076 | 0.254 |
| Q146W | 1.360 | 0.070 | 0.918 | 0.201 |
| Q146T | 1.379 | 0.098 | 1.078 | 0.219 |
| Q146Y | 1.363 | 0.125 | 1.133 | 0.060 |
| Q146G | 1.252 | 0.082 | 0.965 | 0.295 |
| Q146S | 1.392 | 0.092 | 1.179 | 0.181 |
| Q146N | 1.398 | 0.108 | 1.068 | 0.110 |
| Q146H | 1.359 | 0.067 | 1.066 | 0.155 |
| Q146P | 1.203 | 0.057 | 1.009 | 0.148 |
| Q146R | 1.260 | 0.064 | 0.991 | 0.146 |
| Q146E | 1.421 | 0.103 | 0.940 | 0.094 |
| Q146K | 1.235 | 0.055 | 1.060 | 0.154 |
| Q146D | 1.464 | 0.094 | 1.061 | 0.228 |
| N147I | 1.074 | 0.089 | 0.849 | 0.151 |
| N147L | 0.964 | 0.068 | 0.964 | 0.192 |
| N147F | 1.172 | 0.116 | 0.999 | 0.210 |
| N147V | 1.008 | 0.080 | 0.989 | 0.163 |
| N147C | 1.217 | 0.095 | 0.933 | 0.175 |
| N147M | 1.040 | 0.089 | 0.968 | 0.420 |

| Variant ID | Average WT-SecM Score n = 3 | WT-SecM SD | Average P166A-SecM Score n = 3 | P166A-SecM SD |
|---|---|---|---|---|
| N147A | 1.080 | 0.088 | 0.908 | 0.175 |
| N147W | 0.985 | 0.097 | 0.998 | 0.147 |
| N147T | 1.064 | 0.046 | 0.919 | 0.165 |
| N147Y | 1.094 | 0.062 | 1.101 | 0.107 |
| N147G | 1.138 | 0.101 | 1.198 | 0.087 |
| N147S | 1.120 | 0.071 | 1.006 | 0.146 |
| N147H | 1.125 | 0.074 | 0.956 | 0.213 |
| N147P | 0.741 | 0.045 | 1.008 | 0.176 |
| N147Q | 0.957 | 0.050 | 0.966 | 0.151 |
| N147R | 0.868 | 0.055 | 1.063 | 0.179 |
| N147E | 1.036 | 0.044 | 0.793 | 0.267 |
| N147K | 0.918 | 0.045 | 0.897 | 0.065 |
| N147D | 1.095 | 0.063 | 0.833 | 0.117 |
| S148I | 1.332 | 0.071 | 0.998 | 0.139 |
| S148L | 1.295 | 0.070 | 1.150 | 0.132 |
| S148F | 1.208 | 0.059 | 0.911 | 0.246 |
| S148V | 1.295 | 0.087 | 1.109 | 0.165 |
| S148C | 1.273 | 0.077 | 1.138 | 0.220 |
| S148M | 1.311 | 0.103 | 0.897 | 0.236 |
| S148A | 1.358 | 0.069 | 1.114 | 0.163 |
| S148W | 1.177 | 0.061 | 1.070 | 0.281 |
| S148T | 1.379 | 0.053 | 1.046 | 0.175 |
| S148Y | 1.180 | 0.066 | 1.115 | 0.121 |
| S148G | 1.295 | 0.103 | 1.073 | 0.097 |
| S148N | 1.292 | 0.062 | 1.244 | 0.273 |
| S148H | 1.283 | 0.073 | 1.075 | 0.263 |
| S148P | 1.351 | 0.098 | 1.103 | 0.151 |
| S148Q | 1.340 | 0.075 | 0.943 | 0.184 |
| S148R | 1.171 | 0.065 | 1.074 | 0.180 |
| S148E | 1.352 | 0.122 | 1.040 | 0.071 |
| S148K | 1.275 | 0.072 | 1.078 | 0.218 |
| S148D | 1.336 | 0.121 | 0.968 | 0.260 |
| H149I | 1.337 | 0.101 | 0.900 | 0.086 |
| H149L | 1.277 | 0.086 | 0.921 | 0.142 |
| H149F | 1.372 | 0.085 | 1.113 | 0.110 |
| H149V | 1.382 | 0.082 | 0.975 | 0.204 |
| H149C | 1.375 | 0.074 | 1.085 | 0.189 |
| H149M | 1.284 | 0.086 | 0.984 | 0.208 |
| H149A | 1.227 | 0.102 | 1.026 | 0.130 |
| H149W | 1.263 | 0.056 | 0.946 | 0.083 |
| H149T | 1.300 | 0.092 | 0.956 | 0.149 |
| H149Y | 1.283 | 0.050 | 1.152 | 0.127 |
| H149G | 0.972 | 0.058 | 1.080 | 0.110 |

| Variant ID | Average WT-SecM Score n = 3 | WT-SecM SD | Average P166A-SecM Score n = 3 | P166A-SecM SD |
|---|---|---|---|---|
| H149S | 1.200 | 0.085 | 0.919 | 0.148 |
| H149N | 1.189 | 0.059 | 1.121 | 0.039 |
| H149P | 1.210 | 0.053 | 0.965 | 0.144 |
| H149Q | 1.127 | 0.021 | 0.994 | 0.087 |
| H149R | 1.067 | 0.073 | 0.991 | 0.116 |
| H149E | 1.191 | 0.078 | 0.809 | 0.111 |
| H149K | 1.056 | 0.118 | 1.300 | 0.291 |
| H149D | 1.051 | 0.086 | 0.970 | 0.128 |
| S150I | 1.025 | 0.071 | 1.007 | 0.177 |
| S150L | 0.964 | 0.062 | 1.043 | 0.127 |
| S150F | 1.035 | 0.063 | 0.927 | 0.227 |
| S150V | 1.133 | 0.071 | 0.935 | 0.142 |
| S150C | 1.302 | 0.119 | 0.983 | 0.117 |
| S150M | 1.142 | 0.045 | 1.030 | 0.165 |
| S150A | 1.379 | 0.078 | 1.098 | 0.069 |
| S150W | 0.900 | 0.063 | 0.939 | 0.185 |
| S150T | 1.264 | 0.059 | 1.064 | 0.191 |
| S150Y | 1.031 | 0.096 | 0.923 | 0.257 |
| S150G | 1.258 | 0.074 | 1.096 | 0.125 |
| S150N | 1.227 | 0.075 | 0.912 | 0.032 |
| S150H | 1.205 | 0.095 | 0.898 | 0.152 |
| S150P | 1.408 | 0.119 | 1.134 | 0.157 |
| S150Q | 1.255 | 0.091 | 1.164 | 0.134 |
| S150R | 1.185 | 0.073 | 1.077 | 0.117 |
| S150E | 1.150 | 0.067 | 0.930 | 0.091 |
| S150K | 1.223 | 0.108 | 1.109 | 0.078 |
| S150D | 1.332 | 0.097 | 0.954 | 0.135 |
| Y151I | 1.281 | 0.087 | 1.173 | 0.121 |
| Y151L | 1.305 | 0.102 | 1.026 | 0.233 |
| Y151C | 1.171 | 0.090 | 1.013 | 0.149 |
| Y151M | 1.372 | 0.091 | 0.863 | 0.160 |
| Y151W | 1.293 | 0.144 | 0.919 | 0.221 |
| Y151T | 1.049 | 0.051 | 1.043 | 0.163 |
| Y151G | 0.635 | 0.040 | 1.000 | 0.145 |
| Y151S | 0.869 | 0.056 | 0.993 | 0.067 |
| Y151N | 0.690 | 0.020 | 0.949 | 0.216 |
| Y151H | 1.141 | 0.088 | 0.940 | 0.125 |
| Y151P | 0.530 | 0.047 | 0.893 | 0.152 |
| Y151Q | 0.936 | 0.078 | 0.998 | 0.142 |
| Y151R | 0.981 | 0.079 | 0.990 | 0.040 |
| Y151K | 0.720 | 0.036 | 0.916 | 0.033 |
| Y151D | 0.508 | 0.033 | 0.715 | 0.152 |
| S152I | 1.331 | 0.119 | 0.915 | 0.212 |

| Variant ID | Average WT-SecM Score n = 3 | WT-SecM SD | Average P166A-SecM Score n = 3 | P166A-SecM SD |
|---|---|---|---|---|
| S152L | 0.985 | 0.126 | 0.625 | 0.069 |
| S152F | 1.283 | 0.125 | 0.908 | 0.342 |
| S152V | 1.390 | 0.166 | 1.015 | 0.275 |
| S152C | 1.241 | 0.122 | 0.779 | 0.061 |
| S152A | 1.330 | 0.115 | 0.825 | 0.019 |
| S152W | 1.226 | 0.038 | 0.931 | 0.031 |
| S152T | 1.416 | 0.098 | 0.966 | 0.159 |
| S152Y | 1.398 | 0.144 | 1.228 | 0.221 |
| S152G | 1.199 | 0.099 | 0.977 | 0.165 |
| S152N | 1.137 | 0.104 | 0.935 | 0.114 |
| S152H | 1.401 | 0.073 | 0.949 | 0.137 |
| S152P | 0.399 | 0.046 | 0.770 | 0.130 |
| S152Q | 1.339 | 0.071 | 1.132 | 0.122 |
| S152R | 1.342 | 0.094 | 0.801 | 0.115 |
| S152E | 1.269 | 0.094 | 0.844 | 0.160 |
| S152K | 1.448 | 0.095 | 0.916 | 0.276 |
| S152D | 0.926 | 0.082 | 0.881 | 0.143 |
| F153I | 1.053 | 0.055 | 0.966 | 0.102 |
| F153L | 1.065 | 0.057 | 0.903 | 0.089 |
| F153V | 1.004 | 0.057 | 0.983 | 0.234 |
| F153C | 0.793 | 0.041 | 0.887 | 0.201 |
| F153M | 0.953 | 0.063 | 0.883 | 0.227 |
| F153A | 0.580 | 0.046 | 0.863 | 0.168 |
| F153W | 1.167 | 0.122 | 1.142 | 0.010 |
| F153T | 0.777 | 0.053 | 0.870 | 0.186 |
| F153Y | 1.042 | 0.062 | 0.878 | 0.112 |
| F153G | 0.409 | 0.042 | 0.739 | 0.139 |
| F153S | 0.616 | 0.038 | 0.985 | 0.152 |
| F153N | 0.733 | 0.046 | 0.817 | 0.244 |
| F153H | 1.157 | 0.107 | 1.203 | 0.186 |
| F153P | 0.435 | 0.025 | 0.836 | 0.126 |
| F153Q | 0.724 | 0.031 | 1.035 | 0.076 |
| F153R | 0.726 | 0.031 | 1.093 | 0.087 |
| F153E | 0.578 | 0.076 | 0.985 | 0.144 |
| F153K | 0.503 | 0.022 | 0.951 | 0.204 |
| F153D | 0.475 | 0.039 | 0.812 | 0.170 |
| E154I | 1.367 | 0.121 | 0.985 | 0.117 |
| E154L | 1.270 | 0.091 | 1.039 | 0.204 |
| E154F | 1.165 | 0.113 | 1.021 | 0.193 |
| E154V | 1.368 | 0.134 | 1.071 | 0.167 |
| E154C | 1.273 | 0.103 | 1.106 | 0.203 |
| E154M | 1.226 | 0.087 | 1.151 | 0.011 |
| E154A | 1.110 | 0.099 | 0.957 | 0.173 |

| Variant ID | Average<br>WT-SecM Score<br>n = 3 | WT-SecM<br>SD | Average<br>P166A-SecM Score<br>n = 3 | P166A-SecM<br>SD |
|---|---|---|---|---|
| E154W | 1.120 | 0.066 | 1.148 | 0.425 |
| E154T | 1.340 | 0.098 | 1.068 | 0.236 |
| E154Y | 1.074 | 0.101 | 0.931 | 0.183 |
| E154G | 0.836 | 0.037 | 1.041 | 0.101 |
| E154S | 1.299 | 0.084 | 1.230 | 0.123 |
| E154N | 1.230 | 0.090 | 0.882 | 0.118 |
| E154H | 1.436 | 0.127 | 1.093 | 0.192 |
| E154P | 0.379 | 0.043 | 0.729 | 0.102 |
| E154Q | 1.254 | 0.046 | 1.012 | 0.092 |
| E154R | 1.103 | 0.071 | 1.105 | 0.220 |
| E154K | 1.068 | 0.060 | 0.915 | 0.138 |
| E154D | 1.015 | 0.078 | 1.182 | 0.192 |
| I155L | 0.866 | 0.069 | 0.930 | 0.151 |
| I155F | 0.540 | 0.018 | 0.872 | 0.085 |
| I155V | 1.183 | 0.085 | 1.062 | 0.105 |
| I155C | 0.886 | 0.045 | 0.822 | 0.259 |
| I155M | 0.813 | 0.073 | 1.001 | 0.097 |
| I155A | 0.624 | 0.049 | 1.015 | 0.135 |
| I155W | 0.481 | 0.018 | 0.690 | 0.092 |
| I155T | 0.803 | 0.068 | 0.869 | 0.193 |
| I155Y | 0.567 | 0.039 | 0.984 | 0.175 |
| I155G | 0.424 | 0.029 | 0.830 | 0.127 |
| I155S | 0.520 | 0.031 | 0.959 | 0.130 |
| I155N | 0.461 | 0.032 | 0.976 | 0.131 |
| I155H | 0.476 | 0.037 | 1.057 | 0.194 |
| I155P | 0.372 | 0.043 | 0.714 | 0.143 |
| I155Q | 0.645 | 0.056 | 0.891 | 0.148 |
| I155R | 0.552 | 0.026 | 0.815 | 0.144 |
| I155E | 0.683 | 0.055 | 1.003 | 0.251 |
| I155K | 0.612 | 0.059 | 0.960 | 0.144 |
| I155D | 0.423 | 0.046 | 0.831 | 0.128 |
| L156I | 0.995 | 0.070 | 1.006 | 0.083 |
| L156F | 1.002 | 0.091 | 1.040 | 0.237 |
| L156V | 0.995 | 0.043 | 1.058 | 0.224 |
| L156C | 0.994 | 0.021 | 1.153 | 0.090 |
| L156M | 1.059 | 0.100 | 1.008 | 0.186 |
| L156A | 0.691 | 0.040 | 1.039 | 0.129 |
| L156W | 1.059 | 0.037 | 0.986 | 0.251 |
| L156T | 0.565 | 0.048 | 0.962 | 0.162 |
| L156Y | 0.977 | 0.046 | 0.913 | 0.304 |
| L156G | 0.442 | 0.036 | 0.704 | 0.075 |
| L156S | 0.600 | 0.053 | 0.931 | 0.242 |
| L156N | 0.554 | 0.026 | 0.786 | 0.093 |

| Variant ID | Average WT-SecM Score n = 3 | WT-SecM SD | Average P166A-SecM Score n = 3 | P166A-SecM SD |
|---|---|---|---|---|
| L156H | 0.701 | 0.054 | 1.030 | 0.194 |
| L156P | 0.453 | 0.059 | 0.724 | 0.164 |
| L156Q | 0.730 | 0.043 | 0.920 | 0.230 |
| L156R | 0.879 | 0.041 | 0.950 | 0.205 |
| L156E | 0.477 | 0.034 | 0.962 | 0.140 |
| L156K | 0.891 | 0.100 | 1.033 | 0.156 |
| L156D | 0.435 | 0.041 | 0.788 | 0.171 |
| E157I | 1.459 | 0.148 | 1.233 | 0.288 |
| E157L | 1.284 | 0.150 | 1.091 | 0.169 |
| E157F | 1.221 | 0.115 | 1.190 | 0.177 |
| E157V | 1.376 | 0.137 | 1.159 | 0.262 |
| E157C | 1.289 | 0.036 | 1.192 | 0.149 |
| E157A | 1.131 | 0.092 | 1.329 | 0.102 |
| E157W | 1.165 | 0.012 | 1.267 | 0.377 |
| E157T | 1.284 | 0.101 | 1.191 | 0.178 |
| E157Y | 1.129 | 0.093 | 0.916 | 0.117 |
| E157G | 0.884 | 0.053 | 1.005 | 0.089 |
| E157S | 1.169 | 0.080 | 1.029 | 0.136 |
| E157N | 1.205 | 0.079 | 1.094 | 0.182 |
| E157H | 1.119 | 0.070 | 0.782 | 0.268 |
| E157P | 0.412 | 0.072 | 0.637 | 0.171 |
| E157Q | 0.936 | 0.066 | 1.012 | 0.198 |
| E157R | 1.114 | 0.121 | 1.074 | 0.300 |
| E157K | 1.099 | 0.008 | 0.973 | 0.024 |
| E157D | 1.117 | 0.038 | 1.120 | 0.109 |
| R158I | 0.998 | 0.026 | 0.715 | 0.116 |
| R158L | 1.196 | 0.083 | 0.662 | 0.194 |
| R158F | 1.008 | 0.063 | 0.754 | 0.213 |
| R158V | 0.877 | 0.121 | 0.744 | 0.181 |
| R158C | 0.906 | 0.097 | 0.754 | 0.155 |
| R158M | 0.958 | 0.080 | 0.715 | 0.341 |
| R158A | 0.859 | 0.062 | 0.700 | 0.119 |
| R158W | 0.904 | 0.063 | 0.875 | 0.077 |
| R158T | 0.766 | 0.049 | 0.694 | 0.155 |
| R158Y | 0.999 | 0.089 | 0.738 | 0.221 |
| R158G | 0.644 | 0.036 | 0.874 | 0.060 |
| R158S | 0.832 | 0.061 | 0.686 | 0.104 |
| R158H | 0.954 | 0.081 | 0.773 | 0.235 |
| R158P | 0.695 | 0.041 | 0.734 | 0.286 |
| R158Q | 0.868 | 0.023 | 0.599 | 0.147 |
| R158E | 0.609 | 0.021 | 0.975 | 0.221 |
| R158K | 1.233 | 0.067 | 0.622 | 0.188 |
| R158D | 0.570 | 0.047 | 0.985 | 0.279 |

| Variant ID | Average WT-SecM Score n = 3 | WT-SecM SD | Average P166A-SecM Score n = 3 | P166A-SecM SD |
|---|---|---|---|---|
| R159I | 1.181 | 0.145 | 0.375 | 0.087 |
| R159L | 1.171 | 0.095 | 0.526 | 0.082 |
| R159V | 1.227 | 0.137 | 0.370 | 0.014 |
| R159A | 1.246 | 0.110 | 0.573 | 0.098 |
| R159W | 0.984 | 0.031 | 0.794 | 0.048 |
| R159T | 1.263 | 0.096 | 0.470 | 0.068 |
| R159G | 1.136 | 0.116 | 0.588 | 0.110 |
| R159S | 1.229 | 0.104 | 0.555 | 0.092 |
| R159H | 1.203 | 0.084 | 0.438 | 0.233 |
| R159P | 1.090 | 0.088 | 0.618 | 0.082 |
| R159Q | 1.064 | 0.058 | 0.801 | 0.042 |
| R159D | 0.952 | 0.068 | 0.687 | 0.099 |

**Table S2. List of Primers Used for The Production of Mutant Reporter Vectors**

| Primer Name | Primer Sequence (5′→3′) |
|---|---|
| DHFR-L56.rev | ACCAGACCCGCTCCGCCG |
| DHFR-L54.fwd | TCTGGAGCGGCGGAGCGGGTCTGGTGACCCAATTTATCAGAAAACGTTGGTGCCG |
| DHFR-L52.fwd | TCTGGAGCGGCGGAGCGGGTCTGGTATTTATCAGAAAACGTTGGTGCCGGGC |
| DHFR-L50.fwd | TCTGGAGCGGCGGAGCGGGTCTGGTCAGAAAACGTTGGTGCCGGGC |
| DHFR-L48.fwd | TCTGGAGCGGCGGAGCGGGTCTGGTACGTTGGTGCCGGGCCA |
| DHFR-L46.fwd | TCTGGAGCGGCGGAGCGGGTCTGGTGTGCCGGGCCAGCAAAACG |
| DHFR-L44.fwd | TCTGGAGCGGCGGAGCGGGTCTGGTGGCCAGCAAAACGCAACATGGAT |
| DHFR-L42.fwd | TCTGGAGCGGCGGAGCGGGTCTGGTCAAAACGCAACATGGATTGTACCCCCG |
| DHFR-L40.fwd | TCTGGAGCGGCGGAGCGGGTCTGGTGCAACATGGATTGTACCCCCGGG |
| DHFR-L46AL.fwd | CGTGCCGGTCCGTGAGGCAGCGGTTCT |
| DHFR-L46AL.rev | AGAACCGCTGCCTCACGGACCGGCACG |
| DHFR-L46FL.fwd | GGCATTCGTGCCGGTGCGGGCAGC |
| DHFR-L46FL.rev | GCTGCCCGCACCGGCACGAATGCC |
| DHFR-D27F.fwd | CGTGGAACCTGCCTGCCTTTCTCGCCTGGTTTAAAC |
| DHFR-D27F.rev | GTTTAAACCAGGCGAGAAAGGCAGGCAGGTTCCACG |
| DHFR-T113V.fwd | CAAAAGCGCAAAAACTGTATCTGGTTCATATCGACGCAGAAGTGGAAGG |
| DHFR-T113V.rev | CCTTCCACTTCTGCGTCGATATGAACCAGATACAGTTTTTGCGCTTTTG |
| DHFR-W133V.fwd | GATTACGAGCCGGATGACGTGGAATCGGTATTCAGCGA |
| DHFR-W133V.rev | TCGCTGAATACCGATTCCACGTCATCCGGCTCGTAATC |
