## Supplementary material for "Elucidation of Sequence Constraints for Protein Folding on the Ribosome": Document S1

DHFR-L56-SecM-(GS)_5_-GFP

ATGATCAGTCTGATTGCGGCGTTAGCGGTAGATCGCGTTATCGGCATGGAAAACGCCATGCCGTGGAACCTGCCTGCCGATCTCGCCTGGTTTAAACGCAACACCTTAAATAAACCCGTGATTATGGGCCGCCATACCTGGGAATCAATCGGTCGTCCGTTGCCAGGACGCAAAAATATTATCCTCAGCAGTCAACCGGGTACGGACGATCGCGTAACGTGGGTGAAGTCGGTGGATGAAGCCATCGCGGCGGCGGGTGACGTACCAGAAATCATGGTGATTGGCGGCGGTCGCGTTTATGAACAGTTCTTGCCAAAAGCGCAAAAACTGTATCTGACGCATATCGACGCAGAAGTGGAAGGCGACACCCATTTCCCGGATTACGAGCCGGATGACTGGGAATCGGTATTCAGCGAATTCCACGATGCTGATGCGCAGAACTCTCACAGCTATAGCTTTGAGATTCTGGAGCGGCGGAGCGGGTCTGGTATAAAAGACCCAATTTATCAGAAAACGTTGGTGCCGGGCCAGCAAAACGCAACATGGATTGTACCCCCGGGACAGTACTTTATGATGGGAGACTGGATGAGTTCCTTTAGCACCCCGGTTTGGATTAGCCAGGCGCAGGGCATTCGTGCCGGTCCGGGCAGCGGTTCTGGGTCCGGCTCGGGCAGTATGGCCACAACCATGACGGCCCTGACAGAAGGTGCGAAGCTGTTCGAGAAGGAGATTCCCTATATCACAGAATTGGAGGGGGATGTAGAGGGTATGAAGTTTATCATCAAAGGCGAAGGGACAGGGGATGCAACAACTGGAACAATTAAGGCTAAGTACATTTGCACGACCGGCGACGTCCCGGTGCCCTGGTCCACGCTCGTCACCACGCTCACGTACGGAGCCCAGTGCTTTGCCAAATATGGCCCTGAACTTAAAGACTTCTACAAGTCGTGTATGCCGGAGGGATACGTGCAAGAGAGGACGATCACCTTTGAAGGTGACGGAGTATTCAAAACAAGAGCGGAGGTGACGTTCGAGAATGGATCGGTCTATAACCGGGTCAAGCTCAACGGACAGGGCTTTAAGAAAGATGGACACGTCCTTGGGAAGAATTTGGAGTTCAATTTCACCCCGCATTGTCTTTACATCTGGGGTGATCAGGCGAATCACGGGTTGAAATCAGCGTTCAAGATCATGCACGAGATTACGGGGAGCAAAGAGGACTTTATCGTGGCAGACCACACTCAGATGAACACTCCAATCGGAGGGGGTCCCGTACACGTACCCGAGTATCATCACCTGACCGTCTGGACATCGTTTGGAAAAGACCCTGACGACGATGAAACTGATCATCTCAACATTGTGGAAGTGATCAAGGCGGTGGACTTGGAAACATACCGGTGA

DHFR-L54-SecM-(GS)_5_-GFP

ATGATCAGTCTGATTGCGGCGTTAGCGGTAGATCGCGTTATCGGCATGGAAAACGCCATGCCGTGGAACCTGCCTGCCGATCTCGCCTGGTTTAAACGCAACACCTTAAATAAACCCGTGATTATGGGCCGCCATACCTGGGAATCAATCGGTCGTCCGTTGCCAGGACGCAAAAATATTATCCTCAGCAGTCAACCGGGTACGGACGATCGCGTAACGTGGGTGAAGTCGGTGGATGAAGCCATCGCGGCGGCGGGTGACGTACCAGAAATCATGGTGATTGGCGGCGGTCGCGTTTATGAACAGTTCTTGCCAAAAGCGCAAAAACTGTATCTGACGCATATCGACGCAGAAGTGGAAGGCGACACCCATTTCCCGGATTACGAGCCGGATGACTGGGAATCGGTATTCAGCGAATTCCACGATGCTGATGCGCAGAACTCTCACAGCTATAGCTTTGAGATTCTGGAGCGGCGGAGCGGGTCTGGTGACCCAATTTATCAGAAAACGTTGGTGCCGGGCCAGCAAAACGCAACATGGATTGTACCCCCGGGACAGTACTTTATGATGGGAGACTGGATGAGTTCCTTTAGCACCCCGGTTTGGATTAGCCAGGCGCAGGGCATTCGTGCCGGTCCGGGCAGCGGTTCTGGGTCCGGCTCGGGCAGTATGGCCACAACCATGACGGCCCTGACAGAAGGTGCGAAGCTGTTCGAGAAGGAGATTCCCTATATCACAGAATTGGAGGGGGATGTAGAGGGTATGAAGTTTATCATCAAAGGCGAAGGGACAGGGGATGCAACAACTGGAACAATTAAGGCTAAGTACATTTGCACGACCGGCGACGTCCCGGTGCCCTGGTCCACGCTCGTCACCACGCTCACGTACGGAGCCCAGTGCTTTGCCAAATATGGCCCTGAACTTAAAGACTTCTACAAGTCGTGTATGCCGGAGGGATACGTGCAAGAGAGGACGATCACCTTTGAAGGTGACGGAGTATTCAAAACAAGAGCGGAGGTGACGTTCGAGAATGGATCGGTCTATAACCGGGTCAAGCTCAACGGACAGGGCTTTAAGAAAGATGGACACGTCCTTGGGAAGAATTTGGAGTTCAATTTCACCCCGCATTGTCTTTACATCTGGGGTGATCAGGCGAATCACGGGTTGAAATCAGCGTTCAAGATCATGCACGAGATTACGGGGAGCAAAGAGGACTTTATCGTGGCAGACCACACTCAGATGAACACTCCAATCGGAGGGGGTCCCGTACACGTACCCGAGTATCATCACCTGACCGTCTGGACATCGTTTGGAAAAGACCCTGACGACGATGAAACTGATCATCTCAACATTGTGGAAGTGATCAAGGCGGTGGACTTGGAAACATACCGGTGA

DHFR-L52-SecM-(GS)_5_-GFP

ATGATCAGTCTGATTGCGGCGTTAGCGGTAGATCGCGTTATCGGCATGGAAAACGCCATGCCGTGGAACCTGCCTGCCGATCTCGCCTGGTTTAAACGCAACACCTTAAATAAACCCGTGATTATGGGCCGCCATACCTGGGAATCAATCGGTCGTCCGTTGCCAGGACGCAAAAATATTATCCTCAGCAGTCAACCGGGTACGGACGATCGCGTAACGTGGGTGAAGTCGGTGGATGAAGCCATCGCGGCGGCGGGTGACGTACCAGAAATCATGGTGATTGGCGGCGGTCGCGTTTATGAACAGTTCTTGCCAAAAGCGCAAAAACTGTATCTGACGCATATCGACGCAGAAGTGGAAGGCGACACCCATTTCCCGGATTACGAGCCGGATGACTGGGAATCGGTATTCAGCGAATTCCACGATGCTGATGCGCAGAACTCTCACAGCTATAGCTTTGAGATTCTGGAGCGGCGGAGCGGGTCTGGTATTTATCAGAAAACGTTGGTGCCGGGCCAGCAAAACGCAACATGGATTGTACCCCCGGGACAGTACTTTATGATGGGAGACTGGATGAGTTCCTTTAGCACCCCGGTTTGGATTAGCCAGGCGCAGGGCATTCGTGCCGGTCCGGGCAGCGGTTCTGGGTCCGGCTCGGGCAGTATGGCCACAACCATGACGGCCCTGACAGAAGGTGCGAAGCTGTTCGAGAAGGAGATTCCCTATATCACAGAATTGGAGGGGGATGTAGAGGGTATGAAGTTTATCATCAAAGGCGAAGGGACAGGGGATGCAACAACTGGAACAATTAAGGCTAAGTACATTTGCACGACCGGCGACGTCCCGGTGCCCTGGTCCACGCTCGTCACCACGCTCACGTACGGAGCCCAGTGCTTTGCCAAATATGGCCCTGAACTTAAAGACTTCTACAAGTCGTGTATGCCGGAGGGATACGTGCAAGAGAGGACGATCACCTTTGAAGGTGACGGAGTATTCAAAACAAGAGCGGAGGTGACGTTCGAGAATGGATCGGTCTATAACCGGGTCAAGCTCAACGGACAGGGCTTTAAGAAAGATGGACACGTCCTTGGGAAGAATTTGGAGTTCAATTTCACCCCGCATTGTCTTTACATCTGGGGTGATCAGGCGAATCACGGGTTGAAATCAGCGTTCAAGATCATGCACGAGATTACGGGGAGCAAAGAGGACTTTATCGTGGCAGACCACACTCAGATGAACACTCCAATCGGAGGGGGTCCCGTACACGTACCCGAGTATCATCACCTGACCGTCTGGACATCGTTTGGAAAAGACCCTGACGACGATGAAACTGATCATCTCAACATTGTGGAAGTGATCAAGGCGGTGGACTTGGAAACATACCGGTGA

DHFR-L50-SecM-(GS)_5_-GFP

ATGATCAGTCTGATTGCGGCGTTAGCGGTAGATCGCGTTATCGGCATGGAAAACGCCATGCCGTGGAACCTGCCTGCCGATCTCGCCTGGTTTAAACGCAACACCTTAAATAAACCCGTGATTATGGGCCGCCATACCTGGGAATCAATCGGTCGTCCGTTGCCAGGACGCAAAAATATTATCCTCAGCAGTCAACCGGGTACGGACGATCGCGTAACGTGGGTGAAGTCGGTGGATGAAGCCATCGCGGCGGCGGGTGACGTACCAGAAATCATGGTGATTGGCGGCGGTCGCGTTTATGAACAGTTCTTGCCAAAAGCGCAAAAACTGTATCTGACGCATATCGACGCAGAAGTGGAAGGCGACACCCATTTCCCGGATTACGAGCCGGATGACTGGGAATCGGTATTCAGCGAATTCCACGATGCTGATGCGCAGAACTCTCACAGCTATAGCTTTGAGATTCTGGAGCGGCGGAGCGGGTCTGGTCAGAAAACGTTGGTGCCGGGCCAGCAAAACGCAACATGGATTGTACCCCCGGGACAGTACTTTATGATGGGAGACTGGATGAGTTCCTTTAGCACCCCGGTTTGGATTAGCCAGGCGCAGGGCATTCGTGCCGGTCCGGGCAGCGGTTCTGGGTCCGGCTCGGGCAGTATGGCCACAACCATGACGGCCCTGACAGAAGGTGCGAAGCTGTTCGAGAAGGAGATTCCCTATATCACAGAATTGGAGGGGGATGTAGAGGGTATGAAGTTTATCATCAAAGGCGAAGGGACAGGGGATGCAACAACTGGAACAATTAAGGCTAAGTACATTTGCACGACCGGCGACGTCCCGGTGCCCTGGTCCACGCTCGTCACCACGCTCACGTACGGAGCCCAGTGCTTTGCCAAATATGGCCCTGAACTTAAAGACTTCTACAAGTCGTGTATGCCGGAGGGATACGTGCAAGAGAGGACGATCACCTTTGAAGGTGACGGAGTATTCAAAACAAGAGCGGAGGTGACGTTCGAGAATGGATCGGTCTATAACCGGGTCAAGCTCAACGGACAGGGCTTTAAGAAAGATGGACACGTCCTTGGGAAGAATTTGGAGTTCAATTTCACCCCGCATTGTCTTTACATCTGGGGTGATCAGGCGAATCACGGGTTGAAATCAGCGTTCAAGATCATGCACGAGATTACGGGGAGCAAAGAGGACTTTATCGTGGCAGACCACACTCAGATGAACACTCCAATCGGAGGGGGTCCCGTACACGTACCCGAGTATCATCACCTGACCGTCTGGACATCGTTTGGAAAAGACCCTGACGACGATGAAACTGATCATCTCAACATTGTGGAAGTGATCAAGGCGGTGGACTTGGAAACATACCGGTGA

DHFR-L48-SecM-(GS)_5_-GFP

ATGATCAGTCTGATTGCGGCGTTAGCGGTAGATCGCGTTATCGGCATGGAAAACGCCATGCCGTGGAACCTGCCTGCCGATCTCGCCTGGTTTAAACGCAACACCTTAAATAAACCCGTGATTATGGGCCGCCATACCTGGGAATCAATCGGTCGTCCGTTGCCAGGACGCAAAAATATTATCCTCAGCAGTCAACCGGGTACGGACGATCGCGTAACGTGGGTGAAGTCGGTGGATGAAGCCATCGCGGCGGCGGGTGACGTACCAGAAATCATGGTGATTGGCGGCGGTCGCGTTTATGAACAGTTCTTGCCAAAAGCGCAAAAACTGTATCTGACGCATATCGACGCAGAAGTGGAAGGCGACACCCATTTCCCGGATTACGAGCCGGATGACTGGGAATCGGTATTCAGCGAATTCCACGATGCTGATGCGCAGAACTCTCACAGCTATAGCTTTGAGATTCTGGAGCGGCGGAGCGGGTCTGGTACGTTGGTGCCGGGCCAGCAAAACGCAACATGGATTGTACCCCCGGGACAGTACTTTATGATGGGAGACTGGATGAGTTCCTTTAGCACCCCGGTTTGGATTAGCCAGGCGCAGGGCATTCGTGCCGGTCCGGGCAGCGGTTCTGGGTCCGGCTCGGGCAGTATGGCCACAACCATGACGGCCCTGACAGAAGGTGCGAAGCTGTTCGAGAAGGAGATTCCCTATATCACAGAATTGGAGGGGGATGTAGAGGGTATGAAGTTTATCATCAAAGGCGAAGGGACAGGGGATGCAACAACTGGAACAATTAAGGCTAAGTACATTTGCACGACCGGCGACGTCCCGGTGCCCTGGTCCACGCTCGTCACCACGCTCACGTACGGAGCCCAGTGCTTTGCCAAATATGGCCCTGAACTTAAAGACTTCTACAAGTCGTGTATGCCGGAGGGATACGTGCAAGAGAGGACGATCACCTTTGAAGGTGACGGAGTATTCAAAACAAGAGCGGAGGTGACGTTCGAGAATGGATCGGTCTATAACCGGGTCAAGCTCAACGGACAGGGCTTTAAGAAAGATGGACACGTCCTTGGGAAGAATTTGGAGTTCAATTTCACCCCGCATTGTCTTTACATCTGGGGTGATCAGGCGAATCACGGGTTGAAATCAGCGTTCAAGATCATGCACGAGATTACGGGGAGCAAAGAGGACTTTATCGTGGCAGACCACACTCAGATGAACACTCCAATCGGAGGGGGTCCCGTACACGTACCCGAGTATCATCACCTGACCGTCTGGACATCGTTTGGAAAAGACCCTGACGACGATGAAACTGATCATCTCAACATTGTGGAAGTGATCAAGGCGGTGGACTTGGAAACATACCGGTGA

DHFR-L46-SecM-(GS)_5_-GFP

ATGATCAGTCTGATTGCGGCGTTAGCGGTAGATCGCGTTATCGGCATGGAAAACGCCATGCCGTGGAACCTGCCTGCCGATCTCGCCTGGTTTAAACGCAACACCTTAAATAAACCCGTGATTATGGGCCGCCATACCTGGGAATCAATCGGTCGTCCGTTGCCAGGACGCAAAAATATTATCCTCAGCAGTCAACCGGGTACGGACGATCGCGTAACGTGGGTGAAGTCGGTGGATGAAGCCATCGCGGCGGCGGGTGACGTACCAGAAATCATGGTGATTGGCGGCGGTCGCGTTTATGAACAGTTCTTGCCAAAAGCGCAAAAACTGTATCTGACGCATATCGACGCAGAAGTGGAAGGCGACACCCATTTCCCGGATTACGAGCCGGATGACTGGGAATCGGTATTCAGCGAATTCCACGATGCTGATGCGCAGAACTCTCACAGCTATAGCTTTGAGATTCTGGAGCGGCGGAGCGGGTCTGGTGTGCCGGGCCAGCAAAACGCAACATGGATTGTACCCCCGGGACAGTACTTTATGATGGGAGACTGGATGAGTTCCTTTAGCACCCCGGTTTGGATTAGCCAGGCGCAGGGCATTCGTGCCGGTCCGGGCAGCGGTTCTGGGTCCGGCTCGGGCAGTATGGCCACAACCATGACGGCCCTGACAGAAGGTGCGAAGCTGTTCGAGAAGGAGATTCCCTATATCACAGAATTGGAGGGGGATGTAGAGGGTATGAAGTTTATCATCAAAGGCGAAGGGACAGGGGATGCAACAACTGGAACAATTAAGGCTAAGTACATTTGCACGACCGGCGACGTCCCGGTGCCCTGGTCCACGCTCGTCACCACGCTCACGTACGGAGCCCAGTGCTTTGCCAAATATGGCCCTGAACTTAAAGACTTCTACAAGTCGTGTATGCCGGAGGGATACGTGCAAGAGAGGACGATCACCTTTGAAGGTGACGGAGTATTCAAAACAAGAGCGGAGGTGACGTTCGAGAATGGATCGGTCTATAACCGGGTCAAGCTCAACGGACAGGGCTTTAAGAAAGATGGACACGTCCTTGGGAAGAATTTGGAGTTCAATTTCACCCCGCATTGTCTTTACATCTGGGGTGATCAGGCGAATCACGGGTTGAAATCAGCGTTCAAGATCATGCACGAGATTACGGGGAGCAAAGAGGACTTTATCGTGGCAGACCACACTCAGATGAACACTCCAATCGGAGGGGGTCCCGTACACGTACCCGAGTATCATCACCTGACCGTCTGGACATCGTTTGGAAAAGACCCTGACGACGATGAAACTGATCATCTCAACATTGTGGAAGTGATCAAGGCGGTGGACTTGGAAACATACCGGTGA

DHFR-L44-SecM-(GS)_5_-GFP

ATGATCAGTCTGATTGCGGCGTTAGCGGTAGATCGCGTTATCGGCATGGAAAACGCCATGCCGTGGAACCTGCCTGCCGATCTCGCCTGGTTTAAACGCAACACCTTAAATAAACCCGTGATTATGGGCCGCCATACCTGGGAATCAATCGGTCGTCCGTTGCCAGGACGCAAAAATATTATCCTCAGCAGTCAACCGGGTACGGACGATCGCGTAACGTGGGTGAAGTCGGTGGATGAAGCCATCGCGGCGGCGGGTGACGTACCAGAAATCATGGTGATTGGCGGCGGTCGCGTTTATGAACAGTTCTTGCCAAAAGCGCAAAAACTGTATCTGACGCATATCGACGCAGAAGTGGAAGGCGACACCCATTTCCCGGATTACGAGCCGGATGACTGGGAATCGGTATTCAGCGAATTCCACGATGCTGATGCGCAGAACTCTCACAGCTATAGCTTTGAGATTCTGGAGCGGCGGAGCGGGTCTGGTGGCCAGCAAAACGCAACATGGATTGTACCCCCGGGACAGTACTTTATGATGGGAGACTGGATGAGTTCCTTTAGCACCCCGGTTTGGATTAGCCAGGCGCAGGGCATTCGTGCCGGTCCGGGCAGCGGTTCTGGGTCCGGCTCGGGCAGTATGGCCACAACCATGACGGCCCTGACAGAAGGTGCGAAGCTGTTCGAGAAGGAGATTCCCTATATCACAGAATTGGAGGGGGATGTAGAGGGTATGAAGTTTATCATCAAAGGCGAAGGGACAGGGGATGCAACAACTGGAACAATTAAGGCTAAGTACATTTGCACGACCGGCGACGTCCCGGTGCCCTGGTCCACGCTCGTCACCACGCTCACGTACGGAGCCCAGTGCTTTGCCAAATATGGCCCTGAACTTAAAGACTTCTACAAGTCGTGTATGCCGGAGGGATACGTGCAAGAGAGGACGATCACCTTTGAAGGTGACGGAGTATTCAAAACAAGAGCGGAGGTGACGTTCGAGAATGGATCGGTCTATAACCGGGTCAAGCTCAACGGACAGGGCTTTAAGAAAGATGGACACGTCCTTGGGAAGAATTTGGAGTTCAATTTCACCCCGCATTGTCTTTACATCTGGGGTGATCAGGCGAATCACGGGTTGAAATCAGCGTTCAAGATCATGCACGAGATTACGGGGAGCAAAGAGGACTTTATCGTGGCAGACCACACTCAGATGAACACTCCAATCGGAGGGGGTCCCGTACACGTACCCGAGTATCATCACCTGACCGTCTGGACATCGTTTGGAAAAGACCCTGACGACGATGAAACTGATCATCTCAACATTGTGGAAGTGATCAAGGCGGTGGACTTGGAAACATACCGGTGA

DHFR-L42-SecM-(GS)_5_-GFP

ATGATCAGTCTGATTGCGGCGTTAGCGGTAGATCGCGTTATCGGCATGGAAAACGCCATGCCGTGGAACCTGCCTGCCGATCTCGCCTGGTTTAAACGCAACACCTTAAATAAACCCGTGATTATGGGCCGCCATACCTGGGAATCAATCGGTCGTCCGTTGCCAGGACGCAAAAATATTATCCTCAGCAGTCAACCGGGTACGGACGATCGCGTAACGTGGGTGAAGTCGGTGGATGAAGCCATCGCGGCGGCGGGTGACGTACCAGAAATCATGGTGATTGGCGGCGGTCGCGTTTATGAACAGTTCTTGCCAAAAGCGCAAAAACTGTATCTGACGCATATCGACGCAGAAGTGGAAGGCGACACCCATTTCCCGGATTACGAGCCGGATGACTGGGAATCGGTATTCAGCGAATTCCACGATGCTGATGCGCAGAACTCTCACAGCTATAGCTTTGAGATTCTGGAGCGGCGGAGCGGGTCTGGTCAAAACGCAACATGGATTGTACCCCCGGGACAGTACTTTATGATGGGAGACTGGATGAGTTCCTTTAGCACCCCGGTTTGGATTAGCCAGGCGCAGGGCATTCGTGCCGGTCCGGGCAGCGGTTCTGGGTCCGGCTCGGGCAGTATGGCCACAACCATGACGGCCCTGACAGAAGGTGCGAAGCTGTTCGAGAAGGAGATTCCCTATATCACAGAATTGGAGGGGGATGTAGAGGGTATGAAGTTTATCATCAAAGGCGAAGGGACAGGGGATGCAACAACTGGAACAATTAAGGCTAAGTACATTTGCACGACCGGCGACGTCCCGGTGCCCTGGTCCACGCTCGTCACCACGCTCACGTACGGAGCCCAGTGCTTTGCCAAATATGGCCCTGAACTTAAAGACTTCTACAAGTCGTGTATGCCGGAGGGATACGTGCAAGAGAGGACGATCACCTTTGAAGGTGACGGAGTATTCAAAACAAGAGCGGAGGTGACGTTCGAGAATGGATCGGTCTATAACCGGGTCAAGCTCAACGGACAGGGCTTTAAGAAAGATGGACACGTCCTTGGGAAGAATTTGGAGTTCAATTTCACCCCGCATTGTCTTTACATCTGGGGTGATCAGGCGAATCACGGGTTGAAATCAGCGTTCAAGATCATGCACGAGATTACGGGGAGCAAAGAGGACTTTATCGTGGCAGACCACACTCAGATGAACACTCCAATCGGAGGGGGTCCCGTACACGTACCCGAGTATCATCACCTGACCGTCTGGACATCGTTTGGAAAAGACCCTGACGACGATGAAACTGATCATCTCAACATTGTGGAAGTGATCAAGGCGGTGGACTTGGAAACATACCGGTGA

DHFR-L40-SecM-(GS)_5_-GFP

ATGATCAGTCTGATTGCGGCGTTAGCGGTAGATCGCGTTATCGGCATGGAAAACGCCATGCCGTGGAACCTGCCTGCCGATCTCGCCTGGTTTAAACGCAACACCTTAAATAAACCCGTGATTATGGGCCGCCATACCTGGGAATCAATCGGTCGTCCGTTGCCAGGACGCAAAAATATTATCCTCAGCAGTCAACCGGGTACGGACGATCGCGTAACGTGGGTGAAGTCGGTGGATGAAGCCATCGCGGCGGCGGGTGACGTACCAGAAATCATGGTGATTGGCGGCGGTCGCGTTTATGAACAGTTCTTGCCAAAAGCGCAAAAACTGTATCTGACGCATATCGACGCAGAAGTGGAAGGCGACACCCATTTCCCGGATTACGAGCCGGATGACTGGGAATCGGTATTCAGCGAATTCCACGATGCTGATGCGCAGAACTCTCACAGCTATAGCTTTGAGATTCTGGAGCGGCGGAGCGGGTCTGGTGCAACATGGATTGTACCCCCGGGACAGTACTTTATGATGGGAGACTGGATGAGTTCCTTTAGCACCCCGGTTTGGATTAGCCAGGCGCAGGGCATTCGTGCCGGTCCGGGCAGCGGTTCTGGGTCCGGCTCGGGCAGTATGGCCACAACCATGACGGCCCTGACAGAAGGTGCGAAGCTGTTCGAGAAGGAGATTCCCTATATCACAGAATTGGAGGGGGATGTAGAGGGTATGAAGTTTATCATCAAAGGCGAAGGGACAGGGGATGCAACAACTGGAACAATTAAGGCTAAGTACATTTGCACGACCGGCGACGTCCCGGTGCCCTGGTCCACGCTCGTCACCACGCTCACGTACGGAGCCCAGTGCTTTGCCAAATATGGCCCTGAACTTAAAGACTTCTACAAGTCGTGTATGCCGGAGGGATACGTGCAAGAGAGGACGATCACCTTTGAAGGTGACGGAGTATTCAAAACAAGAGCGGAGGTGACGTTCGAGAATGGATCGGTCTATAACCGGGTCAAGCTCAACGGACAGGGCTTTAAGAAAGATGGACACGTCCTTGGGAAGAATTTGGAGTTCAATTTCACCCCGCATTGTCTTTACATCTGGGGTGATCAGGCGAATCACGGGTTGAAATCAGCGTTCAAGATCATGCACGAGATTACGGGGAGCAAAGAGGACTTTATCGTGGCAGACCACACTCAGATGAACACTCCAATCGGAGGGGGTCCCGTACACGTACCCGAGTATCATCACCTGACCGTCTGGACATCGTTTGGAAAAGACCCTGACGACGATGAAACTGATCATCTCAACATTGTGGAAGTGATCAAGGCGGTGGACTTGGAAACATACCGGTGA

DHFR-L46-SecM-**P166A**-(GS)_5_-GFP (Full Length Control)

ATGATCAGTCTGATTGCGGCGTTAGCGGTAGATCGCGTTATCGGCATGGAAAACGCCATGCCGTGGAACCTGCCTGCCGATCTCGCCTGGTTTAAACGCAACACCTTAAATAAACCCGTGATTATGGGCCGCCATACCTGGGAATCAATCGGTCGTCCGTTGCCAGGACGCAAAAATATTATCCTCAGCAGTCAACCGGGTACGGACGATCGCGTAACGTGGGTGAAGTCGGTGGATGAAGCCATCGCGGCGGCGGGTGACGTACCAGAAATCATGGTGATTGGCGGCGGTCGCGTTTATGAACAGTTCTTGCCAAAAGCGCAAAAACTGTATCTGACGCATATCGACGCAGAAGTGGAAGGCGACACCCATTTCCCGGATTACGAGCCGGATGACTGGGAATCGGTATTCAGCGAATTCCACGATGCTGATGCGCAGAACTCTCACAGCTATAGCTTTGAGATTCTGGAGCGGCGGAGCGGGTCTGGTGTGCCGGGCCAGCAAAACGCAACATGGATTGTACCCCCGGGACAGTACTTTATGATGGGAGACTGGATGAGTTCCTTTAGCACCCCGGTTTGGATTAGCCAGGCGCAGGGCATTCGTGCCGGT**GCG**GGCAGCGGTTCTGGGTCCGGCTCGGGCAGTATGGCCACAACCATGACGGCCCTGACAGAAGGTGCGAAGCTGTTCGAGAAGGAGATTCCCTATATCACAGAATTGGAGGGGGATGTAGAGGGTATGAAGTTTATCATCAAAGGCGAAGGGACAGGGGATGCAACAACTGGAACAATTAAGGCTAAGTACATTTGCACGACCGGCGACGTCCCGGTGCCCTGGTCCACGCTCGTCACCACGCTCACGTACGGAGCCCAGTGCTTTGCCAAATATGGCCCTGAACTTAAAGACTTCTACAAGTCGTGTATGCCGGAGGGATACGTGCAAGAGAGGACGATCACCTTTGAAGGTGACGGAGTATTCAAAACAAGAGCGGAGGTGACGTTCGAGAATGGATCGGTCTATAACCGGGTCAAGCTCAACGGACAGGGCTTTAAGAAAGATGGACACGTCCTTGGGAAGAATTTGGAGTTCAATTTCACCCCGCATTGTCTTTACATCTGGGGTGATCAGGCGAATCACGGGTTGAAATCAGCGTTCAAGATCATGCACGAGATTACGGGGAGCAAAGAGGACTTTATCGTGGCAGACCACACTCAGATGAACACTCCAATCGGAGGGGGTCCCGTACACGTACCCGAGTATCATCACCTGACCGTCTGGACATCGTTTGGAAAAGACCCTGACGACGATGAAACTGATCATCTCAACATTGTGGAAGTGATCAAGGCGGTGGACTTGGAAACATACCGGTGA

DHFR-L46-SecM-**STOP**-(GS)_5_-GFP (Arrested Length Control)

ATGATCAGTCTGATTGCGGCGTTAGCGGTAGATCGCGTTATCGGCATGGAAAACGCCATGCCGTGGAACCTGCCTGCCGATCTCGCCTGGTTTAAACGCAACACCTTAAATAAACCCGTGATTATGGGCCGCCATACCTGGGAATCAATCGGTCGTCCGTTGCCAGGACGCAAAAATATTATCCTCAGCAGTCAACCGGGTACGGACGATCGCGTAACGTGGGTGAAGTCGGTGGATGAAGCCATCGCGGCGGCGGGTGACGTACCAGAAATCATGGTGATTGGCGGCGGTCGCGTTTATGAACAGTTCTTGCCAAAAGCGCAAAAACTGTATCTGACGCATATCGACGCAGAAGTGGAAGGCGACACCCATTTCCCGGATTACGAGCCGGATGACTGGGAATCGGTATTCAGCGAATTCCACGATGCTGATGCGCAGAACTCTCACAGCTATAGCTTTGAGATTCTGGAGCGGCGGAGCGGGTCTGGTGTGCCGGGCCAGCAAAACGCAACATGGATTGTACCCCCGGGACAGTACTTTATGATGGGAGACTGGATGAGTTCCTTTAGCACCCCGGTTTGGATTAGCCAGGCGCAGGGCATTCGTGCCGGTCCG**TGA**GGCAGCGGTTCTGGGTCCGGCTCGGGCAGTATGGCCACAACCATGACGGCCCTGACAGAAGGTGCGAAGCTGTTCGAGAAGGAGATTCCCTATATCACAGAATTGGAGGGGGATGTAGAGGGTATGAAGTTTATCATCAAAGGCGAAGGGACAGGGGATGCAACAACTGGAACAATTAAGGCTAAGTACATTTGCACGACCGGCGACGTCCCGGTGCCCTGGTCCACGCTCGTCACCACGCTCACGTACGGAGCCCAGTGCTTTGCCAAATATGGCCCTGAACTTAAAGACTTCTACAAGTCGTGTATGCCGGAGGGATACGTGCAAGAGAGGACGATCACCTTTGAAGGTGACGGAGTATTCAAAACAAGAGCGGAGGTGACGTTCGAGAATGGATCGGTCTATAACCGGGTCAAGCTCAACGGACAGGGCTTTAAGAAAGATGGACACGTCCTTGGGAAGAATTTGGAGTTCAATTTCACCCCGCATTGTCTTTACATCTGGGGTGATCAGGCGAATCACGGGTTGAAATCAGCGTTCAAGATCATGCACGAGATTACGGGGAGCAAAGAGGACTTTATCGTGGCAGACCACACTCAGATGAACACTCCAATCGGAGGGGGTCCCGTACACGTACCCGAGTATCATCACCTGACCGTCTGGACATCGTTTGGAAAAGACCCTGACGACGATGAAACTGATCATCTCAACATTGTGGAAGTGATCAAGGCGGTGGACTTGGAAACATACCGGTGA

**D27F**-DHFR-L46-SecM-(GS)_5_-GFP

ATGATCAGTCTGATTGCGGCGTTAGCGGTAGATCGCGTTATCGGCATGGAAAACGCCATGCCGTGGAACCTGCCTGCC**TTT**CTCGCCTGGTTTAAACGCAACACCTTAAATAAACCCGTGATTATGGGCCGCCATACCTGGGAATCAATCGGTCGTCCGTTGCCAGGACGCAAAAATATTATCCTCAGCAGTCAACCGGGTACGGACGATCGCGTAACGTGGGTGAAGTCGGTGGATGAAGCCATCGCGGCGGCGGGTGACGTACCAGAAATCATGGTGATTGGCGGCGGTCGCGTTTATGAACAGTTCTTGCCAAAAGCGCAAAAACTGTATCTGACGCATATCGACGCAGAAGTGGAAGGCGACACCCATTTCCCGGATTACGAGCCGGATGACTGGGAATCGGTATTCAGCGAATTCCACGATGCTGATGCGCAGAACTCTCACAGCTATAGCTTTGAGATTCTGGAGCGGCGGAGCGGGTCTGGTGTGCCGGGCCAGCAAAACGCAACATGGATTGTACCCCCGGGACAGTACTTTATGATGGGAGACTGGATGAGTTCCTTTAGCACCCCGGTTTGGATTAGCCAGGCGCAGGGCATTCGTGCCGGTCCGGGCAGCGGTTCTGGGTCCGGCTCGGGCAGTATGGCCACAACCATGACGGCCCTGACAGAAGGTGCGAAGCTGTTCGAGAAGGAGATTCCCTATATCACAGAATTGGAGGGGGATGTAGAGGGTATGAAGTTTATCATCAAAGGCGAAGGGACAGGGGATGCAACAACTGGAACAATTAAGGCTAAGTACATTTGCACGACCGGCGACGTCCCGGTGCCCTGGTCCACGCTCGTCACCACGCTCACGTACGGAGCCCAGTGCTTTGCCAAATATGGCCCTGAACTTAAAGACTTCTACAAGTCGTGTATGCCGGAGGGATACGTGCAAGAGAGGACGATCACCTTTGAAGGTGACGGAGTATTCAAAACAAGAGCGGAGGTGACGTTCGAGAATGGATCGGTCTATAACCGGGTCAAGCTCAACGGACAGGGCTTTAAGAAAGATGGACACGTCCTTGGGAAGAATTTGGAGTTCAATTTCACCCCGCATTGTCTTTACATCTGGGGTGATCAGGCGAATCACGGGTTGAAATCAGCGTTCAAGATCATGCACGAGATTACGGGGAGCAAAGAGGACTTTATCGTGGCAGACCACACTCAGATGAACACTCCAATCGGAGGGGGTCCCGTACACGTACCCGAGTATCATCACCTGACCGTCTGGACATCGTTTGGAAAAGACCCTGACGACGATGAAACTGATCATCTCAACATTGTGGAAGTGATCAAGGCGGTGGACTTGGAAACATACCGGTGA

**T113V**-DHFR-L46-SecM-(GS)_5_-GFP

ATGATCAGTCTGATTGCGGCGTTAGCGGTAGATCGCGTTATCGGCATGGAAAACGCCATGCCGTGGAACCTGCCTGCCGATCTCGCCTGGTTTAAACGCAACACCTTAAATAAACCCGTGATTATGGGCCGCCATACCTGGGAATCAATCGGTCGTCCGTTGCCAGGACGCAAAAATATTATCCTCAGCAGTCAACCGGGTACGGACGATCGCGTAACGTGGGTGAAGTCGGTGGATGAAGCCATCGCGGCGGCGGGTGACGTACCAGAAATCATGGTGATTGGCGGCGGTCGCGTTTATGAACAGTTCTTGCCAAAAGCGCAAAAACTGTATCTG**GTT**CATATCGACGCAGAAGTGGAAGGCGACACCCATTTCCCGGATTACGAGCCGGATGACTGGGAATCGGTATTCAGCGAATTCCACGATGCTGATGCGCAGAACTCTCACAGCTATAGCTTTGAGATTCTGGAGCGGCGGAGCGGGTCTGGTGTGCCGGGCCAGCAAAACGCAACATGGATTGTACCCCCGGGACAGTACTTTATGATGGGAGACTGGATGAGTTCCTTTAGCACCCCGGTTTGGATTAGCCAGGCGCAGGGCATTCGTGCCGGTCCGGGCAGCGGTTCTGGGTCCGGCTCGGGCAGTATGGCCACAACCATGACGGCCCTGACAGAAGGTGCGAAGCTGTTCGAGAAGGAGATTCCCTATATCACAGAATTGGAGGGGGATGTAGAGGGTATGAAGTTTATCATCAAAGGCGAAGGGACAGGGGATGCAACAACTGGAACAATTAAGGCTAAGTACATTTGCACGACCGGCGACGTCCCGGTGCCCTGGTCCACGCTCGTCACCACGCTCACGTACGGAGCCCAGTGCTTTGCCAAATATGGCCCTGAACTTAAAGACTTCTACAAGTCGTGTATGCCGGAGGGATACGTGCAAGAGAGGACGATCACCTTTGAAGGTGACGGAGTATTCAAAACAAGAGCGGAGGTGACGTTCGAGAATGGATCGGTCTATAACCGGGTCAAGCTCAACGGACAGGGCTTTAAGAAAGATGGACACGTCCTTGGGAAGAATTTGGAGTTCAATTTCACCCCGCATTGTCTTTACATCTGGGGTGATCAGGCGAATCACGGGTTGAAATCAGCGTTCAAGATCATGCACGAGATTACGGGGAGCAAAGAGGACTTTATCGTGGCAGACCACACTCAGATGAACACTCCAATCGGAGGGGGTCCCGTACACGTACCCGAGTATCATCACCTGACCGTCTGGACATCGTTTGGAAAAGACCCTGACGACGATGAAACTGATCATCTCAACATTGTGGAAGTGATCAAGGCGGTGGACTTGGAAACATACCGGTGA

**W133V**-DHFR-L46-SecM-(GS)_5_-GFP

ATGATCAGTCTGATTGCGGCGTTAGCGGTAGATCGCGTTATCGGCATGGAAAACGCCATGCCGTGGAACCTGCCTGCCGATCTCGCCTGGTTTAAACGCAACACCTTAAATAAACCCGTGATTATGGGCCGCCATACCTGGGAATCAATCGGTCGTCCGTTGCCAGGACGCAAAAATATTATCCTCAGCAGTCAACCGGGTACGGACGATCGCGTAACGTGGGTGAAGTCGGTGGATGAAGCCATCGCGGCGGCGGGTGACGTACCAGAAATCATGGTGATTGGCGGCGGTCGCGTTTATGAACAGTTCTTGCCAAAAGCGCAAAAACTGTATCTGACGCATATCGACGCAGAAGTGGAAGGCGACACCCATTTCCCGGATTACGAGCCGGATGAC**GTG**GAATCGGTATTCAGCGAATTCCACGATGCTGATGCGCAGAACTCTCACAGCTATAGCTTTGAGATTCTGGAGCGGCGGAGCGGGTCTGGTGTGCCGGGCCAGCAAAACGCAACATGGATTGTACCCCCGGGACAGTACTTTATGATGGGAGACTGGATGAGTTCCTTTAGCACCCCGGTTTGGATTAGCCAGGCGCAGGGCATTCGTGCCGGTCCGGGCAGCGGTTCTGGGTCCGGCTCGGGCAGTATGGCCACAACCATGACGGCCCTGACAGAAGGTGCGAAGCTGTTCGAGAAGGAGATTCCCTATATCACAGAATTGGAGGGGGATGTAGAGGGTATGAAGTTTATCATCAAAGGCGAAGGGACAGGGGATGCAACAACTGGAACAATTAAGGCTAAGTACATTTGCACGACCGGCGACGTCCCGGTGCCCTGGTCCACGCTCGTCACCACGCTCACGTACGGAGCCCAGTGCTTTGCCAAATATGGCCCTGAACTTAAAGACTTCTACAAGTCGTGTATGCCGGAGGGATACGTGCAAGAGAGGACGATCACCTTTGAAGGTGACGGAGTATTCAAAACAAGAGCGGAGGTGACGTTCGAGAATGGATCGGTCTATAACCGGGTCAAGCTCAACGGACAGGGCTTTAAGAAAGATGGACACGTCCTTGGGAAGAATTTGGAGTTCAATTTCACCCCGCATTGTCTTTACATCTGGGGTGATCAGGCGAATCACGGGTTGAAATCAGCGTTCAAGATCATGCACGAGATTACGGGGAGCAAAGAGGACTTTATCGTGGCAGACCACACTCAGATGAACACTCCAATCGGAGGGGGTCCCGTACACGTACCCGAGTATCATCACCTGACCGTCTGGACATCGTTTGGAAAAGACCCTGACGACGATGAAACTGATCATCTCAACATTGTGGAAGTGATCAAGGCGGTGGACTTGGAAACATACCGGTGA

mKate

ATGGTAAGTGAGCTGATAAAGGAAAACATGCACATGAAATTGTACATGGAAGGTACGGTTAATAATCATCACTTCAAGTGTACGTCCGAAGGGGAGGGAAAACCTTACGAAGGCACACAGACCATGCGGATAAAAGCAGTGGAGGGCGGCCCTCTGCCTTTCGCGTTTGACATACTGGCAACGTCTTTTATGTACGGTAGCAAAACATTCATCAATCACACTCAAGGGATCCCCGATTTTTTCAAGCAATCCTTCCCTGAGGGCTTCACTTGGGAGAGAGTAACTACTTATGAGGATGGGGGGGTTCTGACCGCGACGCAAGATACGTCCTTGCAAGACGGGTGCTTGATATACAACGTTAAAATACGGGGGGTAAACTTCCCAAGTAACGGTCCCGTGATGCAGAAAAAGACTCTCGGGTGGGAGGCTAGTACGGAGACCTTGTATCCGGCAGACGGCGGCCTGGAAGGCCGGGCTGATATGGCCCTTAAACTCGTTGGTGGCGGACATCTCATTTGTAACCTTAAGACTACTTATCGGAGCAAAAAGCCGGCTAAAAATCTGAAAATGCCAGGGGTCTATTACGTTGACCGGCGGCTCGAGCGAATCAAAGAAGCCGACAAAGAGACTTATGTTGAACAACATGAGGTGGCGGTCGCGCGGTATTGCGATCTTCCAAGCAAGCTCGGACACAGGTGA
